# Ether glycerophospholipid control of peroxisome degradation confers protection from ischemia

**DOI:** 10.64898/2026.09.28.755133

**Authors:** Madhu Singh, Shariq Qayyum, Giovanny Godoy, Jenane Benhalima, Olivia Seidel, Nina Madl, Katrin Watschinger, Amy Deik, Hasmik Keshishian, Clary B. Clish, Steven A. Carr, Gregory A. Wyant

## Abstract

Ether glycerophospholipids (ether lipids) are a subclass of glycerophospholipids that are synthesized at the peroxisome and constitute nearly 20% of the total phospholipid content in metazoans. Biochemically, ether lipids have an alkyl chain attached by an ether bond at the sn-1 position of the glycerol backbone (alkyl-ether) primarily containing choline or ethanolamine head groups. Some ether lipids contain a cis bond adjacent to the ether linkage (alkenyl-ether) and are commonly referred to as plasmalogens. While the importance of ether lipids in humans is clear, highlighted by the severity of inherited peroxisomal disorders caused by ether lipid deficiency, the molecular roles of these lipids remain poorly understood. Here, we identify a link between ether lipid metabolism, peroxisome degradation, and oxygen availability. We find Fatty acyl-CoA-reductase 1 (FAR1), the rate-limiting enzyme in peroxisomal ether lipid metabolism, acts as a peroxisome receptor necessary for peroxisome degradation when oxygen is limiting. Hypoxia increases choline ether lipid abundance, promotes autophagy activation, and peroxisome degradation. Mechanistically, FAR1 binds ATG8-family protein LC3B and is necessary and sufficient to control peroxisome degradation under hypoxia. FAR1’s enzymatic function as well as its capacity to bind LC3B is necessary to degrade peroxisomes under hypoxia. Similarly, loss of multiple components of peroxisomal ether lipid metabolism blocks peroxisome degradation under hypoxia in cells and in vivo. In mouse models of chronic HIF activation, Far1 is a direct HIF target gene and is necessary for peroxisome degradation in this setting. Increasing ether lipids using ether lipid precursors activates autophagy and promotes autophagy-dependent cardioprotection against ischemic injury. This work reveals FAR1 as a critical regulator of peroxisome homeostasis in the setting of oxygen limitation.

## Introduction

Peroxisomes are single membrane metabolic organelles that are found in all eukaryotic cells but are often regarded as a redundant component to the mitochondria. The importance of peroxisomes to human biology is clear as peroxisome deficiency, such as in a class of metabolic disorders known as peroxisome biogenesis disorders (PBD), is associated with severe developmental defects and early infantile death^1–4^. While peroxisomes serve diverse cellular roles and are commonly deregulated in human disease, the mechanisms that control peroxisome homeostasis remain incompletely understood^5–9^.

Peroxisome abundance is controlled through the combined action of peroxisomal biogenesis and removal^10,11^. Unlike other organelles, peroxisomal abundance is highly regulated with peroxisomal number increasing or decreasing depending on the cellular need^12,13^. While peroxisome proliferation occurs by the growth and division of pre-existing peroxisomes, peroxisome degradation occurs via their selective removal through autophagy^14–18^. Autophagy is a tightly controlled process in which an autophagosome engulfs excess or damaged cellular components that then fuse with the lysosome for their degradation^19^. Several autophagy-related (*ATG*) proteins coordinate autophagosome formation, maturation, and cargo recognition, including the Serine/threonine-protein kinase ULK1-RB1-inducible coiled-coil protein 1/FAK family kinase interacting protein of 200 kDa (FIP200) complex, the Phosphatidylinositol 3-kinase catalytic subunit type 3/Vacuolar protein sorting 34 (VPS34), and ATG8 lipidation machinery, including ATG7 and ATG5^19^.

While autophagy was initially considered a nonspecific process, it is now appreciated selective forms of autophagy exist for numerous cytoplasmic components, such as the endoplasmic reticulum (ER-phagy), mitochondria (mitophagy), and peroxisomes (pexophagy), to maintain their quality or abundance^20,21^. Autophagy cargo that is selectively degraded is directly or indirectly recognized by the ATG8 family proteins, such as Microtubule-associated protein 1 light chain 3 (LC3) and Gamma-aminobutyric acid receptor-associated protein (GABARAP), that are conjugated to the forming autophagosome via the ATG7 machinery. LC3 or GABARAP bind a selective receptor associated with autophagy cargo via a LC3-interacting region (LIR) consisting of [W/F/Y]_0_-X-X-[L/V/I]_3_ where X is any amino acid and conserved positions X_0_ and X_3_ are aromatic and hydrophobic residues, respectively^22^.

Peroxisomes are essential for cellular lipid, fatty acid, and redox metabolism and their function is intimately dependent on molecular oxygen^23–25^. This has led to the hypothesis that low oxygen may control peroxisome homeostasis^26^. Hypoxia-inducible factor (HIF), which consists of a labile HIFα subunit and stable HIFβ (ARNT) subunit, is the master transcription factor that controls genes whose products promote cellular adaptation to low oxygen. The HIFα subunit is regulated through prolyl hydroxylation by α-ketoglutarate (αKG) dependent dioxygenases known as EglNs (also called Prolyl hydroxylase domain, PHDs)^27–29^. When oxygen is present, HIFα is hydroxylated by the EglN enzymes leading to its polyubiquitination and degradation by an E3 ubiquitin ligase complex containing the von Hippel-Lindau protein (pVHL)^28^. PHD activity requires oxygen, reduced iron, α-ketoglutarate, and can sense other inputs that indirectly reflect oxygen availability, such as reactive oxygen species (ROS) and Krebs cycle metabolites, such as succinate and fumarate^30^.

HIF controls hypoxia-induced mitophagy via the expression of Cbl-2/adenovirus E1B 19-kDa interacting protein E (BNIP3)^31–33^. We and others showed that HIF controls peroxisome degradation and that this is a key component to maladaptive remodeling that occurs in the heart under chronic ischemia^26,34,35^. How, mechanistically, HIF controls peroxisomes degradation remains unknown. Here, using quantitative mass spectrometry-based proteomics analysis of human cells grown under hypoxia, we identified Fatty acyl-CoA reductase 1 (FAR1), the rate-limiting step in ether glycerophospholipid (ether lipid) synthesis, as a candidate peroxisome receptor linked to the degradation of peroxisomes under hypoxia. We validate FAR1 is a peroxisome localized membrane protein and show that autophagy is necessary and sufficient to control its abundance in cells and *in vivo*. FAR1 binds LC3B in vivo via a LIR motif found in its soluble N-terminal domain and Far1 is necessary and sufficient to control peroxisome degradation under hypoxia. As HIF is necessary and sufficient to promote peroxisome degradation, we find FAR1 is a direct HIF target gene and FAR1 is necessary to degrade peroxisomes in the setting of chronic HIF activation. Mechanistically, the capacity for FAR1 to bind LC3B as well as its capacity to reduce fatty acyl-CoAs to fatty alcohols are necessary steps to degrade peroxisomes under hypoxia. As ether lipids have been previously shown to protect tissues against injury, we find increasing ether lipids using ether lipid precursors protects the heart from ischemia via autophagy activation. These findings provide a mechanistic basis by which peroxisomes are degraded under hypoxia.

## Results

### Analysis of the hypoxia-induced pexophagy

We and others have previously showed that hypoxia and HIF activation is necessary and sufficient to promote peroxisome degradation^26,34,35^. Our previous experiments in cardiomyocytes did not test whether peroxisomes are degraded under hypoxia via autophagy in a selective or bulk manner^15^.To address this, we utilized our peroxisome Keima assay reported previously (Figure S1A)^35^. The Keima system is widely used to quantitatively measure the lysosome delivery of autophagy substrates as it provides two important advantages. The Keima protein (25 kDa) is resistant to lysosomal proteases, such that when a Keima fusion protein enters the lysosome, it liberates a “processed” Keima (Keima and any peptidic remnants remaining after proteolysis of its fusion partner), that can be detected via immunoblot analysis. Keima is also pH-responsive and undergoes a chromophore resting charge state change upon trafficking to the lysosome (pH 4.5-5) that can be quantified via the ratio of 561 nm/488 nm excitation by flow cytometry or confocal microscopy (Figure S1A). We utilized two Keima peroxisome reporters (Keima-PEX), one in which we fused Keima to the N-terminus of full-length Peroxisomal membrane protein 11 (Keima-PEX11) and another in which we fused Keima to the N terminus of amino acids 237–305 of human PEX26, which is sufficient for localization to and insertion into the peroxisomal membrane (Keima-PEX26). We designed two different Keima reporters to ensure our results were not confined to any specific peroxisomal protein or to a minimal peroxisomal targeting motif.

We confirmed our Keima-PEX11 fusion localized to ATP-binding cassette sub-family D member 3 (ABCD3/PMP70)-positive peroxisomes in immunofluorescence assays (Figure S1B). As the Keima antibody does not work in fixed samples for immunofluorescence, we engineered an in-frame FLAG epitope tag upstream of the Keima sequence to allow for localization studies with an anti-FLAG antibody to detect Keima-PEX11 localization with endogenous peroxisomal proteins (Figure S1B). We validated our previous results using human U2OS cells stably infected to express our Keima-PEX11 reporter and confirmed autophagy activation via hypoxia promotes lysosomal delivery of the processed Keima (Figure S1C-D)^35^. Similarly, using our Keima-PEX26 system, autophagy activation via hypoxia, EglN inhibition using FG-4497 or mTOR inhibition using Torin1 promoted Keima-PEX processing by immunoblotting. At baseline, a fraction of “processed” Keima was observed at 25 kDa and this was reduced upon SAR405 treatment, a VPS34 inhibitor that blocks the autophagy system, consistent with basal turnover of peroxisomes via autophagy, but not by the E1 inhibitor TAK-243, or proteasomal inhibition using MG132 (Figure S1D). E1 inhibition activated HIF in the presence of oxygen due to inactivation of the pVHL E3 ligase and promoted Keima-PEX processing at baseline (Figure S1D). A faster migrating (∼22 kDa) Keima reactive band is detected in our Keima-PEX26 reporter at baseline that does not report on autophagy-mediated delivery of our Keima-PEX system as it is not blocked by VPS34 inhibition.

To test whether peroxisomes are selectively degraded in cardiomyocytes under hypoxia, we inactivated *ATG7* using Crispr-Cas9 (sgATG7) in human AC16 cardiomyocytes that stably express Keima-PEX. Hypoxia promoted Keima-PEX processing in wild-type AC16 cells, and this effect was completely blocked in sgATG7 cells (Figure S2A). To corroborate these results by measuring the abundance of endogenous peroxisomal proteins, we measured peroxisomal membrane protein PEX14, whose abundance has been previously shown to be controlled via autophagy, as well as peroxisomal luminal enzyme dihydroxyacetone phosphate acyltransferase (GNPAT) abundance in human AC16 cardiomyocytes grown under hypoxia^36,37^. In a time-dependent manner, hypoxia decreased PEX14 and GNPAT protein abundance (Figure S2B). Cells in which we inactivated *VHL* using Crispr-Cas9 (sgVHL) reduced PEX14 and GNPAT protein abundance at baseline. These reduced levels were not further decreased when cells were grown under hypoxia consistent with our prior results in mouse hearts (Figure S2B). ATG7 loss inhibited the reduction in PEX14 and GNPAT abundance when cells were grown under hypoxia (Figure S2B). Consistent with prior results in which HIF is required to mediate peroxisome degradation under hypoxia, ARNT loss, which inactivates HIF-mediated transcription, blocked peroxisome degradation under hypoxia as measured by alkylglycerone phosphate synthase (AGPS) and GNPAT abundance (Figure S2C).

To confirm peroxisomes are degraded in an ATG7-dependent manner *in vivo*, we generated autophagy-deficient mice in which we inactivated the *Atg7* gene in the mouse heart. *Atg7*^+/+^ and *Atg7^fl^*^/fl^ mice were crossed with mice expressing Cre recombinase in cardiomyocytes under the control of the myosin heavy chain promoter (aMHC-Cre). Cardiac *Atg7* loss increased the abundance of a known autophagy receptor, sequestosome 1 (SQSTM1/p62), in mouse hearts as determined by immunoblot analysis (Figure S4D). Like SQSTM1/p62, the abundance of multiple peroxisomal localized proteins, such as Catalase, peroxisomal membrane protein 2 (Pxmp2), Peroxisomal biogenesis factor 16 (Pex16), and Pmp70 increased in *Atg7^fl^*^/fl^-aMHC-Cre mouse hearts when compared to hearts isolated from *Atg7*^+/+^-aMHC-Cre mice (Figure S2D).

Next to BRCA1 gene 1 (NBR1) binds ATG8 family proteins and is a known receptor for pexophagy and ubiquitinated protein aggregates in human cells^38,39^. While HIF does not control NBR1 mRNA abundance, hypoxia-induced peroxisome degradation may require NBR1^26,35^. To test this, we inactivated NBR1 using CRISPR-Cas9 in AC16 cardiomyocytes expressing our Keima-PEX. At baseline, NBR1 abundance was undetectable (Figure S3A). Inhibition of lysosomal-dependent degradation of autophagy cargo via Bafilomycin A1 increased NBR1 abundance consistent with its lysosomal-dependent degradation (Figure S3A). Hypoxia or autophagy activation via Torin1 promoted Keima-Pex processing in wild-type cells and this was reduced under either condition but not completely lost following NBR1 inhibition (Figure S3B).

Hypoxia selectively degrades mitochondria (mitophagy). Hypoxia-induced mitophagy requires the mitochondrial outer-membrane protein FUN14 Domain Containing 1 (FUNDC1) and BNIP3^40,41^. Given that peroxisomes function in concert with mitochondria in several biochemical pathways, FUNDC1 or BNIP3 may also play a role in hypoxia-induced pexophagy^42–45^. To test this, we inactivated FUNDC1 and BNIP3 using CRISPR-Cas9 in AC16 cardiomyocytes (sgBNIP3 and sgFUNDC1) (Figure S3C). Hypoxia activated HIF as measured by increased HIF2α abundance and reduced peroxisomal GNPAT abundance in wild-type AC16 cardiomyocytes and this was blocked in cells lacking ATG7 (Figure S3C). BNIP3 or FUNDC1 loss did not affect the reduction in GNPAT in cells grown under hypoxia (Figure S3C). Consistent with prior studies, BNIP3 loss increased FUNDC1 abundance suggesting when BNIP3-mediated mitophagy is inhibited, the cell may compensate with increasing FUNDC1-mediated mitophagy (Figure S3C)^46^.

### Quantitative analysis of the autophagy controlled cellular proteome under hypoxia identifies FAR1

The observation that ATG7 loss, but not of NBR1, FUNDC1, or BNIP3, blocks hypoxia-induced peroxisome degradation, suggested that an unidentified peroxisome receptor(s) is required to mediate pexophagy under hypoxia. To address this, we examined proteome abundance changes in human U2OS cells. Wild-type and sgATG7 U2OS cells were quantitatively profiled by liquid chromatography, tandem mass spectrometry (LC-MS/MS) using chemical labeling of peptides with tandem mass tag (TMT) for precise, relative quantification to look for proteins that decrease in abundance under hypoxia and require an intact ATG7 pathway for their degradation (Figure 1A). As a key feature of selective autophagy receptors is their abundance is regulated in an ATG7-dependent manner, we predicted that the abundance of an unidentified peroxisome receptor(s) would no longer decrease under hypoxia in cells lacking ATG7. Looking for such ATG7-dependent protein changes was used to identify other organelle-specific receptors such as Testis-expressed protein 264 (TEX264) and Yip1 domain family member 3 and 4 (YIPF3/4)^47,48^. Wild-type and sgATG7 U2OS cell cultures were grown in triplicate under 1% O_2_ (hypoxia) to induce autophagy or 21% O_2_ treated with DMSO (normoxia). Eighteen hours later, total cell extracts were processed for TMT and analyzed by Exploris Mass Spectrometry (Figure 1A). As a positive control for autophagy activation, wild-type U2OS were also treated in triplicate with Torin1 for 18 hours (Figure 1A). Immunoblot analysis of wild-type U2OS cell extracts demonstrated increased HIF1α and decreased SQSTM1/p62 abundance, a marker of autophagy induction, in cells grown under hypoxia as well as suppressed S6Kinase Threonine 389 phosphorylation (pS6K T389), a measure of mTOR activity (Figure S6A). mTOR inhibition using Torin1 treatment completely suppressed pS6K phosphorylation and decreased SQSTM1/p62 abundance (Figure S4A). In cells lacking ATG7, while hypoxia increased HIF1α and suppressed pS6K T389 abundance and Torin1 treatment decreased pS6K T389 abundance, SQSTM1/p62 abundance was no longer reduced under all conditions, consistent with ATG7-dependent control of its abundance (Figure S4A). Following successful quality control of cell extracts, U2OS cell extracts were processed for quantitative proteome profiling (Figure 1A and methods). Proteomic analysis of U2OS cell extracts quantified 155,419 peptides and 10,315 proteins across all conditions (Figure S4B). Principal component analysis distinguished triplicate cell extracts for each condition based on treatment group (Figure S4C). Analysis of the wild-type and sgATG7 proteome revealed an increase in the abundance of many established autophagy cargo receptors, such as Nuclear receptor coactivator 4 (NCOA4), NBR1, SQSTM1/p62, Calcium binding and coiled-coil domain 2 (CALCOCO2), Calcium binding and coiled-coil domain 1 (CALCOCO1), Tax1 binding protein 1 (TAX1BP1), YIPF3, and YIPF4, as well as established autophagy associated proteins, such as GABARAPL2, in cells lacking ATG7 (Figure 1B)^20,21,47,49^. Hypoxia and Torin1 treatment decreased the abundance of many peroxisomal proteins consistent with peroxisome degradation and activated autophagy in wild-type cells as measured by decreased abundance of known autophagy cargo receptors and associated proteins, such as TEX264, CALCOCO2, YIPF3, FUNDC1, SQSTM1/p62, NBR1, Cell Cycle Progression Gene 1 (CCPG1), Immunity-related GTPase family Q (IRGQ), and YIPF4, in an ATG7-dependent manner (Figure 1C-E). Hypoxia increased HIF1α abundance as well as the protein products of downstream HIF target genes, such as BNIP3 and DEPP1, in wild-type and sgATG7 cells (Figure 1D-E).

**Figure 1.**
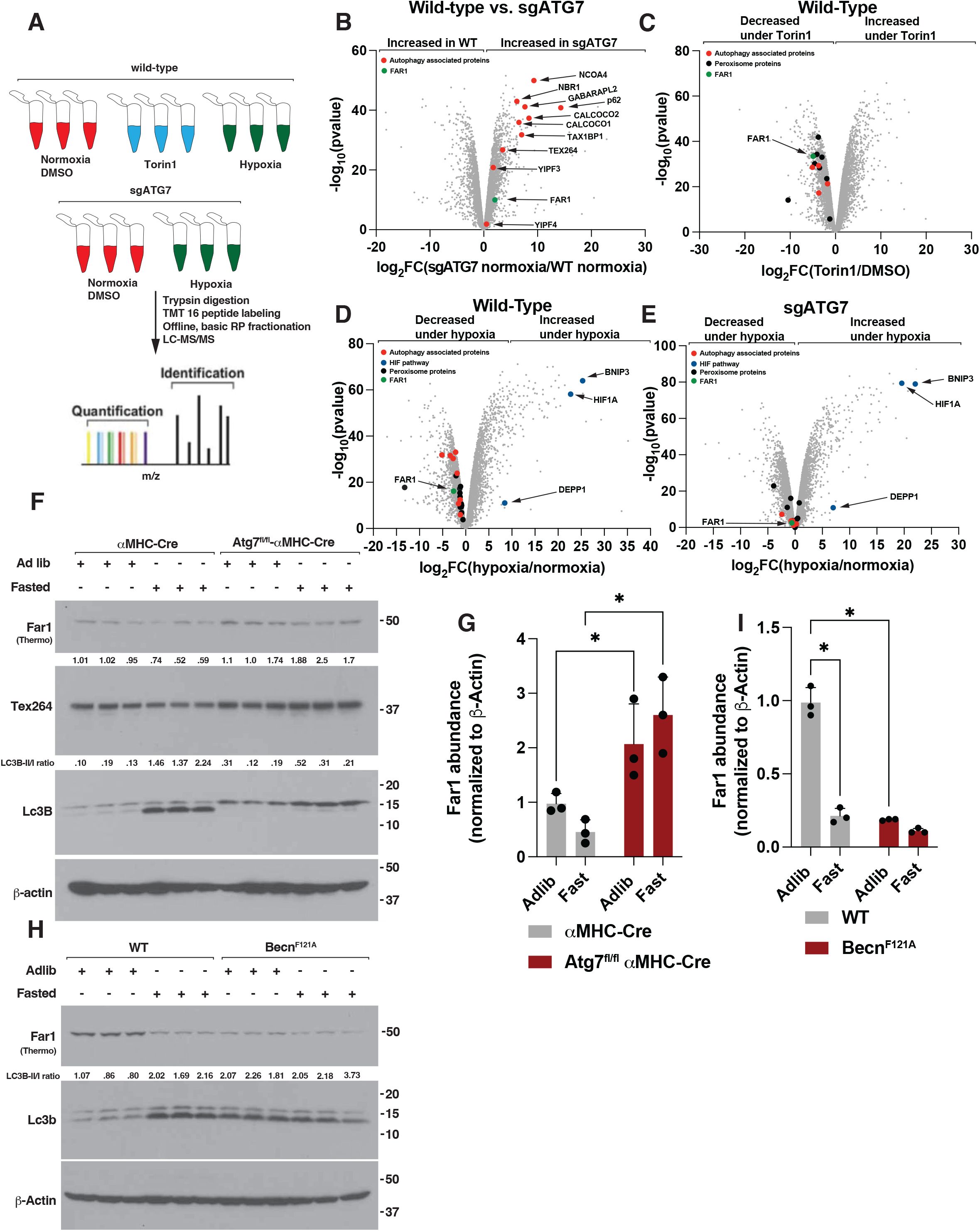
Quantitative analysis of the autophagy controlled cellular proteome under hypoxia identifies FAR1. **(A)** Schematic of proteomics workflow used for analysis of autophagy controlled cellular proteome under hypoxia. **(B-E)** Volcano plots comparing wild-type versus sgATG7 U2OS cells (**B**), wild-type cells treated with Torin1 or DMSO (**C**), hypoxia (0.1% O2) versus normoxia in wild-type cells (**D**), and sgATG7 cells (**E**). Highlighted in red are known autophagy-related proteins, FAR1 is shown in green, HIF-related proteins in blue, and peroxisome-localized proteins in black. **(F)** Immunoblot analysis of Atg7^+/+^-αMHC-Cre and Atg7^fl/fl^-αMHC-Cre mouse hearts. Where indicated, mice were fasted for 48 hours or fed ad libitum prior to euthanasia and tissue harvesting. **(G)** Densitometry analysis of Far1 abundance shown in (F). N=3, *indicates P<0.05. **(H)** Immunoblot analysis of mouse hearts derived from wild-type and Beclin^F121A^ mice. Where indicated, mice were fasted for 48 hours or fed ad libitum prior to euthanasia and tissue harvesting. **(I)** Densitometry analysis of Far1 abundance shown in (H). N=3, *indicates P<0.05. Quantification of protein abundance is shown above individual immunoblots. Far1 immunoblots indicate which Far1 antibody was used to detect Far1.

To narrow our search for candidate peroxisome receptors, we prioritized proteins whose abundance decreased in an ATG7-dependent manner under autophagy activation (Torin1 and hypoxia). As many organelle-specific autophagy receptors contain a transmembrane domain for stable association with its organelle as well as a soluble portion with the capacity to bind the autophagy machinery, we looked for peroxisome embedded proteins that also have the capacity to bind the cytosolic autophagy machinery via a cytoplasmic facing domain^40,47–50^. Among the proteins whose abundance was reduced by hypoxia and Torin1 treatment in an ATG7-dependent manner was the peroxisomal protein Fatty acyl-CoA reductase 1 (FAR1), the rate-limiting enzyme necessary for peroxisomal ether lipid synthesis (Figure 1B-E)^51^. Ether lipids are a specialized class of glycerophospholipids comprised of alkyl-ether and alkenyl-ether (plasmalogens) lipids of which the latter are distinguished from alkyl-ether lipids via the addition of a vinyl ether bond at the *sn*-1 position via the ER localized plasmanylethanolamine desaturase 1 (PEDS1, former gene symbol *TMEM189*)^52–54^. FAR1 abundance was increased at baseline in cells lacking ATG7 when compared to wild-type cells in our proteomics dataset (Figure 1B). FAR1 was of interest as it has not been associated with autophagy before, it contains a single c-terminal transmembrane segment that is necessary for peroxisomal localization and a large N-terminal domain that faces into the cytosol, and we noticed that FAR1 bound ATG8-familiy protein LC3B in a published dataset of anti-FLAG-LC3B affinity proteomics in 293T cells^50,55^.

To test whether FAR1 abundance is controlled via autophagy, we generated mice in which we inactivated the *Atg7* gene in the mouse heart or skeletal muscle (Figure 1F-G and S4D-F). *Atg7*^+/+^ and *Atg7^fl^*^/fl^ mice were crossed with mice expressing Cre recombinase in cardiomyocytes under the control of the myosin heavy chain promoter (aMHC-Cre) or in skeletal muscle cells under the control of the human alpha-skeletal actin promoter (Acta-Cre). Cardiac or skeletal muscle *Atg7* loss increased the abundance of known autophagy receptors like Nbr1 and Tex264 in mouse hearts and quadricep muscles as determined by immunoblot analysis (Figure 1F and S4D-E)^38^. Like Nbr1 and Tex264, Far1 abundance increased in mouse hearts and quadriceps muscles derived from *Atg7^fl^*^/fl^-aMHC-Cre and *Atg7^fl^*^/fl^-Acta-Cre mice when compared to hearts and quadriceps muscles isolated from *Atg7*^+/+^-aMHC-Cre and *Atg7*^+/+^-Acta-Cre mice (Figure 1F-G and Figure S4D-F). Autophagy activation via fasting decreased Far1 abundance in mouse hearts derived from *Atg7*^+/+^-αMHC-Cre when compared to mouse hearts isolated from ad libitum fed *Atg7*^+/+^-αMHC-Cre mice and this decrease was lost in hearts derived from *Atg7^fl^*^/fl^-αMHC-Cre mice (Figure 1F-G). The apparent increase in Far1 abundance upon Atg7 loss was not due to an increase in *Far1* mRNA abundance (Figure S4G).

To determine whether autophagy activation is sufficient to reduce Far1 abundance in vivo, we compared Far1 abundance in mouse hearts derived from wild-type or Beclin1^F121A/F121A^ knock-in mice (Becn^F121A^), which express a Phe121Ala mutation that decreases its interaction with its negative regulator Bcl2 leading to higher levels of basal autophagic flux^56^. We fasted wild-type and Becn^F121A^ mice for 48 hours (to induce autophagy) or provided them with food ad libitum prior to euthanasia and tissue analysis. Fasting activated autophagy, as measured by increased Lc3b lipidation, when compared to hearts derived from ad libitium fed mice in immunoblot assays (Figure 1H)^57^. Becn^F121A^ mice exhibit enhanced autophagy at baseline as measured by increased Lc3b lipidation in immunoblot assays in mouse heart lysates when compared to wild-type littermates (Figure 1H). Fasting decreased Far1 abundance in mouse hearts derived from fasted wild-type mice when compared to mouse hearts isolated from ad libitum fed wild-type mice (Figure 1H-I). Far1 was constitutively low in ad libitum fed Becn^F121A^ mouse heart lysates when compared to wild-type mice and this was not further decreased in Becn^F121A^ mouse hearts isolated from fasted Becn^F121A^ mice (Figure 1H-I). Collectively, these data suggest Far1 abundance is controlled via autophagy.

### FAR1 is a peroxisome localized membrane protein that is trafficked to the lysosome and degraded under autophagy activation

Using an antibody raised to detect endogenous FAR1 in cells, we validated FAR1 localizes to PMP70-positive and Peroxisomal biogenesis factor 19 (PEX19)-positive peroxisomes in cardiomyocytes using immunofluorescence (Figure 2A and S5A). Additionally, we utilized the PEROXO-IP method to biochemically measure FAR1 abundance in intact peroxisomes (Figure 2B)^58^. We captured peroxisomes via anti-hemagglutinin (HA) immunoprecipitation (IP) from 293FT cells stably expressing amino acids 237-305 of human PEX26, which are sufficient for its proper localization and insertion in the peroxisomal membrane, fused to a 3xhaemagglutinin (HA) and a monomeric enhanced green fluorescent protein (EGFP) (3xHA-eGFP-PEX26; referred to as HA-PEX) or amino acids 237-305 of human PEX26 fused to a 3xMyc and EGFP (3xMyc-eGFP-PEX26; referred to as Control-PEX). Immunoblot analysis of anti-HA immunoprecipitates isolated from Control-PEX and HA-PEX revealed enrichment of peroxisomal proteins, such as GNPAT and PEX14, as well as FAR1 in our HA-PEX IPs relative to a variety of other organelle and cytosolic proteins, such as 130 kDa cis-Golgi matrix protein 1 (GM130, Golgi apparatus), β-actin (cytoplasmic), Prohibitin 1 (PHB1, mitochondria), and ribosomal protein S6 (S6, cytoplasmic) (Figure 2B). Using subcellular fractionation assays, we validated endogenous FAR1 is associated with membranes in human AC16 cardiomyocytes consistent with it residing in the peroxisomal membrane (Figure S5B). In live cells, exogenous FAR1 when fused to mNeonGreen (mNG-FAR1) did not co-localized with mitochondria as marked by MitoTracker (Figure S5C). mNG-FAR1-positive peroxisomes did, however, associate with MitoTracker-positive mitochondria consistent with direct mitochondrial-peroxisome contacts between these organelles (Figure S5C)^59^.

**Figure 2.**
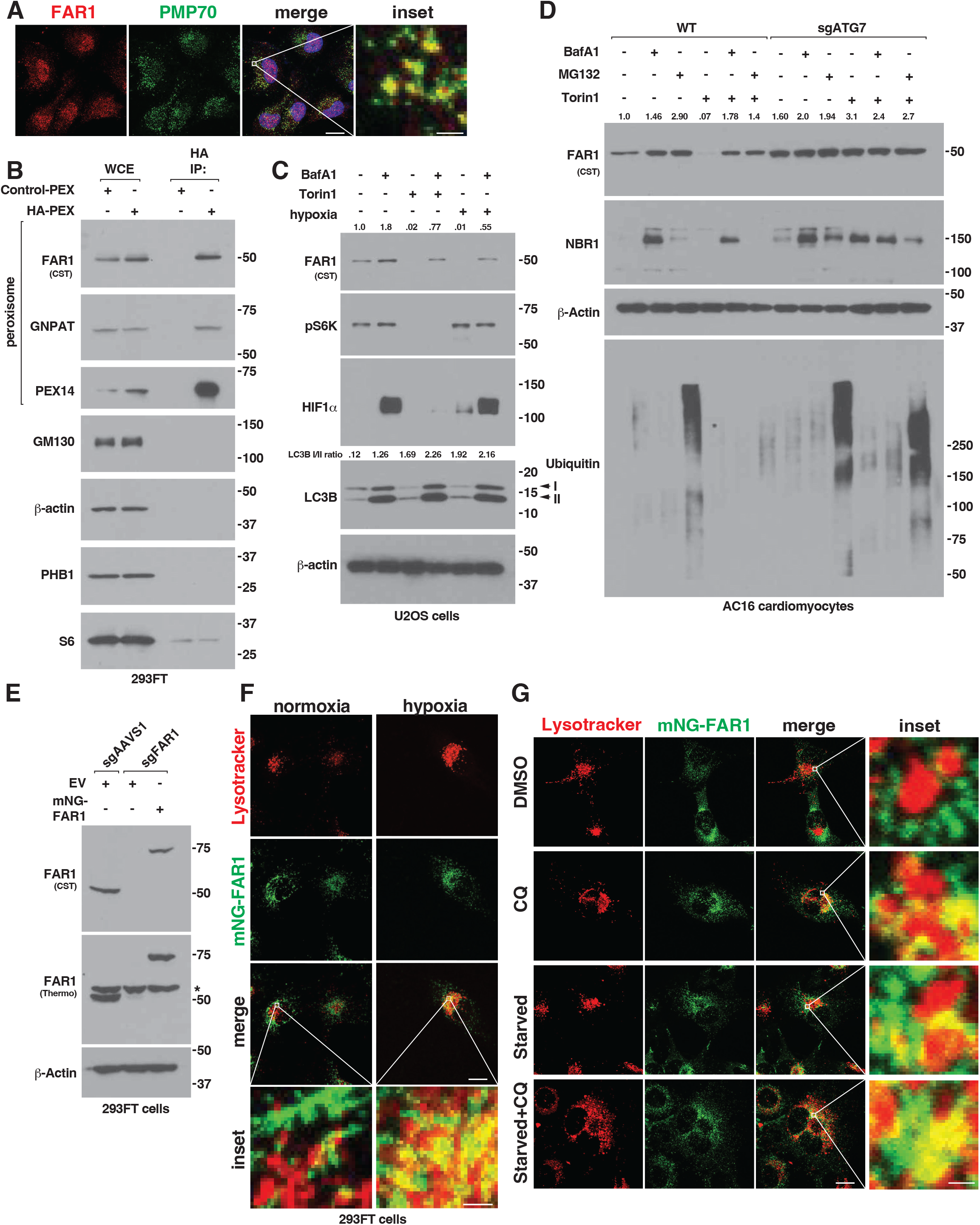
FAR1 is a peroxisome localized membrane protein that is trafficked to the lysosome and degraded under autophagy activation. (**A**) Confocal microscopy analysis of mouse neonatal cardiomyocytes. Scale bar indicates 10 um, inset scale bar indicates 1 um. (**B**) Immunoblot analysis of anti-HA immunoprecipitates isolated from 293FT cells stably expressing Control-PEX and HA-PEX. WCE, whole cell extract. Far1 immunoblot indicate which Far1 antibody was used to detect Far1. (C-D) Immunoblot analysis of human U2OS osteosarcoma cells (**C**) and wild-type and sgATG7 human AC16 cardiomyocytes (**D**). Where indicated, cells were treated with Bafilomycin A1 (BafA1, 500 nM), MG132 (1 uM), Torin1 (250 nM), grown at 0.1% O_2_ (hypoxia), or DMSO and grown at normoxia for 18 hours prior to cell lysis. Quantification of protein abundance is shown above individual immunoblots. Far1 immunoblots indicate which Far1 antibody was used to detect Far1. (**E**) Immunoblot analysis of clonal sgAAVS1 and sgFAR1 293FT cells stably expressing mNeonGreen-FAR1 (mNG-FAR1) or empty vector (EV). *indicates non-specific band. Far1 immunoblots indicate which Far1 antibody was used to detect Far1. (**F-G**) Confocal microscopy analysis of sgFAR1 293FT cells (F) and sgFAR1 human AC16 cells (G) stably expressing mNG-FAR1 and labeled with 100 nM lysotracker (LT). Where indicated, cells were grown under 1% O_2_ (hypoxia), treated with chloroquine (CQ, 30 uM), starved of nutrients (starved), treated with chloroquine and starved of nutrients, or grown at normoxia and treated with DMSO prior to imaging. Cells were nutrient starved and CQ treated for 8 hours while cells were grown under hypoxia for 24 hours. Scale bar indicates 10 um, inset scale bar indicates 1 um.

Upon autophagy activation, autophagy cargo and their receptors are trafficked to the lysosome for their degradation. To test whether FAR1 is degraded via the lysosome when autophagy is activated, we measured FAR1 abundance under hypoxia, Torin1, or nutrient starvation. Indeed, Torin1 treatment, nutrient starvation, or hypoxia decreased FAR1 abundance in human U2OS cells, AC16 cardiomyocytes, 293FT, and mouse C2C12 myoblasts and this reduction was restored in cells treated with the lysosomal inhibitors Bafilomycin A1 (BafA1) or chloroquine (CQ), or in cells lacking ATG7 (Figure 2C-D and S6A-D). Bafilomycin A1 and chloroquine treatment alone increased FAR1 supporting FAR1 is degraded via basal autophagy flux (Figure 2C-D and S6A-D). Proteosome inhibition using MG132 increased FAR1 abundance like Bafilomycin A1 and restored FAR1 abundance following Torin1 treatment (Figure 2D and S6D). The apparent increase in FAR1 abundance in the setting of lysosomal inhibition with chloroquine or proteasome inhibition with MG132 was not due to an increase in *FAR1* mRNA (Figure S6E). In cells lacking ATG7, FAR1 abundance at baseline is increased and is unaffected by lysosomal or proteasomal inhibition (Figure 2D). To measure whether FAR1 is trafficked to the lysosome under autophagy activation, we generated clonal sgFAR1 293FT cells using CRISPR-Cas9 and introduced mNG-FAR1 at endogenous levels by retroviral infection and measured FAR1 co-localization with lysosomes using live cell imaging following hypoxia (Figure 2E-F). We confirmed mNG-FAR1 in sgFAR1 293FT cells localized to PEX14-positive peroxisomes using immunofluorescence (Figure S6F). In cells grown under hypoxia, FAR1 increased its co-localization with lysosomes as marked by LysoTracker when compared to cells grown under normoxia using confocal microscopy and image analysis (Figure 2F and S6G). Consistent with this, inhibition of lysosomal degradation using chloroquine or autophagy activation via nutrient starvation increased the co-localization of mNG-FAR1 with Lysotracker when compared to untreated cells grown in the presence of full nutrients (Figure 2G and S6H). Nutrient starvation in the presence of CQ treatment increased the co-localization of FAR1 with lysosomes when compared to nutrient starvation alone (Figure 2G and S6H).

### FAR1 binds LC3B in multiple tissues and in cells

We hypothesized FAR1 may interact with ATG8-family proteins, such as LC3B, like other known autophagy receptors. To test this in vivo, we utilized transgenic *CAG-RFP-EGFP-LC3* mice in which a CAG promotor sequence drives the expression of red fluorescent protein (RFP) fused to both EGFP and microtubule-associated protein 1 light chain 3 alpha (LC3)^60^. Using anti-RFP affinity beads, we immunoprecipitated Lc3 from mouse hearts and brains isolated from wild-type and *CAG-RFP-EGFP-LC3* mice fed ad libitum, deprived of food for 48 hours (fasted, to activate autophagy), or housed at 10% O_2_ (hypoxia) for 7-days followed by immunoblotting (Figure 3A-B). Anti-RFP immunoprecipitates isolated from mouse hearts and brains derived from *CAG-RFP-EGFP-LC3* mice, but not wild-type mice, bound established autophagy receptors, such as Nbr1 and Yipf3 (Figure 3A-B)^38,39,47,49^. Hypoxia or fasting reduced the amount of Nbr1 and Yipf3 bound to Lc3 presumably due to autophagy-dependent degradation under these conditions (Figure 3A-B). Far1 bound Lc3 in a similar pattern to Nbr1 and Yipf3 in this assay (Figure 3A-B).

**Figure 3.**
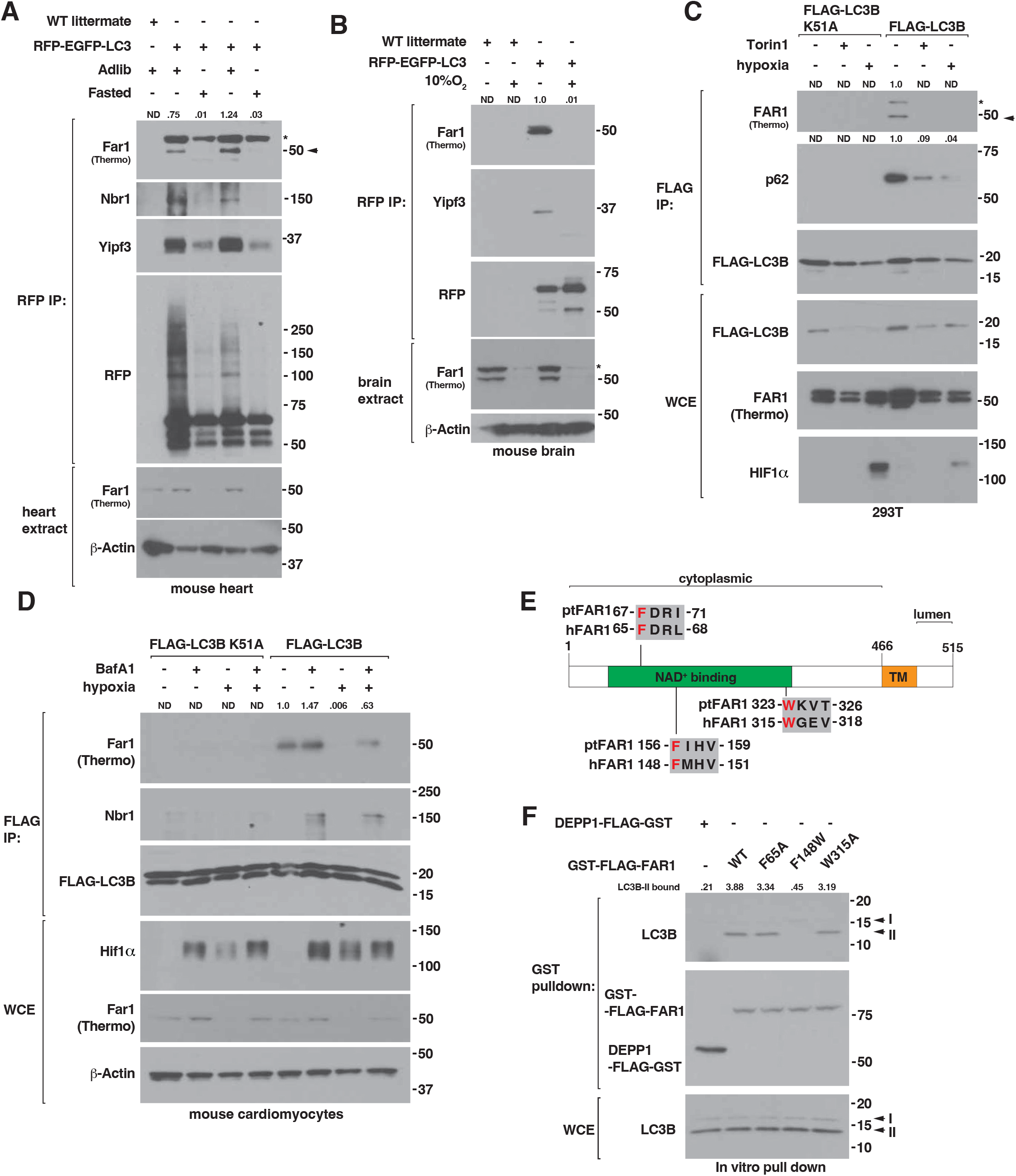
FAR1 binds LC3B in multiple tissues and in cells. (A-B) Immunoblot analysis of anti-RFP immunoprecipitates isolated from mouse heart **(A)** and brain **(B)** lysates prepared from wild-type (WT littermate) and *RFP-EGFP-LC3* mice. Where indicated, mice were ad libitum fed and housed at normal atmospheric oxygen (Ad lib), fasted for 48 hours, or housed at 10% O_2_ for 7-days prior to euthanasia and tissue harvesting. Arrow indicates Far1 while * indicates non-specific band. Quantification of protein abundance is shown above individual immunoblots. Far1 immunoblots indicate which Far1 antibody was used to detect Far1. **(C-D)** Immunoblot analysis of anti-FLAG immunoprecipitates isolated from human 293T cells **(C)** and mouse neonatal cardiomyocytes **(D)** stably expressing FLAG-LC3B K51A or FLAG-LC3B. Where indicated, cells were treated with Torin1 (250 nM), Bafilomycin A1 (BafA1, 500 nM), grown at 0.1% O_2_ (hypoxia), or treated with DMSO and grown at normoxia for 18 hours prior to cell lysis. Arrow indicates FAR1 while * indicates non-specific band. WCE, whole cell extract. Quantification of protein abundance is shown above individual immunoblots. Far1 immunoblots indicate which Far1 antibody was used to detect Far1. **(E)** Schematic of human FAR1 (hFAR1) protein domains and putative LIR sequence conservation compared to Paramecium tetraurelia Far1 (ptFAR1) homolog GSPATT00037955001. **(F)** Immunoblot analysis of in vitro binding reaction of immobilized recombinant Glutathione S-transferase (GST)-FLAG-FAR1, GST-FLAG-FAR1 LIR variants, or DEPP1-FLAG-GST mixed with 293FT cell lysates. 293FT cells were treated with chloroquine (CQ, 30 uM) for 18 hours prior to cell lysis and subsequent mixture with 3 ug recombinant GST-FLAG fusion protein. WCE, whole cell extract.

Selective autophagy receptors have LC3-interacting regions (LIRs) that bind to a LC3B LIR recognition motif (LIRM). Mutation of LC3B lysine 51 (K51) disrupts the LC3B LIRM and inhibits the LC3B-autophagy receptor interaction^61^. As hypoxia promotes peroxisome degradation, we hypothesized that hypoxia controls the FAR1-LC3 interaction, and this interaction is mediated with a LIR motif. To address this, we stably expressed wild-type and K51A LC3B fused to a FLAG epitope in human 293FT cells and mouse neonatal cardiomyocytes and grew these cells under 1% O_2_ (hypoxia), treated them with Torin1 (to activate autophagy), or grew these cells under normoxic conditions treated with DMSO followed by anti-FLAG immunoprecipitation. Anti-FLAG immunoprecipitates isolated from FLAG-LC3B expressing 293FT cells and mouse neonatal cardiomyocytes bound known autophagy receptors, such as SQSTM1/p62, TEX264, NBR1, and FAR1 in a K51-dependent manner (Figure 3C-D and S7A). Lysosomal inhibition with chloroquine increased the amount of SQSTM1/p62, TEX264, NBR1, and FAR1 bound to wild-type LC3B in cardiomyocytes (Figure 3D and S7A). Autophagy activation via hypoxia or mTOR inhibition with Torin1 reduced the amount of SQSTM1/p62, TEX264, and FAR1 bound to wild-type LC3B (Figure 3C-D and S7A). This apparent reduction was due to lysosomal-mediated degradation of SQSTM1/p62, TEX264, and FAR1 as chloroquine treatment under hypoxia restored their abundance bound to LC3B (Figure 3D and S7A). The capacity for FAR1 to bind LC3B did not require its enzymatic role in reducing fatty acids to their respective fatty alcohols as an enzymatically dead FAR1 variant (FAR1 D365G) bound LC3B to similar degree as wild-type FAR1 (Figure S7B)^62^.

To identify a FAR1 point mutant that disrupts its interaction with LC3B, we generated single FAR1 point mutations in residues that correspond to canonical LIR sequences of a consensus motif [W/F/Y]xx[L/I/V] surrounded by at least one proximal acidic residue^22^. We focused on three putative FAR1 LIR regions conserved down to the Paramecium tetraurelia FAR1 homolog GSPATT00037955001 (ptFAR1) that are contained within FAR1’s soluble N-terminal domain (Figure 3E). We performed GST-pulldown analyses of recombinant GST-fusion proteins consisting of wild-type FAR1, FAR1 LIR variants, or a control protein DEPP1 with HEK293FT cell lysates prepared from chloroquine-treated cells to enrich for autophagosome bound lipidated LC3B (LC3-II) that is otherwise degraded in the lysosome. We discovered that mutating FAR1 F148 to tryptophan (FAR1 F148W) reduced its interaction with LC3B when compared to wild-type FAR1 (Figure 3F). FAR1 F148W fused to mNeonGreen (mNG-Far1 F148W) localized to peroxisomes, like wild-type FAR1, as measured by co-localization with mRFP-PEX26 in cardiomyocytes by live cell imaging and confocal microscopy (Figure S8A).

### FAR1 is necessary and sufficient to degrade peroxisomes under hypoxia

To test the role of FAR1 in the degradation of peroxisomes under hypoxia, we stably expressed our Keima-PEX reporter in cells in which we inactivated FAR1 using CRISPR-Cas9 (sgFAR1)^35^. We confirmed FAR1 loss reduced alkyl-ether and alkenyl-ether (plasmalogen) lipid abundance consistent with its role in ether lipid synthesis (Figure S9A-B). Hypoxia promoted Keima-PEX processing in wild-type cardiomyocytes and this effect was lost in cells lacking FAR1 as measured by immunoblot analysis (Figure 4A and S9C-D). Consistent with this, hypoxia promoted FAR1-dependent peroxisomal (Peroxisomal biogenesis factor 14, PEX14) co-localization with LC3B-positive autophagosomes in cardiomyocytes by confocal microscopy (Figure S9E-F). FAR1 loss did not disrupt autophagy activation, as measured by LC3B lipidation, nor the degradation of established autophagy receptors, such as YIPF3, under Torin1 treatment suggesting FAR1 loss does not inhibit the degradation of other selective autophagy cargos, such as the Golgi apparatus, or generally disable the autophagy system (Figure S9G). FAR1 loss reduced HIF stabilization under hypoxia, but this was not due to disruption of the EglN-pVHL axis as cells lacking FAR1 stabilized HIF to a similar degree as wild-type cells when treated with EglN inhibitor FG-4592 or the prolyl hydroxylase inhibitor dimethyloxalylglycine (DMOG) (Figure S9H). DMOG (8 hours treatment) reduced FAR1 abundance that was not reflected in the setting of FG-4592 or hypoxia at this time point (Figure S9H). This difference may be in part due to DMOG-mediated transcriptional repression of peroxisome proliferation like has been shown in other studies^34^.

**Figure 4.**
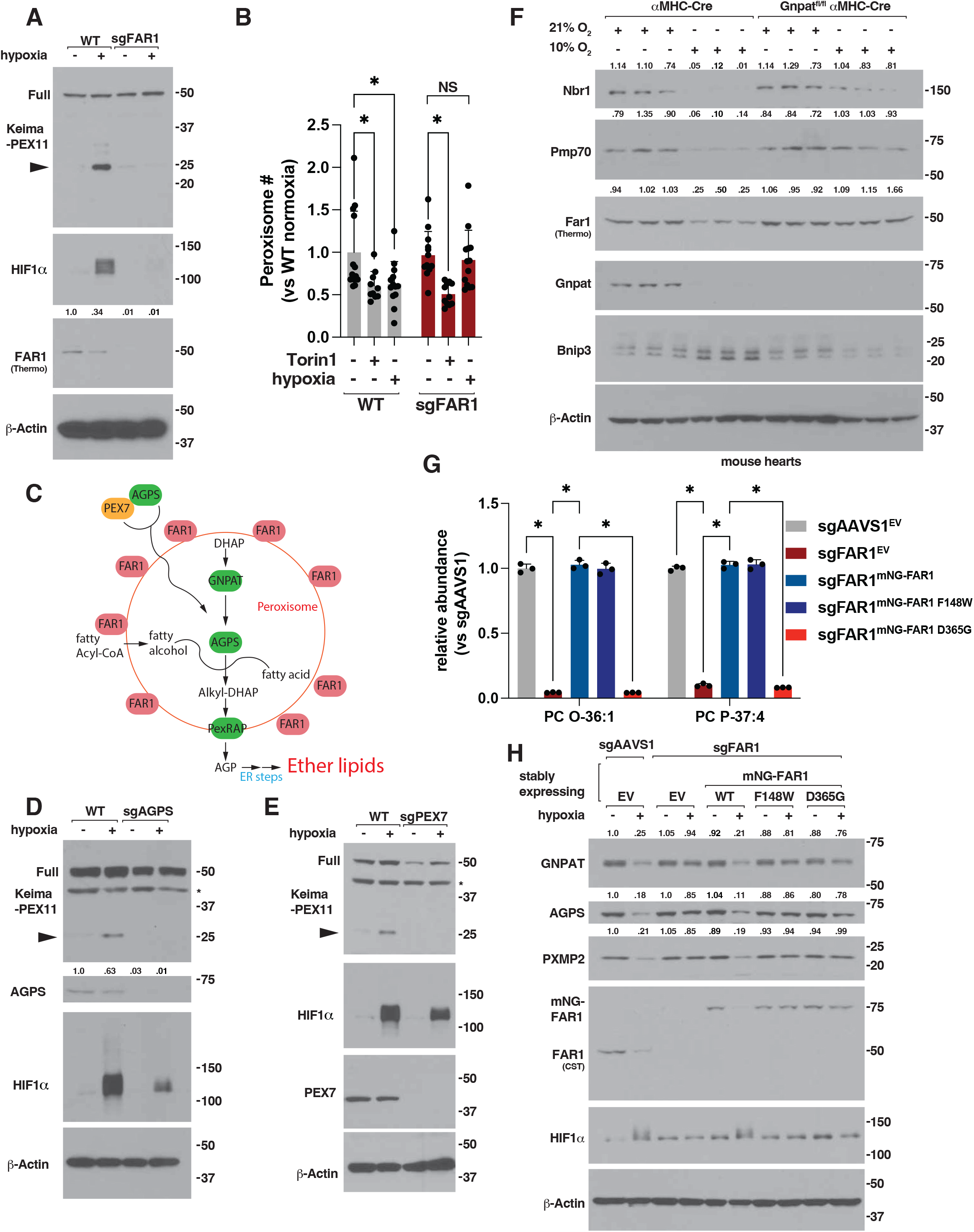
FAR1 and ether lipid metabolism are required for hypoxia-induced peroxisome degradation. **(A)** Immunoblot analysis of wild-type (WT) and sgFAR1 human AC16 cardiomyocytes stably expressing Keima-PEX11. Where indicated, cells were grown under 0.1% O2 (hypoxia) or normoxia for 18 hours prior to cell lysis. Full indicates full-length Keima-fusion protein and arrow indicates liberated Keima protein. Far1 immunoblot indicates which Far1 antibody was used to detect Far1. **(B)** Confocal microscopy analysis and quantification of PEX14-positive peroxisomes in wild-type (WT) and sgFAR1 human AC16 cardiomyocytes. Where indicated, cells were treated with Torin1 (250 nM), grown at 0.1% O_2_ (hypoxia), or treated with DMSO and grown at normoxia for 24 hours prior to fixation. * indicates P<0.05. NS, non-significant. N>10 images per group. **(C)** Schematic of peroxisomal ether lipid metabolism. **(D-E)** Immunoblot analysis of wild-type, **(D)** sgAGPS, or **(E)** sgPEX7 human AC16 cardiomyocytes stably expressing Keima-PEX11. Where indicated, cells were grown under 0.1% O2 (hypoxia) or normoxia for 18 hours prior to cell lysis. Full indicates full-length Keima-fusion protein and arrow indicates liberated Keima protein. * indicates non-specific band. **(F)** Immunoblot analysis of Gnpat^+/+^-αMHC-Cre and Gnpat^fl/fl^-αMHC-Cre mouse hearts. Where indicated, mice were housed at 10% O_2_ (hypoxia) or 21% O_2_ (atmospheric oxygen) for 7 days prior to euthanasia. Quantification of protein abundance is shown above individual immunoblots. Far1 immunoblot indicates which Far1 antibody was used to detect Far1. **(G)** Lipidomic analysis of alkyl and alkenyl PC species in sgAAVS1 and sgFAR1 HEK-293FT cells stably expressing wild-type mNeonGreen-FAR1 or variants (mNG-FAR1) or empty vector (EV). * indicates P<0.05. N=3 per group. **(H)** Immunoblot analysis of clonal sgAAVS1 and sgFAR1 AC16 cells stably expressing wild-type mNeonGreen-FAR1 or variants (mNG-FAR1) or empty vector (EV). Where indicated, cells were grown at 0.1% O_2_ (hypoxia) or normoxia for 36 hours prior to cell lysis. Quantification of protein abundance is shown above individual immunoblots. Far1 immunoblot indicates which Far1 antibody was used to detect Far1.

Several other types of pexophagy have been reported in mammalian cells, like starvation-induced or ubiquitin-mediated, with mutually distinct factors required for each form of pexophagy. Starvation-induced pexophagy is mediated via mTOR inhibition. Hypoxia and mTOR inhibition via Torin1 reduced peroxisome number as measured by immunofluorescence and confocal microscopy (Figure 4B). FAR1 loss maintained peroxisome abundance under hypoxia but did not affect peroxisome loss under Torin1 treatment (Figure 4B). Peroxisomal biogenesis factor 1 (PEX1) loss is sufficient to induce ubiquitin-dependent pexophagy in human cells^63,64^. To test the role of FAR1 in the setting of PEX1 loss, we targeted PEX1 using Crispr-Cas9 in wild-type and sgFAR1 cardiomyocytes. PEX1 loss reduced PEX14-positive peroxisomes as measured by immunofluorescence and this was increased but not fully restored in cells lacking FAR1 (Figure S10A-B).

FAR1 initiates ether lipid synthesis in the peroxisome which is then completed in the ER (Figure 4C)^51^. The requirement of FAR1 for peroxisome degradation under hypoxia may be related to its role in ether lipid metabolism, its capacity to bind LC3B, or both. We analyzed genetic perturbation datasets from the DepMap project to identify genes with correlated essentialities with FAR1 loss to inform our understanding of FAR1 function^65^. As expected, FAR1 co-essentiality mapping revealed numerous components of peroxisomal ether lipid synthesis, such as *AGPS* and *GNPAT*, ER-localized PEDS1 and Selenoprotein I (*EPT1*), as well acyl-CoA synthetase long-chain family 4 (*ACSL4*), an enzyme that supports phospholipid metabolism necessary for ether lipid synthesis (Figure S11A). FAR1 loss in cardiomyocytes increased mRNA abundance of multiple components of the peroxisomal arm of ether lipid synthesis, such as *AGPS*, the FAR1 paralog *FAR2*, and *GNPAT*, but did not affect the expression of ER localized *PEDS1* by quantitative PCR (qPCR) analysis (Figure S11B). Despite an increase in *AGPS* and *GNPAT* mRNA in cells lacking FAR1, we did not detect a reproducible increase in AGPS or GNPAT at the protein level in cells via immunoblot (Figure S11C). Peroxisomal biogenesis factor 7 (PEX7), the peroxisomal targeting signal 2 (PTS2) receptor necessary for peroxisomal localization of a set of peroxisomal matrix enzymes, also scored very highly presumably due to its requirement in proper peroxisomal localization of AGPS (Figure S11A)^66^. Consistent with this, PEX7 co-essentiality mapping revealed *AGPS*, *PEDS1*, *FAR1*, *EPT1*, and *GNPAT* supporting that the PEX7-PTS2 pathway is necessary for cellular ether lipid metabolism (Figure S11D). As these results suggest ether lipid metabolism is an essential function of FAR1, we tested whether inactivation of other members of ether lipid synthesis blocked peroxisome degradation under hypoxia. Indeed, AGPS and PEX7 loss blocked hypoxia induced peroxisome degradation in cardiomyocytes using our Keima-PEX reporter (Figure 4D-E, S11E-F). Inhibition of hypoxia-induced peroxisome degradation in cells lacking an intact ether lipid metabolism pathway was specific as loss of ACSL4, which is necessary for polyunsaturated lipid but not ether lipid metabolism, did not block Keima-PEX processing under hypoxia (Figure S11G). Like FAR1 loss, loss of AGPS and PEX7 but not ACSL4, reduced HIF stabilization under hypoxia but this was not due to disruption of the EglN-pVHL axis as cells lacking AGPS or PEX7 stabilized HIF to a similar degree as wild-type cells when treated with EglN inhibitor FG-4592 (Figure S11H-I). PEX7 loss reduced AGPS abundance as PEX7 is necessary for proper AGPS import into peroxisomes (Figure S11H)^66^.

To test the role of ether lipid metabolism in hypoxia-induced peroxisome degradation *in vivo*, we inactivated *Gnpat* in the mouse heart (Figure 4F). *Gnpat*^+/+^ and *Gnpat^fl^*^/fl^ mice were crossed with αMHC-Cre mice and we housed these mice at 10% O_2_ (hypoxia) or 21% O_2_ (atmospheric oxygen) for 7-days to induce autophagy. Hypoxia decreased the protein abundance of Gnpat as well as other peroxisome localized proteins like Pmp70, Nbr1, and Far1 in mouse hearts isolated from *Gnpat*^+/+^ αMHC-Cre (Figure 4F). Cardiac *Gnpat* loss reduced the loss of Pmp70, Nbr1, and Far1 abundance under hypoxia consistent with the key role of ether lipid metabolism in peroxisome degradation under hypoxia (Figure 4F). Hypoxia activated HIF as measured by increased Bnip3 abundance, a classic HIF target gene, in hearts derived from *Gnpat*^+/+^ αMHC-Cre mice and this required *Gnpat*, consistent with our results in cells in which inactivation of ether lipid metabolism inhibits HIF under hypoxia (Figure 4F).

To dissect the contribution of FAR1’s enzymatic role in ether lipid metabolism versus its capacity to bind LC3B to its role in peroxisome degradation under hypoxia, we stably expressed wild-type FAR1, our LC3B-binding deficient FAR1 variant (FAR1 F148W), a enzymatically dead FAR1 variant (FAR1 D365G) fused to mNeonGreen (mNG) at near endogenous FAR1 levels, or empty vector (EV) in sgFAR1 AC16 cardiomyocytes and grew these cells and wild-type cells under hypoxia to induce peroxisome degradation or normoxia^62^. We confirmed restoration of wild-type FAR1, as well as our LC3B-binding deficient FAR1 variant (FAR1 F148W), in sgFAR1 cells restored ether lipid abundance when compared to wild-type cells and this effect was lost in sgFAR1 cells stably expressing FAR1 D365G (Figure 4G and S12A-D). Hypoxia decreased peroxisome abundance in immunoblot assays as measured by GNPAT, PXMP2, FAR1, and AGPS abundance in EV-expressing wild-type cells and this effect was blocked in sgFAR1 cells expressing EV (Figure 4H). Re-expression of wild-type FAR1-mNG in sgFAR1 cells restored peroxisome degradation under hypoxia while re-expression of FAR1 F148W or FAR1 D365G did not (Figure 4H). These results suggest FAR1’s capacity to bind LC3B and support ether lipid metabolism are necessary for hypoxia to degrade peroxisomes.

Next, we asked whether increasing FAR1 would be sufficient to promote Keima-PEX processing in cells. mNG-FAR1 overexpression increased Keima-PEX processing at baseline when compared to cells expressing EV (S13A-B). To test whether FAR1’s capacity to bind LC3B is critical for FAR1 to promote Keima-PEX processing, we overexpressed our FAR1 F148W variant that disrupts its interaction with LC3B fused to mNG (mNG-Far1 F148W) (S13A-B). While wild-type FAR1 increased Keima-PEX processing at baseline, this effect was lost in cells expressing mNG-FAR1 F148W (Figure S13A-B).

To determine the specificity of FAR1 in peroxisome degradation, we tested whether FAR1 loss impacts the autophagy-dependent degradation of other organelles, such as mitophagy and ER-phagy. We stably expressed our previously reported Mito-Keima, in which Keima is fused to the outer mitochondrial membrane localization sequence of OMP25 (Keima-OMP25), in wild-type and sgFAR1 cardiomyocytes or wild-type cardiomyocytes stably expressing mNG-FAR1^67^. As expected, autophagy activation using Torin1 or hypoxia promoted Keima-OMP25 processing. FAR1 loss did not affect mitophagy induced by mTOR inhibition with Torin1 but reduced hypoxia-induced mitophagy (Figure S13C). Conversely, FAR1 overexpression was not sufficient to promote Keima-OMP25 processing at baseline and did not positively or negatively affect Keima-OMP25 processing under autophagy activation with Torin1 (Figure S13C). To measure ER-phagy, we stably expressed a previously validated ER-phagy reporter in which an N-terminal ER signal sequence is fused to monomeric RFP and GFP sequences and the ER retention sequence KDEL in wild-type and sgFAR1 cardiomyocytes and measured ER-phagy using fluorescence microscopy (Figure S13D-E)^50^. Upon ER-phagy activation, the ER is transported to the lysosome via autophagy and the GFP signal is quench in lysosomes and RFP puncta intensity indicate ER-phagy activity^50^. Autophagy activation via Torin1 increased ER-phagy in wild-type cardiomyocytes and this was unaffected by FAR1 loss (Figure S13D-E). Hypoxia did not significantly increase ER-phagy, at least in the cardiomyocytes that we studied (Figure S13D-E).

### Far1 is a direct HIF target gene and is necessary to degrade peroxisomes under chronic HIF activation

We and others have previously shown chronic HIF activation is sufficient to promote peroxisome degradation via autophagy through an unknown mechanism^26,35^. These results would suggest that HIF controls the expression of an unidentified peroxisome receptor. While our results in cells under acute hypoxia have shown FAR1 is degraded via autophagy, these experiments do not test whether HIF controls *FAR1* expression and whether FAR1 is necessary to degrade peroxisomes in the setting of chronic HIF activation. To address this, we asked whether *FAR1* is a transcriptional target of HIF. We analyzed publicly available genome-wide chromatin immunoprecipitation (ChIP) sequencing datasets that used an anti-HIF2α antibody and isogenic 786-O cells in which HIF2α was or was not eliminated using Crispr-Cas9 and ChIP experiments that utilized an anti-FLAG antibody and 786-O cells that have been modified to append a FLAG-HA epitope tag to the N-terminus of HIF2α expressed from the endogenous HIF2α locus to determine whether HIF binds the *FAR1* promoter^68^. Indeed, endogenous HIF2α bound the *FAR1* promoter in both datasets (Figure S14A). We confirmed HIF2α bound the *FAR1* promoter during hypoxia in an ARNT-dependent manner in ChIP assays in human AC16 cardiomyocytes (Figure 5A). Thus, FAR1 is a direct HIF target gene.

**Figure 5.**
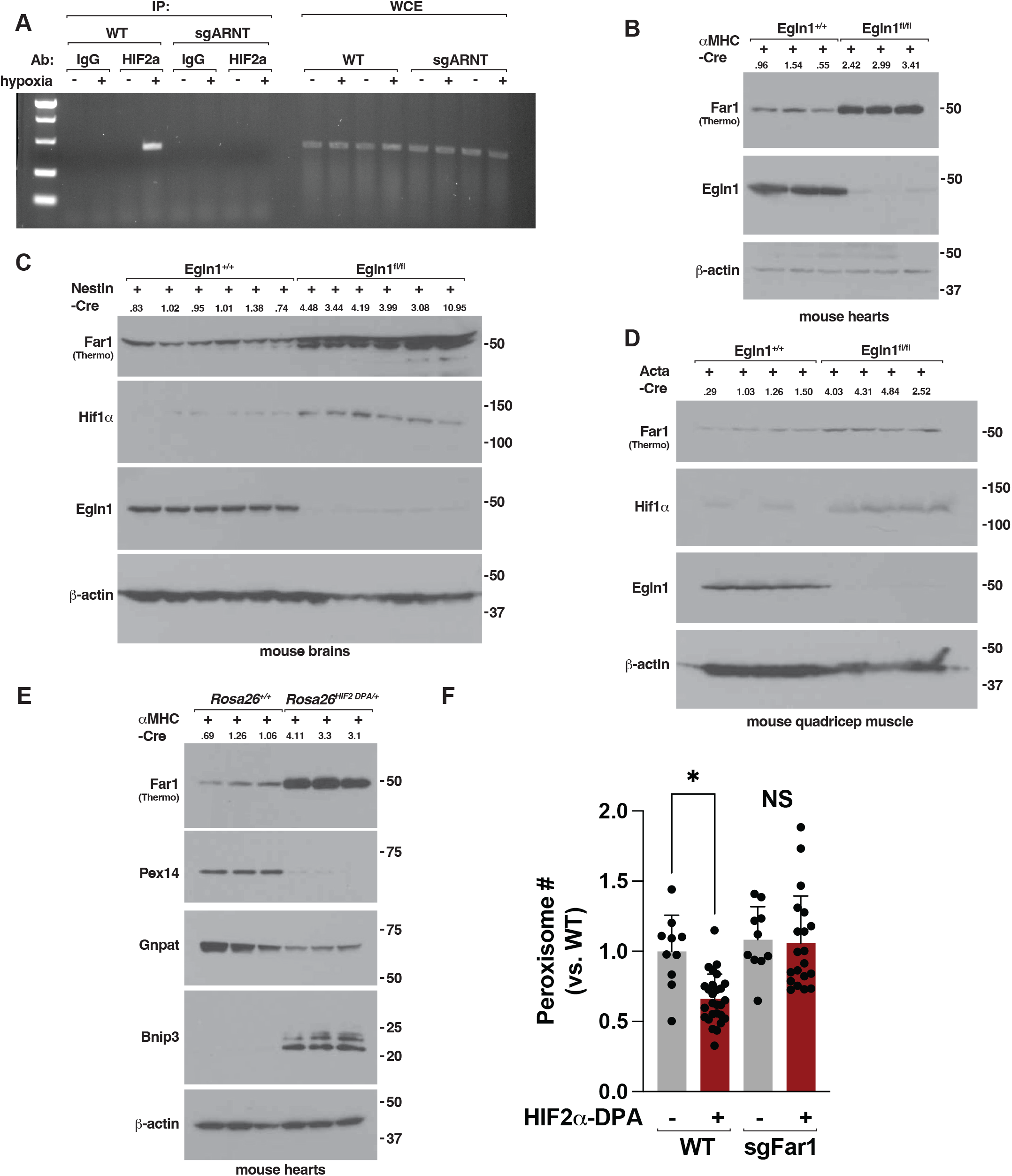
Far1 is increased in mouse models of chronic HIF activation. **(A)** HIF2α chromatin immunoprecipitation (ChIP) in wild-type (WT) or sgARNT human AC16 cardiomyocytes grown at hypoxia (0.1% O_2_) or normoxia for 18 hours. IgG was used as a control antibody. WCE, whole cell extract. **(B-E)** Immunoblot analysis of male and female 6–8-week-old **(B)** mouse hearts isolated from Egln1^+/+^ aMHC-Cre and Egln1^fl/fl^ aMHC-Cre mice, **(C)** mouse brains isolated from Egln1^+/+^ Nestin-Cre and Egln1^fl/fl^ Nestin-Cre mice, and **(D)** mouse quadricep muscles isolated from Egln1^+/+^ Acta-Cre and Egln1^fl/fl^ Acta-Cre mice. **(E)** Immunoblot analysis of mouse hearts isolated from male 12-week-old aMHC-Cre and Rosa26^HIF2DPA/+^ aMHC-Cre mice. Far1 immunoblots indicate which Far1 antibody was used to detect Far1. **(F)** Confocal microscopy analysis and quantification of Pex14-positive peroxisomes in wild-type or sgFar1 cardiomyocytes derived from aMHC-Cre and Rosa26^HIF2DPA/+^ aMHC-Cre mice. * indicates P<0.05. NS, non-significant. N>10 images per group.

We next asked whether chronic HIF activation increases FAR1. We inactivated *Egln1*, the primary HIFα hydroxylase, in mouse hearts, brains, and skeletal muscle^69^. As expected, *Egln1* loss increased HIF1α abundance in mouse brains and quadricep muscles. *Egln1* loss increased Far1 abundance in mouse hearts, brains, and quadricep muscles when compared to wild-type mice expressing αMHC-Cre, Nestin-Cre, or Acta-Cre (Figure 5B-D). As Egln1 has HIF-independent functions, we tested whether HIF activation alone is sufficient to increase Far1. We crossed αMHC-Cre mice with mice that carry a conditional HIF2α allele that contains a LoxP-Stop-LoxP cassette upstream of a cDNA encoding a non-hydroxylatable HIF2α (HIF2α dPA) fused to an HA epitope that escapes recognition by the VHL E3 ligase and is sufficient to activate HIF signaling in the presence of oxygen (Figure 5E)^29^. Hearts derived from HIF2α dPA expressing mice revealed increased Bnip3 abundance in immunoblot assays consistent with HIF activation when compared to hearts derived from wild-type αMHC-Cre expressing mice (Figure 5E). Hearts derived from HIF2α dPA expressing mice revealed increased Far1 abundance and decreased peroxisome abundance, as measured by Pex14 and Gnpat, when compared to hearts derived from wild-type αMHC-Cre expressing mice (Figure 5E). To test whether Far1 is necessary to degrade peroxisomes in the setting of chronic HIF activation, we isolated cardiomyocytes from wild-type and HIF2α dPA expressing mice and inactivated Far1 using Crispr-Cas9. Far1 loss increased peroxisome abundance in HIF2α dPA cardiomyocytes supporting HIF controls Far1 to promote peroxisome degradation (Figure 5F).

### Ether lipid precursors are sufficient to induce autophagy

Our loss of function studies in cells lacking components of ether lipid metabolism and our rescue experiments using an enzymatically dead FAR1 variant suggest these lipids play an important role in the degradation of peroxisomes under hypoxia. As ether lipids have been shown to increase under hypoxia, we hypothesize there exists a link between ether lipid metabolism and autophagy^70–72^. In mouse hearts isolated from animals housed under hypoxia, we found hypoxia elevated multiple alkyl-ether (ether) and alkenyl-ether (plasmalogen) phosphatidylcholine species, including PC O-32:1 (an alkyl phosphatidylcholine), PC O-36:1, PC O-38:3, and PC P-37:4 (an alkenyl phosphatidylcholine), in a Far1-dependent manner *in vivo* using lipidomic profiling (Figure 6A-B). This was specific to choline ether lipids as ester linked phosphatidylcholine abundance was either reduced or unchanged under these conditions nor did we detect an increase in ethanolamine ether lipids (Figure 6A-B). Far1 loss in the heart reduced alkyl-ether and alkenyl-ether phosphatidylcholine and ethanolamine species when compared to hearts derived from wild-type mice consistent with its key role in ether lipid metabolism (Figure S15A). To ask whether HIF mediated the elevation of choline ether lipids under hypoxia, we inactivated Arnt in mouse hearts. Arnt loss blocked HIF activation, as measured by Bnip3 abundance, and reduced but did not fully inhibit the increase in choline ether lipids in mouse hearts under hypoxia *in vivo* (6C and S15B). The increase in alkenyl-ether abundance under hypoxia was not due to an increase in PEDS1 activity (Figure S15C).

**Figure 6.**
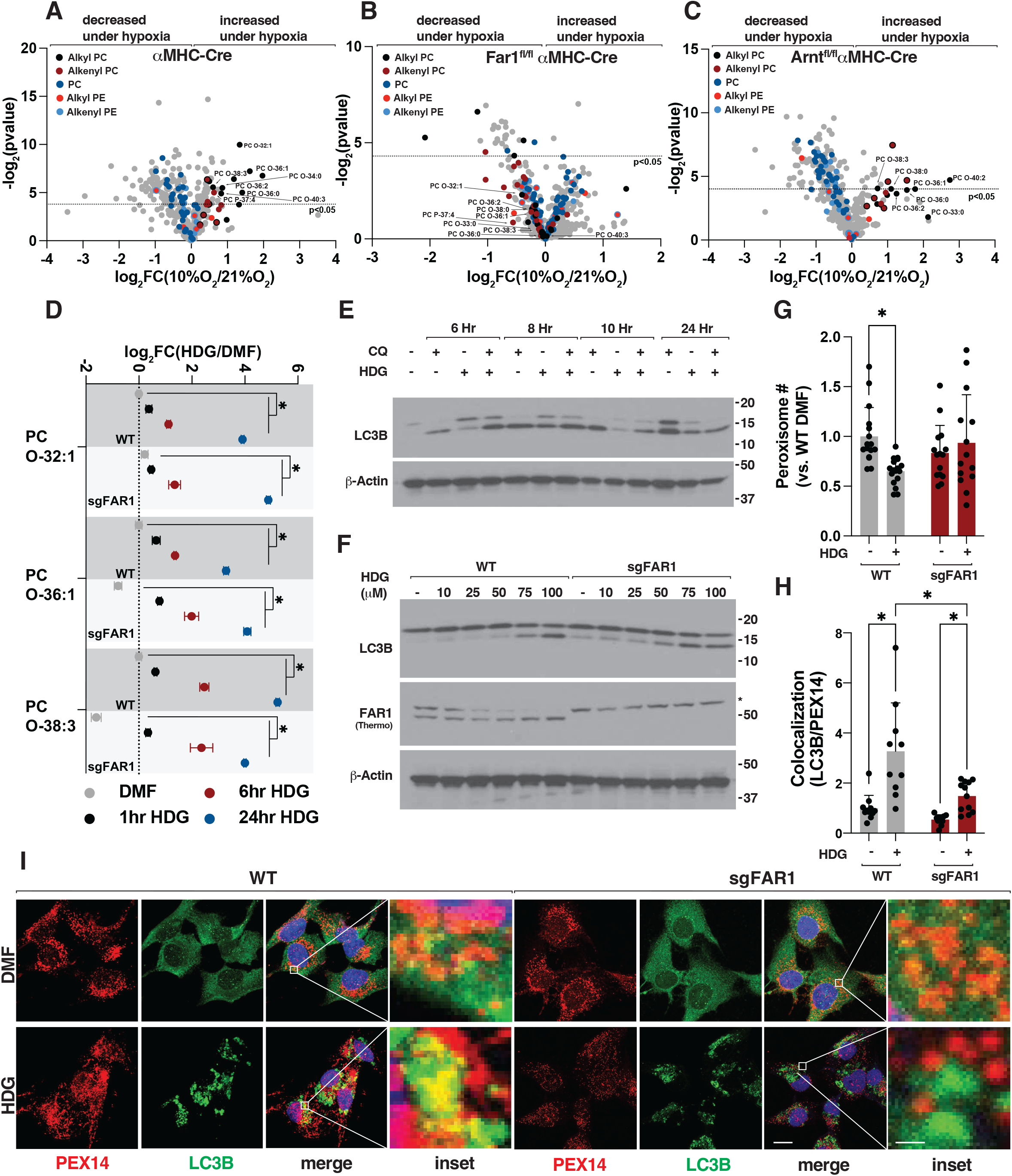
Increasing ether lipids using ether lipid precursors is sufficient to induce autophagy. (A-C) Volcano plots comparing normalized lipid intensities in mouse hearts isolated from (A) aMHC-Cre, (B) Far1^fl/fl^ aMHC-Cre, and (C) Arnt^fl/fl^ aMHC-Cre mice housed under 10% O_2_ (hypoxia) or 21% O_2_ (normoxia, normal atmospheric oxygen) for 7-days prior to euthanasia, tissue harvesting, and lipid extraction. Black dots indicate alkyl (ether) phosphatidylcholine (PC), dark red dots indicate alkenyl PC (plasmalogens), dark blue dots indicate ester linked PC species, red dots indicate alkyl (ether) phosphotidylethanolamine (PE), light blue dots indicate alkenyl PE (plasmalogens) species. Dots that are black and dark red indicate ether PC species that are annotated as either the alkyl or alkenyl form while dots that are red and light blue indicate ether PE species that are annotated as either the alkyl or alkenyl form. Dotted line indicates P<0.05. **(D)** Lipidomic profiling of alkyl PC species in wild-type and sgFAR1 human AC16 cardiomyocytes. Where indicated, cells were treated with 1-O-hexadecyl-sn-glycerol (100 uM, HDG) or DMF for indicated time points (hours, hr) prior to lipid extraction. N=3, *P<0.05. **(E-F)** Immunoblot analysis of wild-type and sgFAR1 (F) human AC16 cardiomyocytes. Where indicated, cells were treated with HDG (100 ׈M or indicated dose), chloroquine (CQ, 30 uM) (E) or vehicle. Unless otherwise stated, cells were treated with indicated compounds for 8 hours. *indicates non-specific band. Far1 immunoblot indicates which Far1 antibody was used to detect Far1. **(G)** Image analysis of PEX14 abundance in wild-type and sgFAR1 human AC16 cardiomyocytes. Where indicated, cells were treated with HDG (100 uM) or DMF for 8 hours prior to cell fixation. N>10, *P<0.05. **(H-I)** Image analysis (H) and confocal microscopy (I) of PEX14 and LC3B co-localization in wild-type sgFAR1 human AC16 cardiomyocytes. Where indicated, cells were treated with HDG (100 uM) or DMF for 8 hours prior to cell fixation. N>10, *P<0.05. Scale bar indicates 10 um, inset scale bar indicates 1 um.

Given that hypoxia increases choline ether lipid abundance in mouse hearts and hypoxia promotes peroxisome degradation via autophagy, we hypothesize increasing choline ether lipids may be sufficient to activate autophagy. To test this, we took advantage of the fact that ether lipid metabolism is compartmentalized into a FAR1-driven peroxisomal arm that then is completed in the ER. Supplementing cells with an ether lipid precursor, such as 1-O-hexadecyl-sn-glycerol (HDG), increases both choline and ethanolamine ether lipids by bypassing the FAR1-dependent peroxisomal steps of ether lipid synthesis and increasing ether lipid abundance through ER-dependent ether lipid metabolism^73^. We validated HDG supplementation increased many choline and ethanolamine alkyl-ether and alkenyl-ether lipids, including those that were increased in mouse hearts under hypoxia, such as PC O-32:1, PC O-36:1, and PC O-38:3, in a time and dose-dependent manner in wild-type and sgFAR1 cells (Figure 6D and S15D-F). In a time- and dose-dependent manner, HDG promoted autophagy flux in cells co-treated with chloroquine to inhibit lysosomal degradation as measured by LC3B lipidation using immunoblot assays and this required an intact autophagy pathway as this effect was lost in cells in which we inactivated key autophagy genes ATG7 or Beclin1 using Crispr-Cas9 (Figure 6E-F and S16A-E). HDG-mediated LC3B lipidation did not require FAR1 consistent with HDG acting downstream of FAR1-dependent ether lipid metabolism (Figure 6F and S16B, D). Consistent with this, HDG increased WIPI-2 puncta, a marker of the forming autophagosome, like Torin1 treatment (Figure S16F)^74^. We saw similar results with another ether lipid precursor distinct from HDG, 1-O-tetradecyl glycerol (TDG), which promoted LC3B lipidation in an ATG7-dependent manner to a similar degree as Torin1 or starvation (Figure S16D-E). HDG and TDG increased FAR1 abundance, where tested, in a dose-dependent manner (Figure 6F and S16D, G-I).

Next, we asked whether increasing ether lipids is sufficient restore peroxisome degradation in cells lacking FAR1. To test this, we measured peroxisome abundance by immunofluorescence and confocal microscopy following HDG treatment. HDG reduced PEX14-positive peroxisomes in wild-type cells consistent with peroxisome degradation in the setting of elevated choline ether lipids and this required FAR1 (Figure 6G). Accordingly, HDG treatment increased the co-localization of LC3B and PEX14-positive peroxisomes in wild-type cells but did not fully restore the colocalization of LC3B and PEX14 in cells lacking FAR1 (Figure 6H-I). These results support increasing choline ether lipids using ether lipid precursors activates autophagy, but this requires FAR1 to degrade peroxisomes.

### Ether lipid precursors promotes autophagy dependent ischemic protection

Whether increased ether lipids during hypoxia and their control of autophagy activation play an adaptive, detrimental, or homeostatic role in cellular function under hypoxia remains unclear. Prior studies in isolated rat hearts have shown HDG (also known as chimyl alcohol) reduced ischemia reperfusion (IR) injury ex vivo through an unknown mechanism^75^. Similarly, ether lipid precursors have been shown to protect cells in the setting of oxidative stress, neurodegeneration, cognition, and heart disease^76–79^. Our results in cardiomyocytes suggest that ether lipid precursors may provide tissue protection via autophagy activation. To test this, we perfused isolated Langendorff mouse hearts with HDG. When compared to vehicle, HDG perfusion increased LC3B lipidation in mouse hearts using immunoblot assays and protected mouse hearts in a Langendorff model of global myocardial ischemia, as reflected by a faster recovery (lowering) of end-diastolic pressure (LVEDP) (Figure 7A-B). We validated HDG increased choline and ethanolamine ether lipid abundance in mouse hearts using LC-MS-based lipidomics, including many of the same ether lipid species we detected in hearts isolated from mice housed under hypoxia, such as PC O-36:2, PC O-36:0, PC O-36:1, PC O-32:1, PC O-38:3, and PC O-38:0 (Figure 7C). To ask whether HDG protected mouse hearts against IR injury via autophagy activation, we treated Atg7^+/+^ aMHC-Cre and Atg7^fl/fl^-aMHC-Cre mice with HDG or vehicle followed by 45 minutes of left anterior descending artery (LAD) occlusion followed by reperfusion (IR). HDG treatment increased Lc3b lipidation in an Atg7-dependent manner and reduced infarct size in Atg7^+/+^ aMHC-Cre mice and this protection was lost in Atg7^fl/fl^-aMHC-Cre (Figure 7D-E). Collectively, these results suggest ether lipid precursors protect against ischemic injury via autophagy.

**Figure 7.**
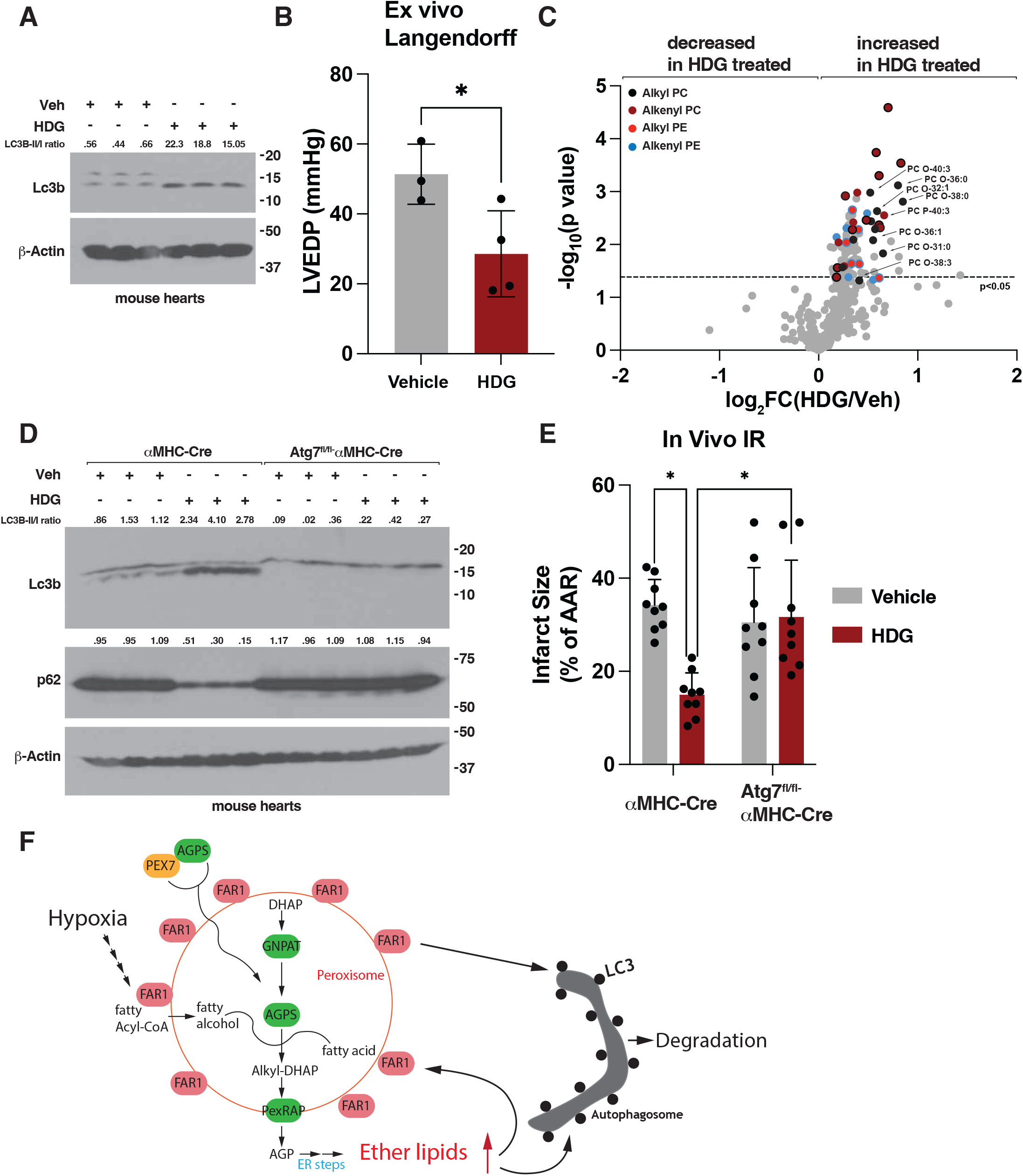
Increasing ether lipids using ether lipid precursors promotes autophagy dependent ischemic protection. **(A)** Immunoblot analysis of Langendorff mouse hearts perfused with 1-O-hexadecyl-sn-glycerol (100 uM, HDG) or vehicle containing ethanol. **(B)** Left ventricular end-diastolic pressure (LVEDP) in Langendorff assays after global I/R injury in wild-type C57BL/6J mice. Where indicated the hearts were infused with HDG (100 uM) or vehicle containing ethanol for 10 minutes prior to ischemia. N>3 per group, *P<0.05. **(C)** Volcano plots comparing normalized lipid intensities in mouse hearts isolated from mice treated with HDG (10 mg/kg, intraperitoneal injection) or vehicle once daily for 7-days prior to euthanasia, tissue harvesting, and lipid extraction. Black dots indicate alkyl (ether) phosphatidylcholine (PC), dark red dots indicate alkenyl PC (plasmalogens) species, red dots indicate alkyl (ether) phosphotidylethanolamine (PE), and light blue dots indicate alkenyl PE (plasmalogens) species. Dots that are black and dark red indicate ether PC species that are annotated as either the alkyl or alkenyl form while dots that are red and light blue indicate ether PE species that are annotated as either the alkyl or alkenyl form. Dotted line indicates P<0.05. **(D)** Immunoblot analysis of Atg7^+/+^ aMHC-Cre and Atg7^fl/fl^ aMHC-Cre mouse hearts. Where indicated, mice were treated with HDG (10 mg/kg, intraperitoneal injection) or vehicle once daily for 7-days prior to euthanasia and tissue harvesting. **(E)** Myocardial infarct size after in vivo cardiac ischemia/reperfusion (I/R) injury in Atg7^+/+^ aMHC-Cre and Atg7^fl/fl^ aMHC-Cre mice. Where indicated. mice were treated with HDG (10 mg/kg, intraperitoneal injection) or vehicle once daily for 7-days prior to ischemia. N>4 per group, *P<0.05. **(F)** Working model of hypoxia control of FAR1-dependent peroxisome degradation.

## Discussion

Our studies extend previous findings that HIF promotes peroxisome degradation, and we identify a candidate peroxisome receptor that is critical to regulate this process. Our results suggest FAR1 fulfills multiple key criteria to function as a selective receptor for hypoxia induced peroxisome degradation. Specifically, 1) autophagy is necessary and sufficient to control FAR1 abundance and FAR1 traffics to the lysosome under autophagy activation, 2) FAR1 is localized to the peroxisome membrane, 3) FAR1 binds to ATG8-family protein LC3B in a LIR-dependent manner thus providing a link between the peroxisome and the autophagy machinery, 4) FAR1 is a HIF target *in vivo*, and 5) FAR1 is necessary and sufficient to mediate peroxisome degradation under hypoxia and HIF activation. Thus, FAR1 is a key regulator of peroxisome abundance when oxygen is limiting.

Mutations in the human *PEX* genes in peroxisomal biogenesis disorders (PBDs), Zellweger syndrome spectrum (ZSS), neonatal adrenoleukodystrophy, infantile Refsum disease, and Rhizomelic chondrodysplasia punctata (RCDP) lead to neurodegeneration, bone development defects, liver failure, and early infantile death^1^. The primary defect in these settings remain unclear, however it has been suggested pexophagy is responsible for 65% of PBD cases^80^. We previously showed that chronic ischemia leads to constitutive pexophagy in the heart. It remains to be determined whether peroxisome loss in this setting is an adaptive, maladaptive, or homeostatic process, although peroxisomes are selectively essential under hypoxia and peroxisomal metabolism improves cardiac recovery following ischemia in animal models^70,75,81^. Low peroxisome number has been observed in various tumor cells, and HIF-driven clear cell renal carcinomas deplete peroxisomes via pexophagy^26,82^. Similarly, peroxisome-derived plasmalogens are elevated in aggressive cancers and promote ferroptosis sensitivity^83–88^. It will be necessary to test whether FAR1 inhibition alters the natural course of disease in these settings.

FAR1 is the rate-limiting enzyme in peroxisomal ether lipid metabolism, a class of lipids highly enriched in the brain, heart, and skeletal muscle^51,89^. FAR1 preferentially reduces C16 and C18 fatty acyl-CoAs into fatty alcohols that are further processed in the peroxisome via acyl/alkyl-DHAP reductase (*DHRS7B*) into ether lipid precursors that are completed in the ER^51,89^. While FAR1 is necessary and sufficient to mediate the degradation of peroxisomes under hypoxia, we find ether lipid metabolism is a critical component of peroxisome degradation under hypoxia. Specifically, loss of FAR1’s capacity to reduce fatty acyl-CoAs into fatty alcohols, loss of AGPS or GNPAT, peroxisomal enzymes that are required to generate the first ether linked intermediate in ether lipid synthesis, or loss of PEX7, which is required for peroxisomal AGPS localization, disrupts the capacity to degrade peroxisomes under hypoxia. In line with this, we find ether lipid precursors are sufficient to induce autophagy. Our studies have found hypoxia increases choline ether lipids abundance in a Far1-dependent manner, at least in the heart. As hypoxia has been shown to increase ether lipid abundance, ether lipids may act as a signaling molecule to increase autophagy under hypoxia (Figure 7F)^70–72^. While choline ether lipids are most abundantly found in the heart and skeletal muscle, ethanolamine ether lipids are more widely distributed across tissues^90^. Why hypoxia elevates choline, but not ethanolamine ether lipids, in the heart remains unclear, but choline ether lipids, like Platelet-Activating Factor (PAF), play an established roles in cell signaling^91^. Whether hypoxia controls ethanolamine ether lipids in other tissues remains undetermined.

We frequently detected decreased HIF activation under hypoxia in cells in which we inactivated FAR1 or other members of peroxisomal ether lipid metabolism, like AGPS and PEX7, using CRISPR-Cas9 or in mouse hearts lacking Gnpat. Where tested, the EglN-pVHL axis appeared functionally intact in these cells. As we find FAR1 is a HIF target gene, these results suggest HIF and ether lipid metabolism participate in a positive feedback loop. While how ether lipid metabolism may affect oxygen sensing by HIF remains unclear, lipoprotein-derived fatty acids have been shown to be a physiological modulator of HIF that function in parallel to oxygen suggesting lipid metabolism impacts HIF activity^92^.

Previous studies have reported ethanolamine plasmalogens are sensed in a negative feedback loop where FAR1 is degraded in the setting of elevated ethanolamine plasmalogens to control plasmalogen biosynthesis^51,93,94^. In our studies, we find rather FAR1 abundance is increased (using two validated FAR1 antibodies that detect different FAR1 epitopes) in cells treated with ether lipid precursors similar to a human FAR1 autosomal dominant variant (FAR1 R480C) that results in increased ether lipid synthesis and elevated FAR1 abundance (Figure 6F, S16D, and S16G-I)^51,93,94^. It was hypothesized that the FAR1 R480C variant results in a loss of feedback regulation by cellular ether lipids leading to its increased abundance, but our results suggest a different interpretation by which elevated ether lipids increase FAR1 abundance through an undetermined mechanism and increase autophagy and peroxisome degradation^94^. While peroxisomal β-oxidation was reported to be unchanged in FAR1 R480C patient fibroblasts, muscle biopsies from a FAR1 R480C patient revealed decreased abundance of autophagy-associated proteins in line with our results in which ether lipids activate autophagy-mediated degradation^95^. One of the FAR1 antibodies we tested (Thermo PA5-53585) detects a higher molecular weight (∼55kDA, referred to as p55) protein of unknown identity in immunoblot assays that is not lost in cells in which we inactivate FAR1 using Crispr-Cas9 (Figure 2E). The abundance of this unknown protein decreases in the setting of elevated ether lipids in a FAR1-dependent manner and is elevated in sgFAR1 cells (Figure 6F and S16J). How ether lipids are sensed in cells to control FAR1 abundance and the identity of the “plasmalogen sensor” remains unclear^93^.

Ether lipids increase cell survival against various stressors, and plasmalogens promote tissue protection in animal models^70,75–79^. Plasmalogen replacement therapy has been suggested to be treatment strategy for RCDP and ZSS^73,96,97^. Testing of a novel vinyl-ether synthetic plasmalogen, PPI-1040, in a RCDP mouse model restored plasmalogen levels in plasma to normal levels, increased plasmalogen abundance in peripheral tissues, and led to functional improvements in hyperactivity^98^. The mechanisms by which increased plasmalogens promote tissue protection is likely multi-factorial, but our work suggests ether lipid-mediated cardioprotection requires autophagy. While the mechanism by which elevating choline ether lipids activate autophagy remains undetermined, peroxisomal ether lipid synthesis has been shown to promote lifespan extension in *Caenorhabditis elegans*, and FAR1 overexpression in this context is sufficient to increase lifespan^99^. While uncontrolled ether lipid synthesis leads to features of hereditary spastic paraplegia, cerebral palsy, and cataracts, elevated ether lipids are a marker of extreme longevity in humans^94,100^. As autophagy activation alone is sufficient to prolong lifespan in mice, increasing ether lipids pharmacologically may function as autophagy activators and promote healthy aging^56^.

### Limitations of the study

While FAR1 is required for peroxisome degradation under hypoxia, it remains possible other receptors exist that contribute to hypoxia-induced pexophagy. This is consistent with other organelles, such as the ER and mitochondria, in which multiple partially redundant receptors control ER-phagy and mitophagy^40,48,50,101–106^. Several mechanistic details of the described pathway remained unresolved, such as how hypoxia specifically regulates choline ether lipid abundance, whether this is true in tissues other than the heart, and how increasing choline ether lipids activates autophagy. As our studies using ether lipid precursors elevate both choline and ethanolamine ether lipids, it will be necessary to devise a strategy to modulate choline ether lipids alone to direct test their role in autophagy. Despite these limitations, our findings that FAR1 acts as a central component of hypoxia promotes peroxisome degradation are supported by *in vivo* and *ex vivo* biochemical and genetic studies, including genetic knockouts of ether lipid metabolism, pexophagy flux assays, *in vivo* and *ex vivo* protein-protein interactions, and lipidomic profiling.

## Acknowledgements

We thank the members of the Wyant laboratory, R. Soberman (MGH), W. Kaelin (DFCI), R. Wolfson (HMS), N. Braverman (McGill), and J. Hacia (USC) for helpful discussions and critical reading of the manuscript. Special thanks to the Cardiovascular Physiology Core at Mass General Brigham/Harvard Medical School (HMS) for assistance with cardiac physiology experiments and Mass General Brigham Nikon Center of Excellence for helpful discussions and imaging experimental design. M.S., together with G.A.W, performed and designed experiments, and analyzed and assembled data. S.Q. performed molecular cloning and mRNA analysis. G.G., with the with the help of G.A.W, J.B., and S.Q., performed mouse biochemical experiments and mRNA analysis. N.M. and K.W. performed plasmalogen abundance analysis and PEDS1 activity analysis. O.S., H.K., and S.C., performed proteomic data acquisition and analysis. A.D., and C.B.C performed liquid chromatography mass spectrometry data acquisition and analysis. G.A.W. wrote the article.

## Resource Availability

Additional information or requests for resources will be fulfilled by the lead contact, Gregory Wyant. All plasmids generated in this study are available via addgene or fulfilled by the lead contact. The original mass spectra, spectral library, and the protein sequence databases used for searches have been deposited in the public proteomics repository MassIVE (http://massive.ucsd.edu) and are accessible at ftp://MSV000097181@massive.ucsd.edu when providing the dataset password: hypoxia. If requested, also provide the username: MSV000097181. These datasets will be made public upon acceptance of the manuscript.

## Methods

### Cell Culture

U2OS osteosarcoma cells and C2C12 mouse myoblasts were originally obtained from the American Type Culture Collection (ATCC). 293FT human embryonic kidney cells were originally obtained from Invitrogen. AC16 human cardiomyocytes were originally obtained from Sigma-Aldrich. 293FT and C2C12 cells were maintained in Dulbecco’s minimum essential medium (DMEM) supplemented with 10% fetal bovine serum (FBS), penicillin (100 U/ml), and streptomycin (100 ׈g/ml). U2OS cells were maintained in McCoys medium supplemented with 10% fetal bovine serum (FBS), penicillin (100 U/ml), and streptomycin (100 ׈g/ml). AC16 cells were maintained in DMEM/F12 medium supplemented with 15% FBS, penicillin (100 U/ml), and streptomycin (100 ׈g/ml). All experiments using AC16 cells and C2C12 cells were performed with passage 4-6 cells. Fresh aliquots of 293FT were thawed every 4 weeks. Neonatal cardiomyocytes were isolated from mouse cardiac tissue (see below) and were maintained in DMEM supplemented with 15% FBS, penicillin (100 U/ml), and streptomycin (100 ׈g/ml). Virally infected cells were selected with puromycin (5 µg/ml) or blasticidin (10 µg/ml) as appropriate for the vector used. All mammalian cells were grown in a humidified atmosphere containing 21% oxygen and 5% CO_2_ at 37 °C unless otherwise stated.

### Chemicals

Chloroquine (Selleck) was prepared as a stock solution of 30 mM in dH_2_O and was diluted in cell culture medium at the indicated concentration. Torin1 (CST) was prepared as a stock solution of 1 mM in DMSO and was diluted in cell culture medium at the indicated final concentration. MitoTracker DeepRed FM (CST) was prepared at a 1 mM stock concentration in DMSO and was diluted in cell culture media at a final concentration of 500 nM. LysoTracker Red DND-99 (Thermo) came prepared at a 1 mM stock concentration and was diluted in cell culture media at a final concentration of 100 nM. LysoTracker Blue DND-2 (Thermo) came prepared at a 1 mM stock concentration and was diluted in cell culture media at a final concentration of 100 nM. 1-O-hexadecyl-sn-glycerol (HDG, Fisher) was prepared at a 15 mM stock concentration in 100% ethanol or at a 50 mM stock concentration in DMF. 1-O-Tetradecyl Glycerol (TDG, GoldBio) was prepared at a 50 mM stock concentration in DMF. SAR405 (Selleck) was prepared as a stock solution of 200 mM in DMSO and was diluted in cell culture medium at the indicated final concentration. Bafilomycin A1 (BafA1, CST) was prepared as a stock solution of 1 mM in DMSO and was diluted in the cell culture medium at the indicated concentration. FCCP (Selleck) was prepared as a stock solution of 20 mM and was diluted in cell culture medium at the indicated final concentration. Oligomycin (CST) was prepared as a stock solution of 5 mM in DMSO and was diluted in cell culture medium at the indicated final concentration. TAK-243 (Selleck) was prepared as a stock solution of 190 mM in DMSO and was diluted in cell culture medium at the indicated final concentration. MG-132 (CST) was prepared as a stock solution of 1 mM in DMSO and was diluted in cell culture medium at the indicated final concentration. Hydrogen peroxide (Fisher) was prepared at 1 M stock concentration water and was diluted in cell culture medium at the indicated concentration. N-acetyl-cysteine (Tocris) was prepared at a 50 mM stock concentration in water and was diluted in cell culture medium at the indicated concentration. Tunicamycin (CST) was prepared at a stock concentration of 5 mg/mL in DMSO and was diluted in cell culture medium at the indicated concentration. AICAR (CST) was prepared at a stock concentration of 75 mM in water and warmed at 37 °C and was diluted in cell culture medium at the indicated concentration.

To maintain HDG (1-O-hexadecyl-sn-glycerol) and TDG (1-O-tetradecyl glycerol) activity, it is essential that 1-O-hexadecyl-sn-glycerol and 1-O-tetradecyl glycerol are immediately prepared in ethanol or DMF upon arrival and stored at -80 °C or immediately stored at -80 °C until preparation. HDG and TDG was individually aliquoted to experimental volumes and was not refrozen or re-used.

### Antibodies

Primary antibodies used were: Rabbit anti-FLAG (CST #14793), Mouse anti-FLAG (CST #8146), Rabbit anti-p70 S6 Kinase (CST #9202), Rabbit anti-LAMP1 (CST #9091), Mouse anti-LAMP2 (Santa Cruz sc-18822), Rabbit anti-PEX14 (Proteintech 10594-1-AP), Mouse anti-b-Actin (Santa Cruz sc-47778), Rabbit anti-HIF1A (CST #14179), Rabbit anti-LONP2 (ProteinTech 18035-1-AP), Rabbit anti-GNPAT (ProteinTech 14931-1-AP), Rabbit anti-BNIP3 (CST #3769), Rabbit anti-BNIP3 (CST #44060), Rabbit anti-NDRG1 (CST #5196), Mouse anti-SQSTM1/p62 (Abcam ab56416), Rabbit anti-LC3B (CST #43566), Mouse anti-LC3B (CST #83506), Rabbit anti-catalase (CST #14097), Mouse Anti-Keima-Red (MBL International, M182-3M), Rabbit anti-HIF2A (CST #71565), Rabbit anti-PEX19 (ProteinTech 14713-1-AP), Rabbit anti-MLSTD2 (Far1) (Thermo PA5-53585), Rabbit anti-FUNDC1 (Abcam ab74834), Rabbit anti-ATG7 (CST #8558S), Rabbit anti-YIPF3 (Sigma #HPA014859), Rabbit anti-mCherry (CST #43590S), Rabbit anti-AGPS (ProteinTech 21011-1-AP), Rabbit anti-Ubiquitin (CST #3933S), Mouse anti-CHOP (CST #2895S), Rabbit anti-TEX264 (ProteinTech 25858-1-AP), Rabbit anti-NBR1 (ProteinTech 16004-1-AP), Rabbit anti-PEX7 (ProteinTech 20614-1-AP), Rabbit anti-COPB1 (ProteinTech 27469-1-AP), Rabbit anti-PHD2/EGLN1 (CST #4835S), Rabbit anti-HA (CST #3724S), Rabbit anti-SQSTM1/p62 (CST #39749S), Rabbit anti-FAR1 (CST #38678), Rabbit anti-BNIP3 (CST #3769, mouse specific), Rabbit anti-PXMP2 (Proteintech 24801-1-AP), Rabbit anti-GST (CST #2622), Rabbit anti-GM130 (CST #12480), Rabbit anti-PHB1 (CST #2426), Rabbit anti-S6 (CST #2217), Rabbit anti-Ubiquitin (CST #58395), Rabbit anti-Catalase (CST #14097), Rabbit anti-PEX16 (Proteintech 14816-1-AP), Rabbit anti-ACSL4 (CST #38493). All primary antibodies used for immunoblot were diluted at 1:500 in TBS-T+5% BSA.

### cDNA Synthesis and Quantitative PCR analysis (qPCR)

Total RNA was extracted from AC16 cells and mouse tissues using TRIzol Reagent (Life Technologies, 15596). cDNA was generated by reverse transcription using AffinityScript qPCR cDNA Synthesis kit (Agilent) according to manufacturer’s instructions. qPCR was performed using a QuantStudio 6 (Thermo) using Sybrgreen according to the manufacturer’s instructions. All quantitative calculations were performed using the 2^-ΔΔCt^ method using *b-actin* or *GAPDH* as a reference gene.

The following human qPCR primers were used:

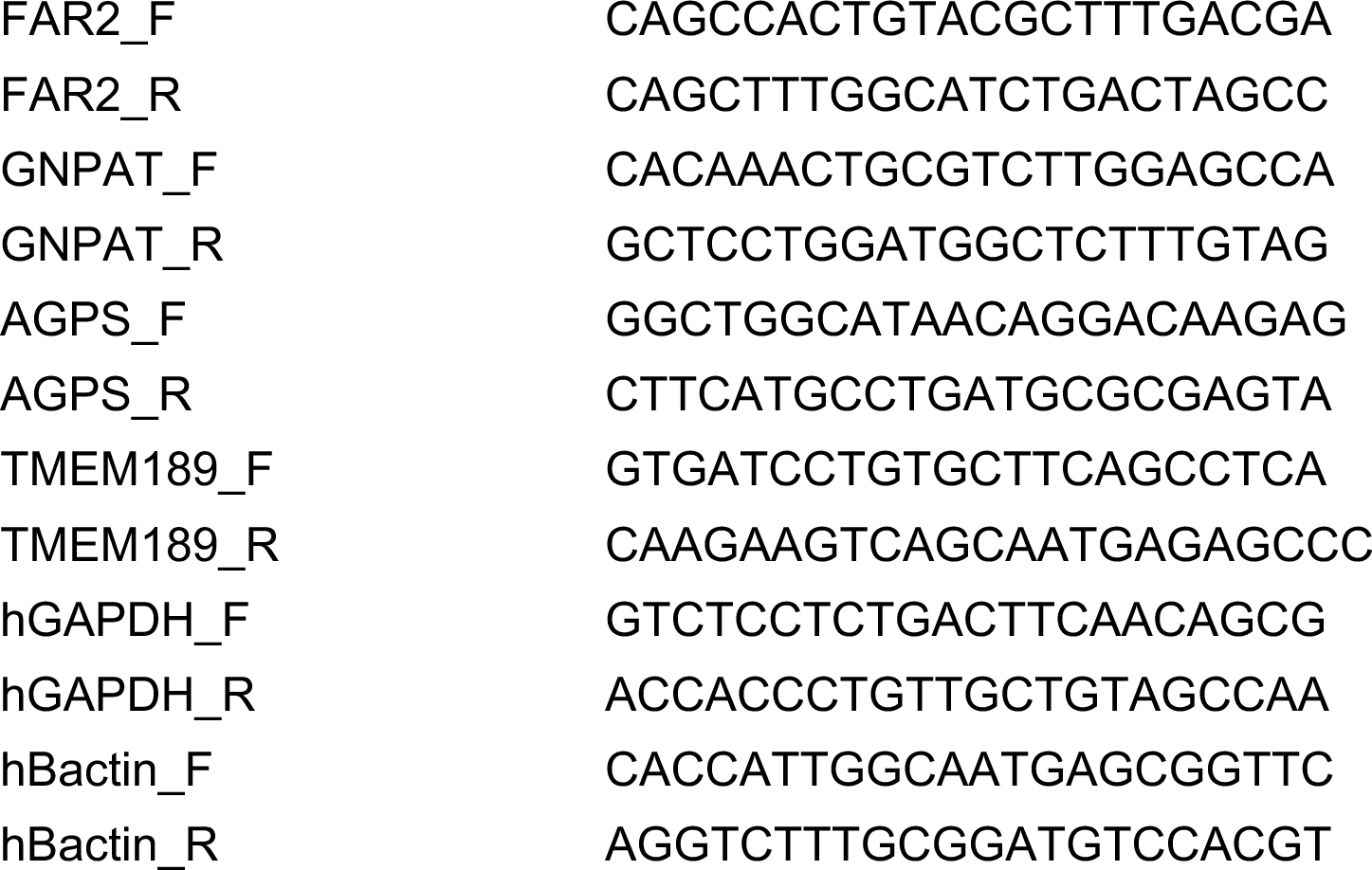

The following mouse qPCR primers were used

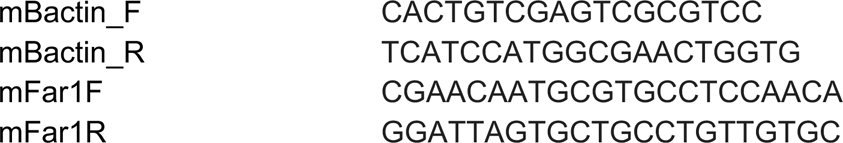

### Cloning of cDNA expression vectors

To make the Far1 expression vector pMXS-mNG-Far1, the mNG-Far1 fusion cDNA was synthesized as a dsDNA gBlock by Integrated DNA Technologies (IDT). The synthetic dsDNA fragment was then PCR amplified using the following primers to introduce a 5’Xhol and 3’NotI site.

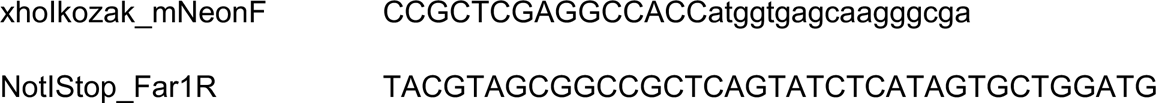

The resulting PCR products was then digested with XhoI and NotI, gel purified, and ligated into pMXS linearized with these two enzymes and transformed into XL-10 gold bacteria. Ampicillin-resistant colonies were screened by XhoI and NotI digestion and verified by Sanger sequencing.

To make the LC3B expression vectors pMXS-FLAG-LC3B and pMXS-FLAG LC3B K51A, the FLAG-LC3B fusion cDNA and K51A point mutant was synthesized as a dsDNA gBlock by integrated DNA Technologies (IDT). The synthetic dsDNA fragment was then PCR ampliflied using the following primers to introduce a 5’EcoRI and 3’NotI site.

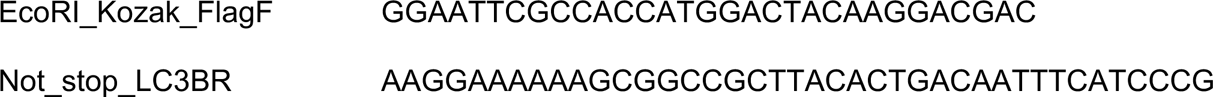

The resulting PCR products was then digested with EcoRI and NotI, gel purified, and ligated into pMXS linearized with these two enzymes and transformed into XL-10 gold bacteria. Ampicillin-resistant colonies were screened by EcoRI and NotI digestion and verified by Sanger sequencing.

### Site-Directed Mutagenesis of FAR1 variants

FAR1 missense variants were generated using the Q5 Site-Directed Mutagenesis Kit (New England Biolabs) according to the manufacturer’s instructions with pMXS-mNG-FAR1 as the template and verified by sanger sequencing before generation of stable cell lines. The following primers were used:

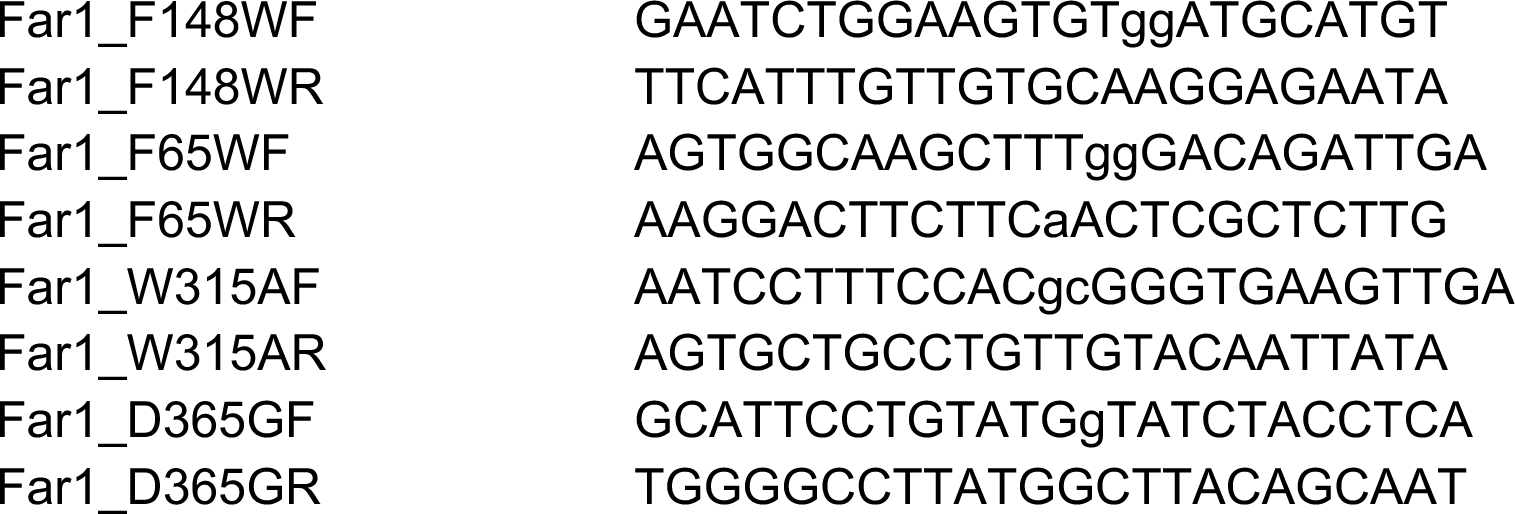

### Plasmids

The following plasmid was purchased from addgene. pCW57-CMV-ssRFP-GFP-KDEL was a gift from Noboru Mizushima (Addgene plasmid #128257; http://n2t.net/addgene:128257; RRID:Addgene_128257). pLJC5-3XHA-EGFP-PEX26 http://n2t.net/addgene:139059 ; RRID:Addgene_139059).

### CRISPR/Cas9 Plasmid Generation

The LentiCRISPR_v2-puromycin or -blasticidin vectors were used to express sgRNAs with Cas9. LentiCRISPR_v2 was digested with BsmBI, gel purified, and ligated with annealed oligonucleotides. The px459 vector was used to expressed sgRNAs with Cas9 to generate single cell crispr clones. Px459 was digested with BsmBI, gel purified, and ligated with annealed oligonucleotides.

Sense and antisense oligonucleotides corresponding to the desired sgRNA were mixed at equimolar ratios (0.25 nanomoles of each sense and antisense oligonucleotide) and annealed by heating to 100 °C in annealing buffer (1X T4 Ligase buffer, T4 PNK) followed by slow cooling to 30 °C over 3 hours. The annealed oligonucleotides were then diluted 1:200 in dH2O and ligated into the digested CRISPR vectors by incubation with T4 DNA ligase for 30 minutes at room temperature. The ligation reaction was transformed into XL-10 Gold ultracompetent cells and ampicillin-resistant colonies were verified by Sanger sequencing.

The following sgRNAs were used:

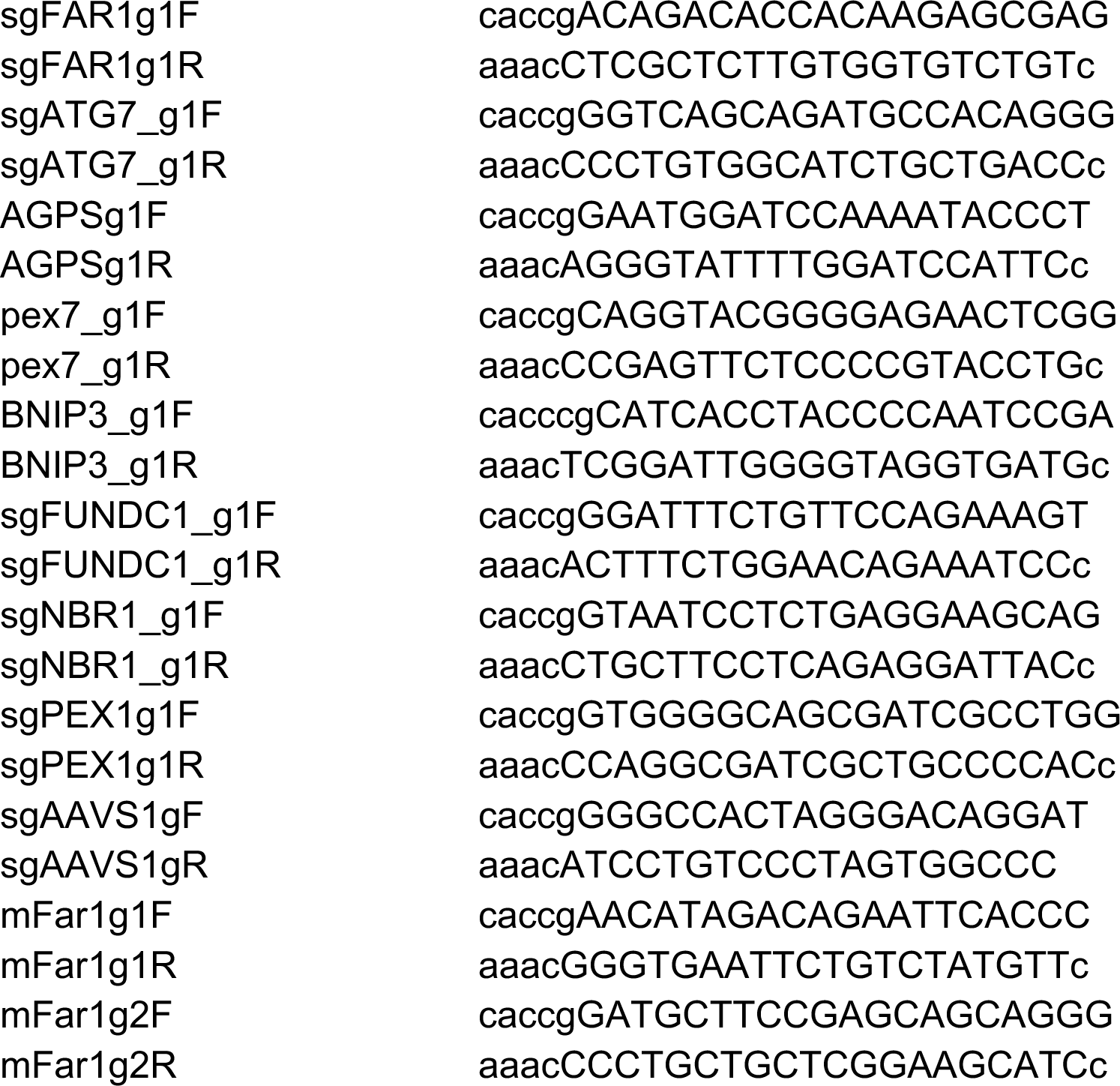

### Lentivirus Generation and Infection

To make the lentiviruses, 1.3 x 10^6^ HEK293FT cells were seeded in a 6-cm plate in DMEM supplemented with 10% FBS. Twenty-four hours later the cells were transfected with 1 mg of the desired lentivirus encoding plasmid together with of the packaging plasmids 0.5 μg psPAX2 and 0.5 μg pMD2.G using XtremeGene9 (Sigma) transfection reagent. Twelve hours after transfection, the medium was aspirated and replaced with 4 mL of fresh medium. Thirty-six hour later the virus-containing supernatants were collected and passed through a 0.45-μm filter to eliminate contaminating mammalian cells.

For lentiviral infections, 1 x 10^6^ cells target cells were combined with 250 μl of virus in 2 mL total volume of medium supplemented with 8 mg/mL polybrene and plated in 6-well plates. The plates were then immediately spun in an Eppendorf 581R centrifuge at 2,200 r.p.m. for 45 minutes at 37 °C. Twelve hours after infection, the virus-containing media was aspirated and replaced with fresh DMEM+10% FBS supplemented with Pen/Strep. Twenty-four hours post infection, the infected cells were trypsinized and replated in media that contained 5 mg/mL puromycin or 10 mg/mL blasticidin as appropriate for that virus.

For lentiviral infections involving mouse neonatal cardiomyocytes, cells were transduced with virus one day after isolation using low glucose DMEM + 7.5% FBS supplemented with polybrene (8 mg/ml) and Pen/Strep.

### Immunoblot Analysis

Cells grown in 10-cm tissue culture dishes were washed once on ice with ice cold 1x phosphate-buffered saline (PBS) and then 1 mL ice cold PBS was added to each plate. The cells were then detached by scraping, transferred to a 1.5 mL Eppendorf tube, and pelleted at 1500 x g for 1 minute at 4 °C. The PBS was then aspirated, and the cell pellets were resuspended in lysis buffer (40 mM Hepes pH 7.4, 150 mM NaCl, 1.5 mM MgCl_2_, 1% Triton-X 100, Complete Mini EDTA-free Protease Inhibitor (Sigma), and Phosstop Phosphatase Inhibitor (Sigma)) and lysed by gentle rocking for 20 minutes at 4 ׈C. For experiments involving Keima-fusion proteins, cell pellets were resuspended in RIPA buffer and lysed by gentle rocking for 20 minutes at 4 ׈C. The lysate was then clarified by centrifugation at 17,000 x g for 10 minutes at 4 ׈C and transferred to a new Eppendorf tube. The protein concentration of the whole cell extract was measured using the Bradford Assay and then normalized to 2 mg/mL. After normalization, the whole cell extract was denatured by the addition of 2.2% SDS, 11% glycerol, 100 mM DTT, and bromophenol blue.

Samples were resolved by SDS-polyacrylamide gel electrophoresis using 4-20% or 4-12% Tris-Glycine gels (Novex) and transferred onto nitrocellulose membranes using a Bio-Rad Trans-Blot Turbo Transfer system using the high molecular weight setting. Following transfer, membranes were incubated with Ponceau S staining solution (CST) for 3 minutes rocking at room temperature to visualize transferred bands following by washing in Tris-buffered saline +0.1% Tween 20 (TBS-T). Membranes were then blocked by incubation in 5% milk/tris-buffered saline + 0.1% Tween 20 with gentle rocking for 1 hour at room temperature. The membranes were washed with TBS-T prior to overnight incubation with primary antibody diluted in TBS-T+5% BSA with gentle rocking at 4 ׈C. After primary antibody incubation, membranes were washed three times with TBS-T and then incubated with horseradish peroxidase (HRP)-conjugated secondary antibody (CST) (1:5000) in 5% milk/TBS-T with gentle rocking for 1 hour at room temperature. The membranes were washed three times with TBS-T and bound antibodies were then detected with enhanced chemiluminescence western blotting reagents (Thermo Fisher Scientific, no. WBKLS0500) or Super-Signal West Pico (Thermo Fisher Scientific, no. PI34078).

Quantification of immunoblots was performed in ImageJ using the analyze gel function and are shown as relative values compared to the control or untreated group. For mouse experiments shown in Figure 1F and 1H, 4F, 5B-E, each blot shows at least three biological replicate mice per treatment group. Each band was quantified individually. The band intensities for the ad libitum fed wild-type control group were averaged and each mouse was then normalized to the average band intensity of the ad libitum fed wild-type control group. Band intensity for each mouse was then normalized to the control protein β-actin. Values are shown as fold change versus ad libitum fed wild-type control.

For quantification of FAR1 abundance shown in Figure 2C-D, Figure 6F, and Figure S19F-I, the FAR1 band intensities were first normalized to the untreated wild-type control and followed by normalization to the control protein β-actin. Values are shown as fold change versus wild-type control. For quantification of peroxisomal protein abundance shown in Figure 4G, peroxisomal protein immunoblots were first normalized to the untreated wild-type control grown at normoxia followed by normalization to the control protein β-actin. Values are shown as fold change versus wild-type normoxia. Quantification of LC3B immunoblots is shown as the ratio of LC3B-II: LC3B-I. Both bands of LC3B were quantified individually and each ratio was normalized to the wild-type control grown in full media at normal atmospheric oxygen or to the wild-type ad libitum fed normal atmospheric oxygen control group.

For quantification of immunoprecipitation experiments shown in Figure 2, band intensity for immunoprecipitated prey protein is normalized to the band intensity of the immunoprecipitated bait protein. Both bands were quantified individually.

For quantification of Keima processing experiments, the ratio of “processed Keima: Full length Keima fusion protein” is shown. Each ratio was normalized to wild-type cells grown in full media at normoxia. Both bands were quantified individually.

Anti-MLSTD2 (FAR1, Thermo) and Anti-FAR1 (CST) immunoblots were incubated for 48 hours at 4 °C prior to secondary antibody. Anti-MLSTD2 recognizes two bands of ∼50kDa and ∼55 kDa (p55) in size in human AC16 cells and 293FT cells, and in mouse brains. At times this band was detected in mouse cardiomyocytes, however it was not consistently detected. The ∼50 kDa immunoreactive band is specific to FAR1 as it is lost in sgFAR1 AC16 and sgFAR1 293FT cells generated using Crispr-Cas9. Anti-FAR1 (CST) recognizes a single band at ∼50 kDa that is specific to FAR1 as it is lost in sgFAR1 AC16 and sgFAR1 293FT cells generated using Crispr-Cas9. Anti-FAR1 (CST) does not recognized Far1 in mouse tissues. Mouse hearts must be perfused with PBS prior to tissue lysis when blotting for anti-MLSTD2 (Far1) as we found non-specific contaminating serum proteins immunoreact with anti-MLSTD2 obscuring the MLSTD2 (Far1) band.

### Subcellular Fractionation

Cells were plated in 10-cm plates at a density of 4 x 10^5^ cells per plate and grown to 80% confluence. At the time of collection, the cells were washed twice with ice cold PBS and once with ice cold Hepes-sucrose buffer (HSB) (250 mM sucrose, 20 mM Hepes-KOH pH 7.4) containing 1 mM EDTA and protease inhibitors. Next, 400 μL of HSB containing 100 mg/mL digitonin (Wako Chemicals) was added to each plate and scraped into a 1.5 mL Eppendorf tube and incubated on ice for 10 minutes. Cells were divided into two different 1.5 mL Eppendorf tubes (180 mL/tube) for whole cell extract and a cytosolic/membrane fraction. The whole cell extract was solubilized by the addition of 36 mL of 6x SDS-PAGE sample buffer. The cytosolic/membrane fraction was centrifuged at 20,000 x g for 15 minutes at 4 ׈C. After centrifugation, 150 μL of the supernatant (cytosolic fraction) was transferred into a new 1.5 mL Eppendorf tube and solubilized by the addition of 33 mL 6x SDS-PAGE sample buffer. The remaining pellet was washed once in 50 μL HSB and then centrifuged at 20,000 x g for 15 minutes at 4 ׈C. After centrifugation, the HSB was aspirated and 200 μL of 1x SDS-PAGE sample buffer was added to the membrane pellet and sonicated. All samples were boiled for 5 minutes prior to western blotting.

### Peroxisome Immunoprecipitation (Peroxo-IP)

Cells were plated in 15-cm plates at a density of 4 x 10^5^ cells per plate and grown to 80% confluence. At the time of collection, the cells were washed twice with ice cold KPBS (136 mM KCl, 10 mM KH_2_PO4, pH 7.25, supplemented with Peirce Protease Inhibitor Tablets (Thermo, A32965)), scraped into 1 mL KPBS and collected into an Eppendorf tube and centrifuged at 4C at 1000xg for 2 min. Pelleted cells were resuspended in 1 mL KPBS and then 50 uL of resuspension was collected into a new Eppendorf as a whole cell extract (WCE) and lysed with 100 uL 1% Triton-X100 lysis buffer (40 mM Hepes pH7.4, 150 mM NaCl, 1% Triton-X100, 150 mM MgCl_2_). The remaining whole cell suspension was then dounce homogenized 30 times on ice in a 2 mL homogenizer. The homogenate cells were then collected in a new Eppendorf tube and centrifuged at 1000xg for 2 min at 4׈C to pellet nuclei and unbroken cells while cellular organelles remained in the supernatant. The supernatant was collected into a new Eppendorf and mixed with 100 uL anti-HA magnetic beads (Pierce) and incubated rocking at 4 ׈C for 5 minutes before being gently washed three times in 1 mL KPBS using a DynaMag Spin Magnet, changing tubes between each wash. Beads containing bound peroxisomes were resuspended in 100 uL 1% Triton-X100 lysis buffer to extract proteins and incubated at 4׈C for 10 minutes. Samples were then clarified by centrifugation at max speed for 10 min at 4׈C and clarified lysates were mixed with SDS sample loading dye.

### Chromatin Immunoprecipitation (ChIP) PCR

To assess HIF2α bound to the FAR1 promoter, HIF2α was chromatin immunoprecipitated using HIF-2α XP anti-rabbit (CST #59973) or IgG (CST #2729) and the SimpleChIP Plus Enzymatic Chromatin IP Kit (CST #9005) according to manufacturer’s instructions. To prepare chromatin, AC16 human cardiomyocytes were plated in 15-cm plates at a density of 8 x 10^5^ cells per plate and grown until 80% confluence. At time of cell harvest, media was aspirated and 20 mL of fixation solution (540 ׈L of 37% formaldehyde was added to 20 mL DMEM) was added to each plate and cells were incubated at room temperature gently shaking for 10 minutes. Following fixation, 2 mL 10X Glycine solution was added to each plate and placed at room temperature for 5 minutes. The glycine solution was then aspirated after which the cells were washed twice in ice cold PBS followed by the addition of 2 mL PBS containing protease inhibitors. The cells were then collected via scraping and transferred to a 15 mL conical tube. The tube was then centrifuged at 2000 x g for 5 minutes at 4C and the supernatant was aspirated without disturbing the pellet. Following, the pellet was resuspended in 1mL Buffer A containing DTT and protease inhibitors and left on ice for 10 minutes and mixed by inversion every 3 min. The nuclei were then pelleted by centrifugation at 2000xg for 5 minutes at 4׈C and the supernatant was aspirated, and the pellet was resuspended in 1 mL ice cold Buffer B containing DTT and transferred to a 1.5 mL Eppendorf tube. At this stage, 0.5 uL micrococcal nuclease was added to each Eppendorf tube and incubated for 20 minutes at 37׈C to digest DNA to a length of 150-900 bp. During the 20 minutes incubation, the Eppendorf tube was mixed by inversion every 3 minutes. Following incubation, the digestion was stopped via the addition of 10 uL 0.5 M EDTA to each Eppendorf and placing the tube on ice for 2 mins. The nuclei were then pelleted by centrifugation at 16,000 x g for 1 minutes at 4׈C and the supernatant was aspirated without disturbing the pellet. The nuclear pellet was then resuspended in 100 uL ChIP buffer containing protease inhibitors and incubated on ice for 10 minutes. After 10 minutes, the resuspended nuclei pellet was sonicated on wet ice in a 4׈C walk in cold room for 3 x 30 seconds with 30 seconds on wet ice between pulses. Following sonication, the nuclei lysates were clarified by centrifugation at 9,400 x g for 10 minutes at 4׈C. The supernatant containing the cross-linked chromatin was then collected and transferred to a new Eppendorf tube. At this stage, ∼ 50 uL of the chromatin preparation was placed into a new tube for analysis of chromatin digestion and chromatin concentration. To prepare chromatin immunoprecipitation, 5 ug of digested, crosslinked chromatin was resuspended in 400 uL ChIP buffer containing protease and phosphatase inhibitors and 10 uL of diluted chromatin mixture was removed and placed into a new tube, which represents 2% input sample from each ChIP reaction, and mixed with 150 uL ChIP elution buffer.

For each immunoprecipitation, 500 uL of diluted chromatin was mixed with 4 ug of immunoprecipitating antibody and the samples were incubated at 4C rocking overnight in the presence of 30 uL ChIP-grade Protein G magnetic beads. Following immunoprecipitation, the protein G beads were collected by magnetic separation and washed 3 times in 1 mL low salt wash followed by 1 wash in high salt. Following, the final salt wash was aspirated and the magnetic beads were mixed with 150 uL ChIP elution buffer and placed at 65׈C for 30 minutes. Post elution, the input sample and all eluted immunoprecipitations were decross-linked via the addition of 6 uL 5M NaCL and 2 uL Proteinase K and incubated at 65׈C for 2 hours. Following, eluted DNA was purified using DNA spin columns.

At this stage, processed chromatin was analyzed via PCR utilizing the following program and primers:

94 ׈C - 3minutes

25 cycles (94 ׈C -20 seconds, 61 ׈C -30 seconds, 72 ׈C -30 seconds)

10 ׈C hold

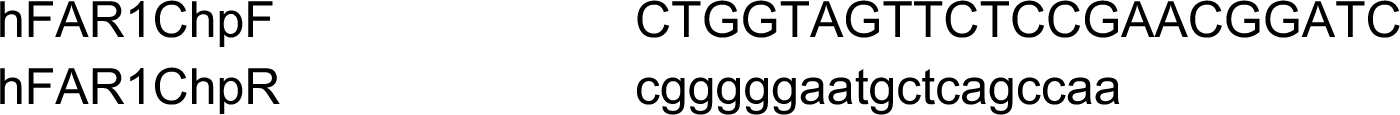

### Mice

The *Atg7^fl^*^/fl^ (JAX stock #034429), Egln1*^fl^*^/fl^(JAX stock #009672), BecnF121A (JAX stock #033360), *CAG-RFP-EGFP-LC3* (JAX stock #027139), Rosa26 HIF2 DPA (JAX stock #009674), Nestin-Cre (JAX stock #003771), HSA-Cre79 (Acta-Cre, JAX stock #006149), and aMHC-Cre (JAX stock #011038) mice were imported from Jackson Laboratory and were previously described. The Gnpat^fl/fl^ were a kind gift from Joe Hacia (USC) and were imported from the Jackson Laboratory. The Far1^fl/fl^ were purchased from GemPharmatech. Experimental protocols were approved by the Mass General Brigham Institutional Animal Care and Use Committee. Animals were housed at 24°C in a 12-hour light/12-hour dark cycle where food and water were accessible ad libitum. All mouse experiments were performed on littermate male and female mice aged 6-15 weeks. Mouse experiments involving Egln1*^fl^*^/fl^, Gnpat^fl/fl^, *Atg7^fl^*^/fl^, Far1^fl/fl^, and Rosa26 HIF2 DPA mice utilized Cre expressing wild-type mice to control for possible Cre toxicity.

For experiments involving mouse fasting, mice to be fasted were moved into a clean cage with new bedding with free access to water and deprived of food for either 24 or 48 hours prior to euthanasia.

For experiments involving hypoxia, mice housed at hypoxia were placed in a hypoxia chamber at 10% O_2_ for 7-days and food, water, and bedding was refreshed every other day prior to euthanasia.

### Isolation and Culture of Primary Mouse Neonatal Cardiomyocytes

Neonatal mice were sacrificed within the first 24 hours after birth. Beating hearts were removed and placed on ice in Hanks Balanced Salt Solution (HBSS) supplemented with sodium bicarbonate (1.6 mM). The great vessels were dissected away, and the hearts were minced with a razor blade. Minced hearts (∼40 hearts) were placed in 15 mL conical tubes containing HBSS with trypsin (1 mg/mL) for 2 hours at 4 ׈C. The tubes were then centrifuged at 15000 x g for 10 minutes at 4 ׈C. The supernatant was collected, and the minced hearts were subjected to 5 digestion steps (3 minutes each) with Collagenase Type II (125 U/mL) in HBSS at 37 ׈C. After each 3 minutes digestion, the supernatant was removed and diluted in isolation media (low glucose DMEM + 15% FBS supplemented with Pen/Strep). The cells were filtered (100 mm) and subjected to a Percoll gradient centrifugation (2100 x g for 30 minutes). The pellet was then resuspended in isolation media and cultured overnight on fibronectin coated 10-cm plates.

### Immunoprecipitations

For each immunoprecipitation, cells were equilibrated to serum-free DMEM 2 hours prior to immunoprecipitation. Cells were rinsed once with ice-cold PBS and lysed immediately with 1 mL Triton Lysis Buffer [1% Triton-X, 10 mM β-glycerol phosphate, 10 mM pyrophosphate, 40 mM HEPES pH 7.4, 2.5 mM MgCl_2_; and 1 tablet of EDTA-free protease inhibitor (Roche) per 25 ml lysis buffer]. The cell lysates were clarified by centrifugation at 17,000 x g in an Eppendorf microcentrifuge for 10 minutes at 4 °C. For anti-Flag immunoprecipitations, 30 ul of a 50% slurry of the FLAG affinity gel (Sigma) was then added to the cleared cell lysates and incubated with rotation overnight at 4 °C. For anti-mNG immunoprecipitation, 10 μL of mNeonGreen-Trap magnetic agarose (ChromoTek) was added to the cleared cell lysates and incubated with rotation overnight at 4 °C. For anti-RFP immunoprecipitations, 10 μL of RFP-Trap magnetic agarose (ChromoTek) was added to the cleared cell lysates and incubated with rotation overnight at 4 °C. For immunoprecipitations from tissues, tissues were lysed in 1 mL Triton Lysis Buffer by mechanical disruption using a Qiagen TissueLyser II on the highest setting for 10 minutes. The cell lysates were clarified by centrifugation at 17,000 x g in an Eppendorf microcentrifuge for 10 minutes at 4 °C.

Following immunoprecipitation, the beads were washed three times with Triton Lysis Buffer containing 150 mM NaCl. Immunoprecipitated proteins were denatured and eluted by boiling the beads in 50 μl sample buffer for 5 minutes, resolved by 4-20% or 4-12% SDS–PAGE and analyzed by immunoblotting.

### Protein Purification

For isolation of FAR1 protein, 293FT cells stably expressing the indicated protein with a N-terminal tandem GST FLAG affinity tag were grown to confluence in ∼10 confluent 15-cm plates, washed once room temperature PBS, scraped into 50 mL conical tubes, and pelleted in a clinical centrifuge at 4 ׈C. The pellets were gently resuspended in ice cold PBS and repelleted. The supernatant was then aspirated, and the remaining isolated cell pellets were suspended in ∼20 ml Buffer A (20 mM Hepes pH 7.4, 150 mM NaCl, 2 mM DTT) containing protease inhibitors. The cells were then disrupted by douncing 30 times on ice with pre-chilled douncers. Unbroken cells and the nuclear fraction were removed by centrifugation at 10,000 x g for 20 mins at 4 ׈C. The supernatant was then collected and centrifuged at 150,000 x g for 90 mins at 4 ׈C to isolate the membrane fraction. The pelleted membrane fraction was then solubilized using 40 mL Buffer B (20 mM Hepes pH 7.4, 500 mM NaCl, 2 mM DTT, and 1% DDM) for at least 4 hrs with rotation at 4 ׈C. Following membrane fraction solubilization, the insoluble material was pelleted by centrifugation at 20,000 x g for 10 mins at 4 ׈C and the supernatant containing the soluble material was transferred to a new Eppendorf. At this stage, 50 ׈L FLAG M2-affinity beads (Sigma) (equilibrated to Buffer B) was then added followed by 3 hrs immunoprecipitation at 4 ׈C. The beads were then washed 3 times in 1 mL Buffer C (20 mM Hepes pH 7.4, 500 mM NaCl, 2 mM DTT, and 0.1% DDM). After each wash, the beads were pelleted at 1500 x g for 1 min and transferred to a new Eppendorf tube. Following washing, the bound proteins were eluted by incubation in Buffer C without NaCl containing 1 mg/mL FLAG peptide with rotation for 1 hr at 4 ׈C. The eluted protein was concentrated using Amicon centrifuge filters and ultracentrifuged for 30 mins to remove aggregates and snap frozen in glycerol unless used immediately.

### Liquid chromatography tandem mass spectrometry (LC-MS/MS) proteomics

#### Cell lysis and protein extraction

LC-MS/MS analyses of 3 samples per condition (5 conditions, 15 total samples) were performed at the Broad Institute of MIT and Harvard. Conditions were as followed: WT U2OS cells grown at 21% oxygen for 18 hours, WT U2OS cells grown at 21% oxygen and treated with 250 nM Torin1 for 18 hours, WT U2OS cells grown at 1% oxygen for 18 hours, ATG7 KO U2OS cells grown at 21% oxygen for 18 hours, and ATG7 KO U2OS cells grown at 1% oxygen for 18 hours. Cell pellets of 40 million cells were suspended in cold, freshly-prepared lysis buffer (5% Sodium Dodecyl Sulfate (ThermoFisher, Waltham, Massachusetts), 50 mM triethylammonium bicarbonate buffer (Sigma-Aldrich, St. Louis, Missouri), 50 mM magnesium chloride (Sigma-Aldrich, St. Louis, Missouri), 2 μg/ml Aprotinin (Sigma-Aldrich, St. Louis, Missouri), 10 μg/ml Leupeptin (Roche CustomBiotech, Indianapolis, Indiana), 1 mM phenylmethylsulfonyl fluoride in EtOH (Sigma-Aldrich, St. Louis, Missouri), 10 mM sodium fluoride, 1% phosphatase inhibitor cocktail 2 and 3 (Sigma-Aldrich, St. Louis, Missouri), 5 μL 25KU Benzonase Nuclease (Sigma-Aldrich, St. Louis, Missouri), and deacetylase inhibitors (10 mM Sodium Butyrate, 2 μM SAHA, and 10 mM nicotinamide), vortexed for 10 seconds, and incubated for 5 min on benchtop. Samples were then centrifuged for 10 minutes at 4°C and 18000 rcf to remove debris. Supernatant was removed, and protein concentrations were determined by BCA assay (ThermoFisher, Waltham, Massachusetts).

#### Protein digestion

Protein lysate concentrations were normalized, and protein was reduced with 5 mM dithiothreitol (DTT, Sigma-Aldrich) for 1 hour in a thermomixer set to 37°C and 1000 rpm. Protein was then alkylated with 10 mM iodoacetamide (IAA, Sigma-Aldrich) for 45 minutes in the dark in a thermomixer set to 25°C and 1000 rpm. Following this, samples were acidified with phosphoric acid for a final concentration of 1.2%. Samples were then loaded onto midi S-traps (Protifi, Fairport, New York) in a ratio of 1:6 with chilled S-trap buffer (90% methanol, 100 mM triethylammonium bicarbonate buffer). S-trap midi protocol was then followed: centrifuge for 1 minute at 4,000 rcf, wash with 3 mL of S-trap buffer and spin at 4,000 rcf (repeat 3x), move column to a clean 15 mL tube, add 350 μL of digestion buffer (1:40 enzyme:substrate ratio LysC endopeptidase (1 mAU/μL, Wako Chemicals, Richmond, Virginia), 1:40 enzyme:substrate ratio sequencing grade modified trypsin (Promega, Madison, Wisconsin)), spin once more, then collect digestion buffer that has flowed through the column and pipette it back on top of the column. Tubes were left on the benchtop overnight for a 16-hour digestion. Columns were then eluted sequentially with 500 µL each of 50 mM TEAB, 0.1% formic acid, and 500 µL 0.1% formic acid 50% acetonitrile, with 1-minute centrifugations at 4000 rcf between each elution step. Elution flowthroughs were all collected into the same tube for each sample, frozen, dried using a vacuum concentrator, brought up in 3 ml total volume with 0.1% FA 3% acetonitrile, and desalted using Sep-Pac C18 SPE cartridges (Waters, Milford, MA). Clean peptides were dried using a vacuum concentrator, and peptide concentrations were determined by BCA assay. Following BCA assay, 50 μg aliquots of each sample were created for subsequent tandem mass tag (TMT) labeling step (see below).

#### Tandem mass tag (TMT) labeling of peptides

50 μg aliquots of digested lysate samples were resuspended in 50 mM HEPES pH 8.5 to a final concentration of 1 μg/μL for labeling. Experimental samples were randomized across 15 TMT channels (126C to 134N). 400 μg of each labeling reagent was added to the samples and reacted for 1 hour in a thermomixer set to 25°C and 1000 rpm. A small aliquot was removed from each sample for labeling efficiency and mixing tests, and the remaining sample was snap-frozen in liquid nitrogen and stored at -80°C. Following QC checks, reactions were quenched with 5% hydroxylamine and samples from each multiplex were combined, dried using a vacuum concentrator, and desalted using Sep-Pac C18 SPE cartridges (Waters).

#### Offline bRP fractionation

Each combined multiplex sample was fractionated by high pH reversed phase separation using a 3.5 μm Agilent Zorbax 300 Extend-C18 column (2.1 mm ID x 250 mm length). Samples were resuspended in mobile phase A (5 mM ammonium formate, pH 10, in 2% acetonitrile), centrifuged to remove debris, loaded onto the column and eluted off the column for 96 minutes at a flow rate of 200 μl/minute using mobile phase B (5 mM ammonium formate, pH 10, in 90% acetonitrile) with the following gradient (time(min): %B): 0:0; 7:0; 13:16; 73:40; 77:44; 82:60; 96:60. A total of 96 fractions were collected and concatenated down to 25 fractions by combining non-sequential fractions. These fractions were frozen in liquid nitrogen, dried using a vacuum concentrator, and stored at -80°C until ready for LC-MS/MS analysis.

#### LC-MS/MS analysis of samples

One microgram of each proteome fraction was analyzed on a QE-HFX mass spectrometer (Thermo Fisher Scientific) coupled to an Easy-nLC 1200 LC system (Thermo Fisher Scientific). Samples were separated using 0.1% Formic acid/3% Acetonitrile as buffer A and 0.1% Formic acid /90% Acetonitrile as buffer B on a 27cm 75um ID picofrit column packed in-house with Reprosil C18-AQ 1.9 mm beads (Dr Maisch GmbH) with a 110 min gradient consisting of 2-6% B in 1 min, 6-20% B in 62 min, 20-30% B for 22 min, 30-60% B in 9 min, 60-90% B for 1 min followed by a hold at 90% B for 5 min. The MS method consisted of a full MS scan from 350 to 1800 m/z at 60,000 resolution and an AGC target of 3e6 followed by MS2 scans collected at 45,000 resolution with an AGC target of 1e5 and maximum injection time of 105 ms. Dynamic exclusion was set to 15 seconds. The isolation window used for MS2 acquisition was 0.7 m/z and 20 most abundant precursor ions were fragmented with a normalized collision energy (NCE) of 29 optimized for TMT16 data collection.

#### Raw MS/MS data processing

The raw MS/MS data was searched on Spectrum Mill MS Proteomics Software (Broad Institute). MS2 spectra were extracted from RAW files and merged if originating from the same precursor, or within a retention time window of +/-60 s and *m/z* range of +/-1.4, followed by filtering for precursor mass range of 750-6000 Da and sequence tag length > 0. MS/MS search was performed using a human database downloaded from the UniProt on 08/05/2020 and common contaminants, with digestion enzyme conditions set to “Trypsin allow P”, <4 missed cleavages, fixed modifications (cysteine carbamidomethylation and TMT16 on N-term and lysine), and variable modifications (oxidized methionine, acetylation of the protein N-terminus, pyroglutamic acid on N-term Q, and pyro carbamidomethyl on N-term C). Matching criteria included a 40% minimum matched peak intensity and a precursor and product mass tolerance of +/-20 ppm. Peptide-level matches were validated if found to be below the 1.2% false discovery rate (FDR) threshold and within a precursor charge range of 2-6. A second round of validation was then performed for protein-level matches for proteome datasets, requiring a minimum protein score of 13 and protein level FDR of 0%.

Protein quantification was achieved by taking the ratio of TMT reporter ions for each sample over the median of all channels. TMT16 reporter ion intensities were corrected for isotopic impurities in the Spectrum Mill protein/peptide summary module using the afRICA correction method which implements determinant calculations according to Cramer’s Rule79 and correction factors obtained from the reagent manufacturer’s certificate of analysis for lot number VH310017.

#### Proteomics statistical analysis

After performing Median-MAD normalization, a moderated two-sample t-test was applied to the dataset to compare sample groups. Benjamini-Hochberg corrected p-value thresholds were used to assess differentially expressed proteins between experimental conditions.

#### Lipidomics

Analyses of polar and non-polar plasma lipids were conducted using an LC-MS system comprised of a Shimadzu Nexera X2 U-HPLC (Shimadzu Corp.) coupled to an Exactive Plus orbitrap mass spectrometer (Thermo Fisher Scientific). Lipids were extracted from human AC16 cardiomyocytes grown in 6-well plates seeded at a density of 2 x 10^5^ cells per well. At the time of extraction, the cells were washed twice in ice cold PBS (containing no Mg^2+^ or Ca^2+^) and followed by the addition of 800 μL HPLC grade isopropanol chilled to 4 ׈C prior to use. Cells were then collected by scraping into 1.5 mL Eppendorf tubes on ice and covered protected from light. Following, cell culture wells were rinsed with an additional 200 μL HPLC grade isopropanol and transferred into the eppendorff tube. To extract lipids from mouse hearts, flash frozen tissues (25-100 mg) were homogenized in HPLC grade isopropanol using a Tissue Lyzer (Qiagen). Lipid extracts were then placed at 4 ׈C for 1 hour, vortexed, and then centrifuged at 9,000 x g for 10 minutes at 4 ׈C. The supernatant was then collected and transferred into a clean eppendorff tube and stored at -80 ׈C until analysis.

At time of analysis,10 uL of supernatants were injected directly onto a 100 x 2.1 mm, 1.7 µm ACQUITY BEH C8 column (Waters). The column was eluted isocratically with 80% mobile phase A (95:5:0.1 vol/vol/vol 10mM ammonium acetate/methanol/formic acid) for 1 minute followed by a linear gradient to 80% mobile-phase B (99.9:0.1 vol/vol methanol/formic acid) over 2 minutes, a linear gradient to 100% mobile phase B over 7 minutes, then 3 minutes at 100% mobile-phase B. MS analyses were carried out using electrospray ionization in the positive ion mode using full scan analysis over 220–1100 m/z at 70,000 resolution and 3 Hz data acquisition rate. Other MS settings were: sheath gas 50, in source CID 5 eV, sweep gas 5, spray voltage 3 kV, capillary temperature 300°C, S-lens RF 60, heater temperature 300°C, microscans 1, automatic gain control target 1e6, and maximum ion time 100 ms. Raw data were processed using Progenesis QI (Nonlinear Dynamics) for peak detection and integration of both metabolites of known identity and unknowns. Lipid identities were determined based on comparison to reference plasma extracts and are denoted by total number of carbons in the lipid acyl chain(s) and total number of double bonds in the lipid acyl chain(s). Manual integration of XICs of a subset of identified metabolites was performed to augment results generated using Progenesis QI. Alkyl and alkenyl ether lipids are denoted in volcano plots and graphs found in Figure 6A-C, 7C, S12B, S15A-D, and S18C-E. Black dots indicate alkyl (ether) phosphatidylcholine (PC), dark red dots indicate alkenyl PC (plasmalogens), dark blue dots indicate ester linked PC species, red dots indicate alkyl (ether) phosphotidylethanolamine (PE), light blue dots indicate alkenyl PE (plasmalogens) species. Dots that are black and dark red indicate ether PC species that are determined as the alkyl or alkenyl form while dots that are red and light blue indicate ether PE species that are determined as either the alkyl or alkenyl form.

#### Determination of plasmalogen content in cells

Human AC16 cells were plated in 10 cm plates and grown to 80% confluency. At time of cell harvest, cells were washed twice in ice-cold PBS and isolated by cell scraping in 1 mL ice-cold PBS and transferred to a 1.5 mL Eppendorf tube. Following, cells were pelleted by centrifugation at 1000 x g for 1.5 minutes at room temperature and PBS was aspirated and cell pellets were snap frozen in liquid nitrogen and stored at -80C until further analysis. Cell pellets were homogenized in 200 µL buffer containing 0.1 M Tris and 0.25 M sucrose, pH 7.2 (HCl) by shaking with glass beads, protein concentration was determined by Bradford assay and 200 µL of aqueous homogenates were extracted twice with 500 µL of chloroform/methanol (2:1 v/v). The organic phases were combined and evaporated to dryness. Lipid extracts were dissolved in 100 µL of acetonitrile/ethanol (1:1 v/v) by shaking for 10 min at 37°C. Ten microliters of lipid extracts were then derivatized by mixing with either 40 µL dansylhydrazine (Sigma 03334) (0.45 mg/ml in acetonitrile/2M HCl (930:70 v/v))), which cleaves plasmalogens to the respective aldehydes and yields the corresponding dansylhydrazides, or with 40 µL of dansylhydrazine, which leaves the plasmalogen intact and derivatives free aldehydes only. The resulting mixture was centrifuged for 5 min and then injected into an Agilent 1260 Infinity II HPLC system equipped with a thermostatted autosampler, UV-Vis and fluorescence detectors, and a column thermostat. The dansylhydrazine derivatives eluting from 8 to 10 min were quantified by the fluorescence peak area (excitation 340 nm/emission 525 nm). As standard, octadecanal (Apollo Scientific OR54467) was dissolved in 2.5 mM acetonitrile/ethanol (1:1 v/v) and then diluted with acetonitrile to 20 µM. 10 µl of this solution were derivatized in parallel with the samples.

#### PEDS1 activity assay

To measure PEDS1 activity, human AC16 cells were plated in 10 cm plates and grown to 80% confluency. At time of cell harvest, cells were washed twice in ice-cold PBS and isolated by cell scraping in 1 mL ice-cold PBS and transferred to a 1.5 mL Eppendorf tube. Following, cells were pelleted by centrifugation at 1000 x g for 1.5 minutes at room temperature and PBS was aspirated and cell pellets were snap frozen in liquid nitrogen and stored at -80°C until further analysis. At time of PEDS1 activity analysis, cell pellets were resuspended in buffer containing 0.1 M Tris HCl and 0.25 M sucrose pH 7.2, protein concentration was determined by Bradford assay and all samples diluted to 1 mg/ml in 3.5 mg/ml bovine serum albumin in PBS. 0.1 mg/mL catalase (2000-5000 U/mg), 1 mM NADPH, 2 mM EDTA, and 2 µM pyrene-labeled lyso-substrate were added and assays incubated for 30 min at 37°C. Following incubation, the reaction mixture was divided into two equal portions and the reaction was stopped by addition of acetonitrile/2 M HCl (895:105 v/v) to one aliquot to cleave the vinyl ether bond or by the addition of acetonile/2M acetic acid (895:105 v/v) to the other aliquot as a control. The mixtures were incubated for 30 min at 37°C to complete the cleavage reaction. Following incubation, the samples were centrifuged at 20,000 x g for 5 min and 10 µL of reaction was injected into an Agilent 1260 Infinity II HPLC system equipped with a thermostatted autosampler, UV-Vis and fluorescence detectors, and a column thermostat. At a flow rate of 1 ml/min and an injection volume of 10 μl, a Zorbax Eclipse XDB-C8 4.6 × 50 mm, 3.5 μm particle size column (Agilent Technologies, Vienna, Austria) was eluted with 10 mM potassium phosphate buffer (pH 6.0) containing 79% (v/v) methanol for 3 min, followed by a linear gradient to 100% methanol at 10 min. Methanol (100%) was held until 15 min, and the column was then equilibrated to starting elution buffer until 17 min. Pyrene-labeled compounds were by fluorescence (excitation 340 nm, emission 410 nm). Plasmalogen-derived pyrenedecanal liberated upon HCl treatment was monitored by comparison to an external synthetic pyrenedecanal standard and the amount of pyrenedecanal formed from plasmalogen cleavage was quantified.

#### Immunofluorescence Staining

For immunofluorescence assays, mouse neonatal cardiomyocytes or human AC16 cardiomyocytes were seeded and cultured on coverslips in 6-well plates under the desired conditions. To fix the cells, the media was aspirated, and the cells were washed twice with cold 1X PBS and fixed using 4% paraformaldehyde in 1X PBS or ice cold 100% methanol for 15 minutes at room temperature. Paraformaldehyde-fixed cells were then permeabilized by incubation in 0.05% TritonX-100 in 1X PBS for 5 minutes at room temperature. The coverslips were blocked with Odyssey Blocking Buffer (Li-Cor) for 1 hour at room temperature. The cells were subsequently stained sequentially with primary and secondary antibody diluted in Odyssey blocking buffer for 1 hour each at room temperature, being washed thrice after both primary and secondary antibody staining. The cells were then counterstained with DAPI and mounted onto glass slides using mounting medium and imaged within 48 hours. Anti-mouse Alexa Fluor 488 and Anti-rabbit Alexa Fluor 568 secondary antibodies (Invitrogen) were used at 1:400 dilution.

#### Confocal Microscopy

Live cell and fixed cell confocal microscopy were performed using a Zeiss LSM 900 confocal microscope equipped using a 63 x Plan-Apochromat oil objective, NA 1.4. For live cell imaging experiments, 2 x 10^5^ cells were plated onto 35 mm-glass bottom dishes (No. 1.5, 14 mm glass diameter, MatTek) then incubated in phenol-red free medium (FluoroBriteDMEM, Thermo Fisher) containing 10% fetal bovine serum for 24 hours prior to drug treatment, oxygen deprivation, or nutrient deprivation. At time of imaging, cells were imaged in FluoroBrite DMEM without 10% fetal bovine serum. Series optical sections were collected with a step-size of 0.22 microns. Z series were displayed as maximum z-projections, and gamma, brightness, and contrast were adjusted for each image equally using FiJi software. At least 10 image frames per condition were analyzed without exclusion.

#### Image Analysis

For quantification of co-localization, at least 10 non-overlapping images were acquired from each coverslip for immunofluorescence imaging or from each glass bottom dish for live cell imaging experiments. Raw, unprocessed images were imported into FiJi and converted to 8-bit images, and images of individual channels were thresholded independently to exclude background and converted to binary masks. Co-localization between lysosomes or LC3B-positive autophagosomes and marker of interest was determined using the “AND” function of the image calculator.

For quantification of peroxisome number or WIPI-2 puncta, raw, unprocessed images were imported into FiJi and converted to 8-bit images. Each image was individually thresholded and converted to binary masks. The “Analyze Particles” function was used to identify and count PEX14-positive or WIPI-2-positive particles in the resulting mask and then normalized to number of cells as marked by DAPI-positive nuclei in the image field. Data are plotted as peroxisome number or WIPI-2 puncta normalized to wild-type or untreated condition per cell analyzed.

#### In vivo Ischemia Reperfusion Injury

8–12-week-old mice were anesthetized, intubated, and ventilated. A left thoractotomy was performed and the left anterior descending (LAD) artery was ligated with a 6.0 silk suture. Ischemia was confirmed by myocardial blanching and electrocardiographic (ECG) evidence of injury. The LAD ligature was released 30 minutes later for reperfusion. Overall mouse survival was 70–80% at 24 hours. Mice were terminally anesthetized 24 hours after ischemia with ketamine/xylazine followed by cervical dislocation. Their hearts were harvested, and the ventricles were sectioned from apex to base in 2 mm sections. Sections were incubated in 2% (wt/vol) triphenyltetrazolium (TTC, Sigma) in phosphate-buffered saline for 30 minutes at 37°C followed by 1 hour fixation in formaldehyde. Fraction MI was calculated as the infarcted area divided by the AAR.

#### Langendorff model of Global Ischemia Reperfusion Injury

Mice were heparinized and anesthetized. A thoracotomy was performed, and the heart was rapidly excised. The aorta was cannulated, and the heart was perfused retrograde at a constant flow of ∼3 ml/min with perfusate buffer (124 mM NaCl, 25 mM NaHCO3, 11 mM dextrose, 4 mM KCl, 1 mM MgSO4, 1 mM KH2PO4, 2.8 mM CaCl2). A perfusate filled balloon was inserted into the left ventricle via the mitral valve and connected to a pressure transducer for continuous recording of the left ventricular chamber pressure. Langendorff-perfused hearts were equilibrated by being perfused for 15 minutes prior to induction of global ischemia for 15 minutes and reperfusion for at least 30 minutes. Test compounds stock solutions were diluted to a 20x final concentration in perfusate buffer and then combined to the perfusate line via a syringe pump set to 1/20th the total flow rate.

#### Statistical Analysis

GraphPad Prism version 10 was used for statistical analysis included in the main and supplementary figures. For all experiments, statistical significance was calculated using unpaired, two-tailed Student’s *t* test when comparing two groups or one-way ANOVA when comparing three or more groups. *P* values were considered statistically significant if the *P* value was <0.05. For all figures, * indicates *p* < 0.05 unless otherwise indicated. Error bars represent SD unless otherwise indicated. Data was evaluated for normality using the Shaprio-Wilk test and D’Agostino & Pearson Test. No statistical methods were used to pre-determine the sample size. No sample size calculation was performed for fluorescence microscopy experiments. Unless otherwise indicated, all proteomic and lipidomic measurements were performed on three independent biological replicates for each condition.

## Supplementary Figure Legends

**Supplementary Figure 1.**
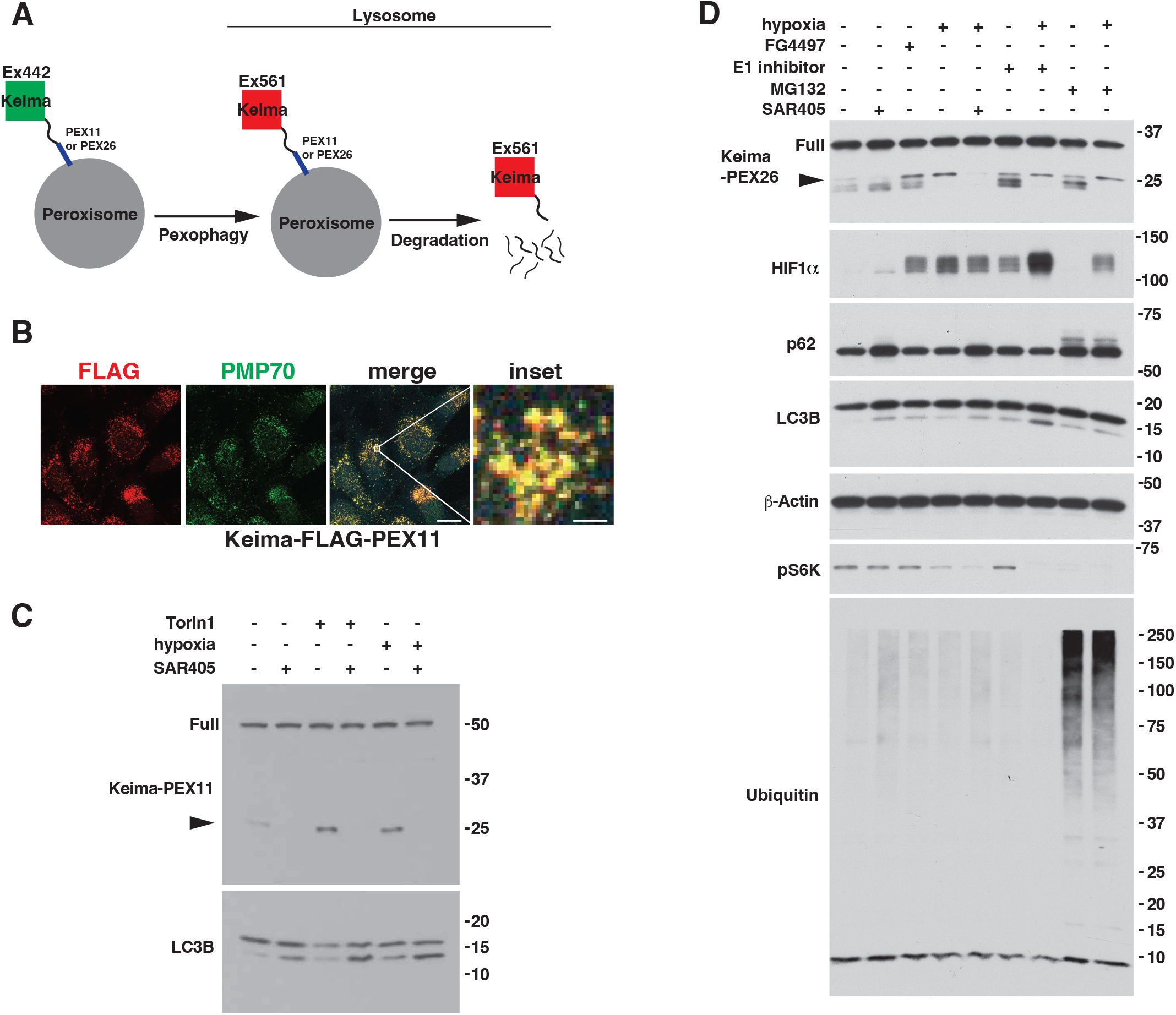
Validation of Peroxisome-Keima reporter system. **(A)** Schematic of Keima Peroxisome reporter system that allows for monitoring of lysosomal delivery of autophagy cargo. The acidic environment of the lysosome causes an increase in the ratio of 561 nm/488 nm excitation. Once the Keima-peroxisome fusion protein enters the lysosome, the Keima-fusion is processed to degrade the fusion protein, while the Keima (and any peptidic remnants left from the fusion partner) is resistant to the lysosomal proteases and maintains its fluorescence in the lysosome. **(B)** Immunofluorescence analysis of human AC16 cardiomyocytes stably expressing Keima-FLAG-PEX11. Scale bar indicates 10 um, inset scale bar indicates 1 um. **(C)** Immunoblot analysis of human U2OS osteosarcoma cells stably expressing Keima-PEX11. Where indicated, cells were grown at 0.1% O2 (hypoxia), treated with 250 nM Torin1, 1 uM SAR405 or normoxia and treated with DMSO for 18 hours. Full indicates full-length Keima-fusion protein and arrow indicates liberated Keima protein. **(D)** Immunoblot analysis of human U2OS osteosarcoma cells stably expressing Keima-PEX26. Where indicated, cells were grown at 0.1% O2 (hypoxia), treated with 100 uM FG-4497, 1 uM TAK-243 (E1 inhibitor), 1 uM MG132, 1 uM SAR405, or normoxia and treated with DMSO for 18 hours. Full indicates full-length Keima-fusion protein and arrow indicates liberated Keima protein.

**Supplementary Figure 2.**
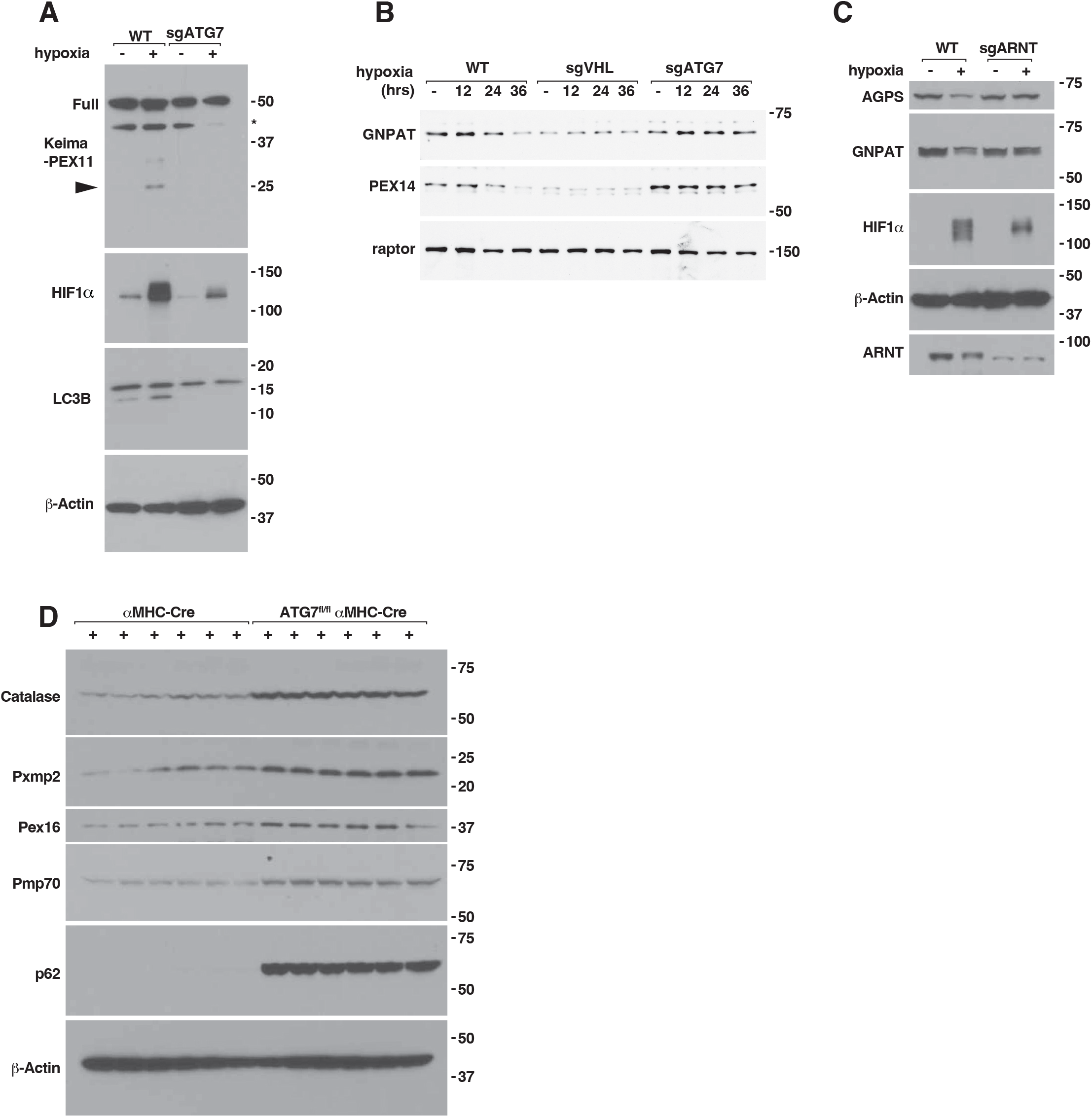
ATG7 is necessary for hypoxia-induced pexophagy. (A-C) Immunoblot analysis of wild-type (WT), sgATG7 (A-B), sgVHL (B), and sgARNT (C) human AC16 cardiomyocytes. In (A), cells are stably expressing Keima-PEX11 and grown at 0.1% O2 (hypoxia) or normoxia for 18 hours. *indicates non-specific band. In (B), cells were grown at 0.5% O2 (hypoxia) or normoxia for indicated time points (hours) prior to cell lysis. In (C), cells were grown at 0.1% O2 (hypoxia) or normoxia for 36 hours prior to cell lysis. **(D)** Immunoblot analysis of Atg7^+/+^-αMHC-Cre and Atg7^fl/fl^-αMHC-Cre mouse hearts.

**Supplementary Figure 3.**
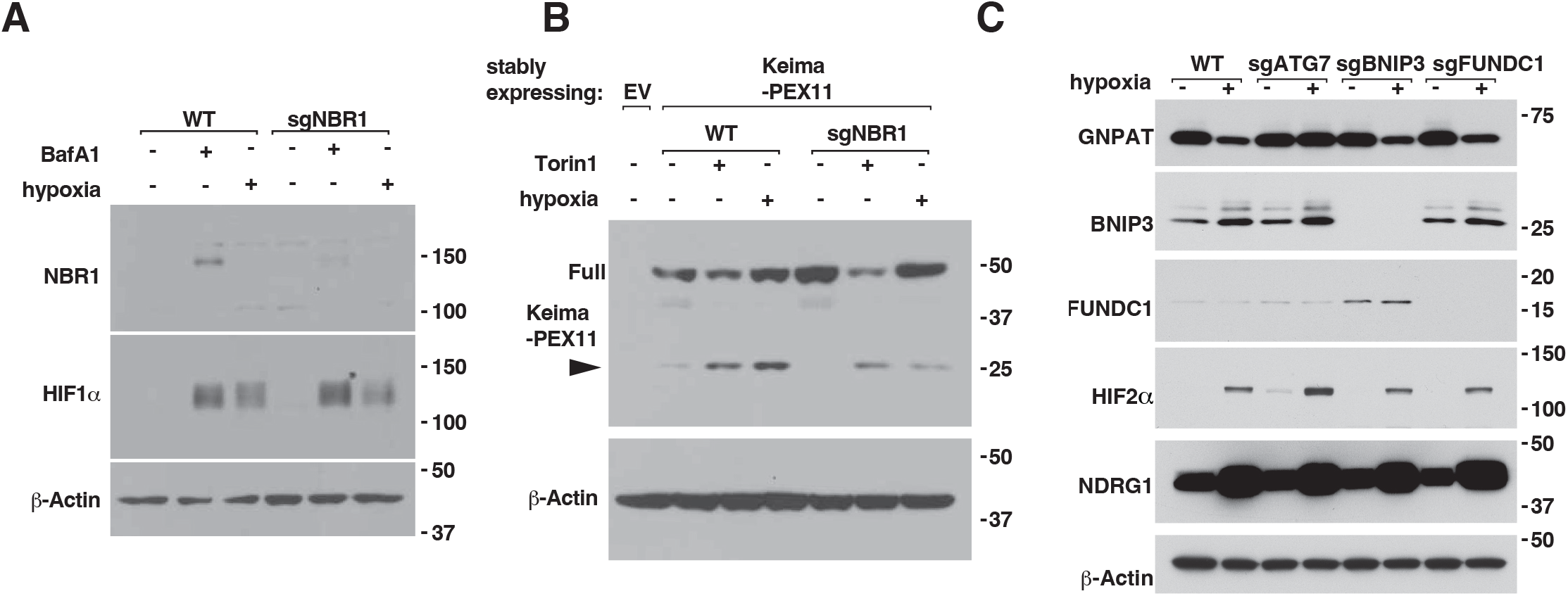
Loss of NBR1, BNIP3, or FUNDC1 does not block hypoxia -induced pexophagy in human AC16 cardiomyocytes. (A-B) Immunoblot analysis of wild-type (WT) and sgNBR1 human AC16 cardiomyocytes. In (A), cells were treated with 500 nM Bafilomycin A1 (BafA1), grown at 0.1% O2 (hypoxia), or normoxia and treated with DMSO for 18 hours. In (B), cells were stably expressing Keima-PEX11 or empty vector (EV). Where indicated, cells were treated with 250 nM Torin1, grown at 0.1% O2 (hypoxia), or normoxia and treated with DMSO for 18 hours. Full indicates full-length Keima-fusion protein and arrow indicates liberated Keima protein. **(C)** Immunoblot analysis of wild-type (WT), sgATG7, sgBNIP3, and sgFUNDC1 human AC16 cardiomyocytes. Where indicated, cells were grown at 0.1% O2 (hypoxia) or normoxia for 36 hours prior to cell lysis.

**Supplementary Figure 4.**
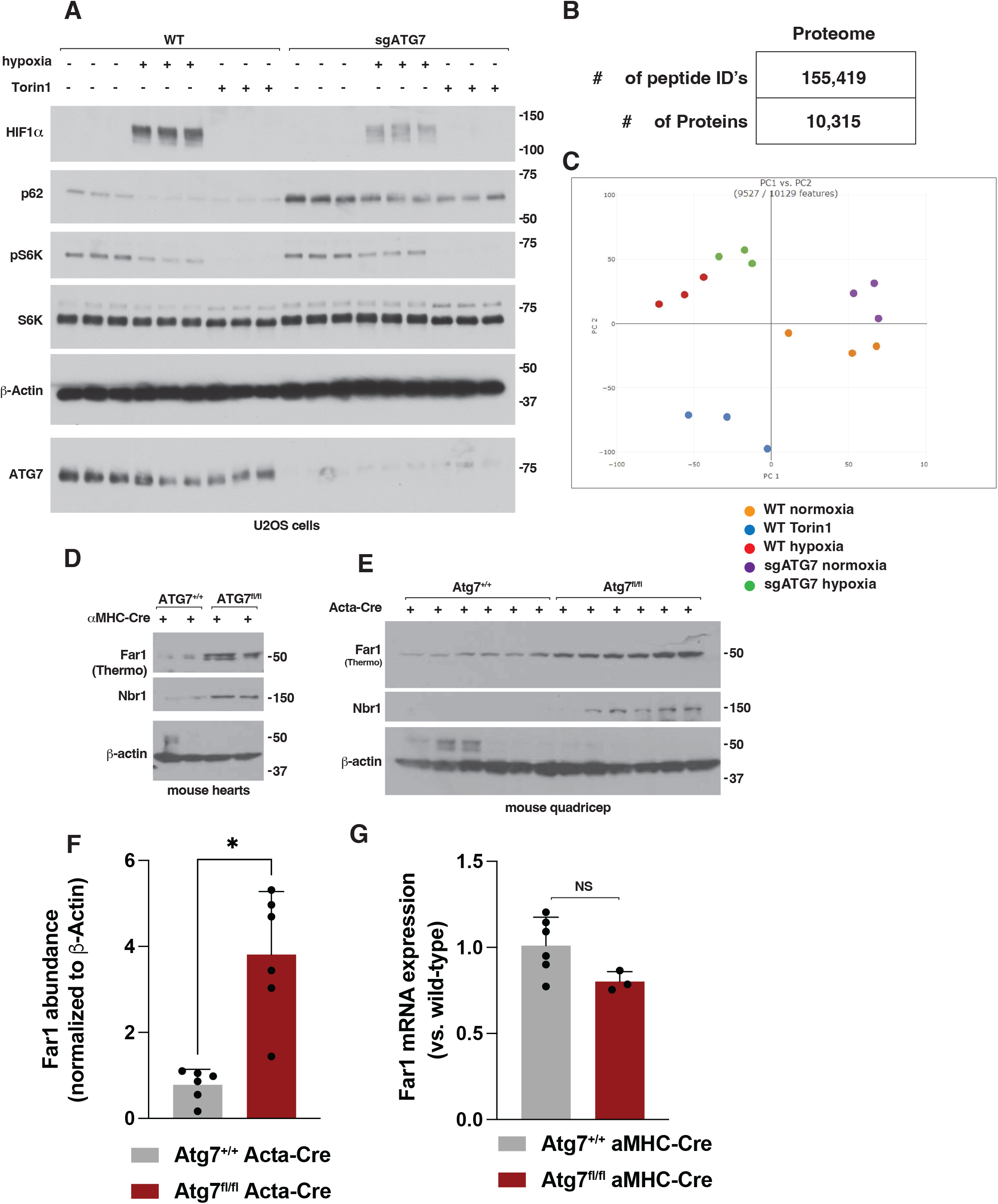
Identification of FAR1 via quantitative proteomics. **(A)** Immunoblot analysis of wild-type and sgATG7 human U2OS osteosarcoma cells. Where indicated, cells were grown at 0.1% O_2_ (hypoxia), treated with Torin1 (250 nM), or treated with DMSO and grown at normoxia for 18 hours prior to cell lysis. **(B)** U2OS proteome analysis of total peptides and total proteins identified. **(C)** Principal component analysis of tandem mass tagging (TMT) analysis of wild-type and sgATG7 U2OS cells grown under 0.1% O_2_ (hypoxia), treated with Torin1 (250 nM) or treated with DMSO and grown at normoxia for 18 hours prior to cell lysis. **(D)** Immunoblot analysis of Atg7^+/+^-αMHC-Cre and Atg7^fl/fl^-αMHC-Cre mouse hearts. Far1 immunoblot indicates which Far1 antibody was used to detect Far1. **(E)** Immunoblot analysis of Atg7^+/+^-Acta-Cre and Atg7^fl/fl^ Acta-Cre mouse quadricep muscles. **(F)** Densitometry analysis of Far1 abundance shown in (E). N=6. *indicates P<0.05. **(G)** Far1 mRNA analysis in mouse hearts isolated from Atg7^+/+^-αMHC-Cre and Atg7^fl/fl^-αMHC-Cre mice. N>3, NS=non-significant.

**Supplementary Figure 5.**
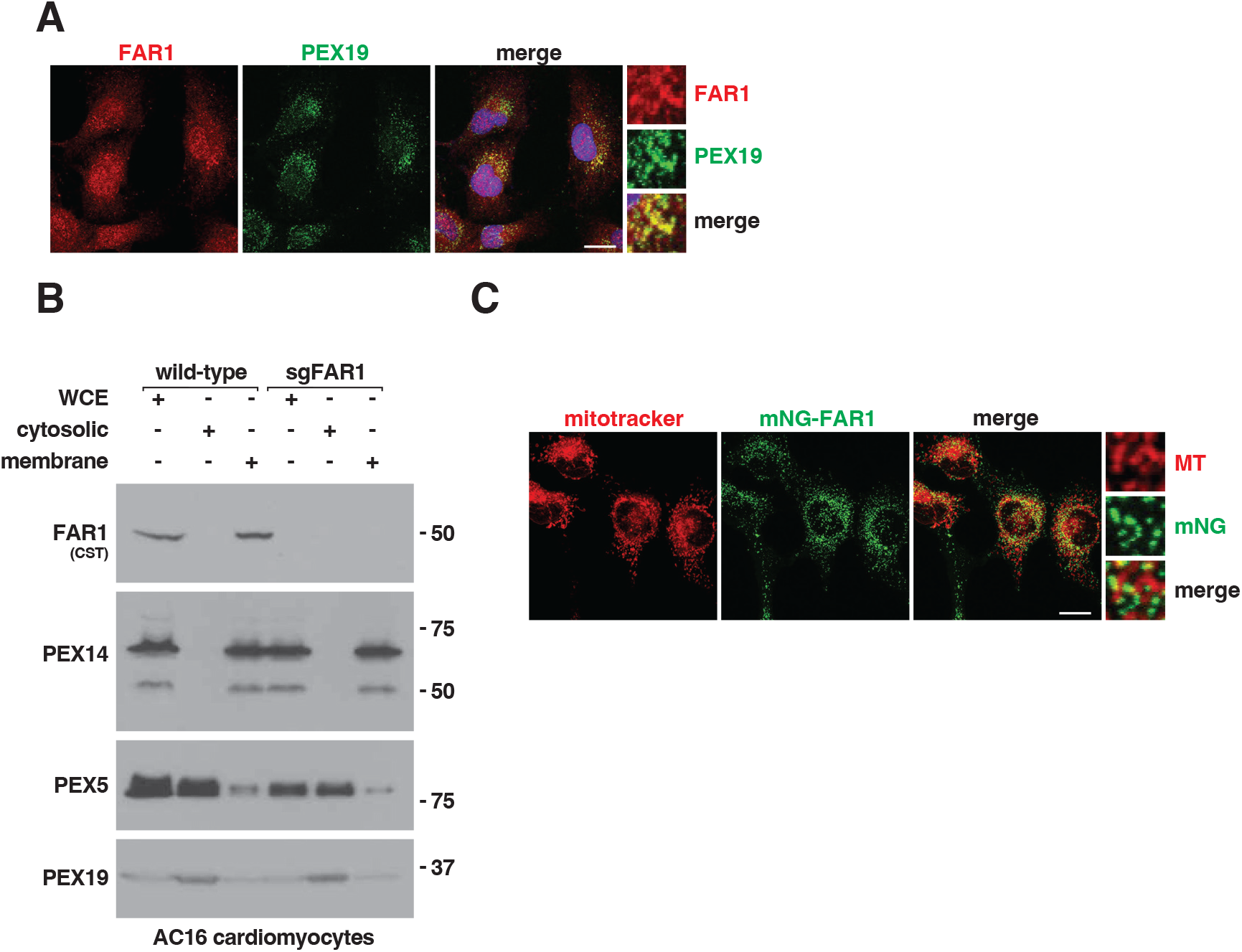
FAR1 localization at the peroxisome. **(A)** Confocal microscopy analysis of mouse neonatal cardiomyocytes. Scale bar indicates 10 um. **(B)** Immunoblot analysis of subcellular fractionation in wild-type and sgFAR1 human AC16 cardiomyocytes. WCE, whole cell extract. Cytosolic, cytosolic fraction. Membrane, membrane fraction. Far1 immunoblot indicates which Far1 antibody was used to detect Far1. **(C)** Confocal microscopy analysis of human AC16 cardiomyocytes stably expressing mNeonGreen-Far1 (mNG-Far1) and labeled with 100 nM MitoTracker Deep Red. Scale bar indicates 10 um.

**Supplementary Figure 6.**
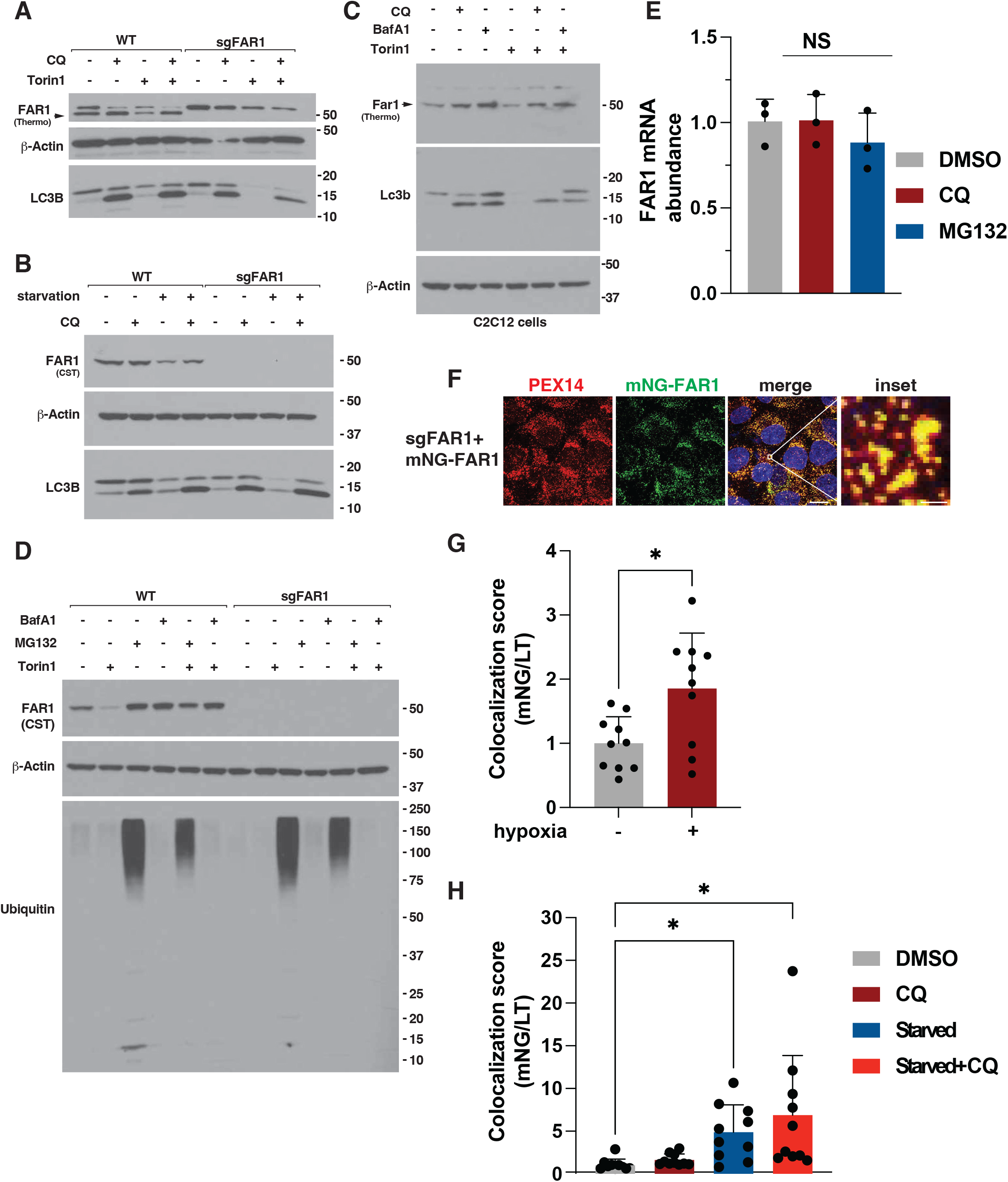
FAR1 is degraded via the lysosome. **(A)** Immunoblot analysis of wild-type and sgFAR1 mouse neonatal cardiomyocytes. Where indicated, cells were treated with 250 nM Torin1, 30 uM Chloroquine (CQ), or DMSO for 18 hours prior to cell lysis. Arrow indicates FAR1. Far1 immunoblot indicates which Far1 antibody was used to detect Far1. **(B)** Immunoblot analysis of wild-type and sgFAR1 human 293FT cells. Where indicated, cells were deprived of nutrients (starvation), treated with 30 uM Chloroquine (CQ), or DMSO for 18 hours prior to cell lysis. Far1 immunoblot indicates which Far1 antibody was used to detect Far1. **(C)** Immunoblot analysis of mouse C2C12 myoblasts. Where indicated, cells were treated with 250 nM Torin1, 500 nM Bafilomycin A1 (BafA1), 30 uM Chloroquine (CQ), or DMSO for 18 hours prior to cell lysis. Arrow indicates Far1. Far1 immunoblot indicates which Far1 antibody was used to detect Far1. **(D)** Immunoblot analysis of wild-type and sgFAR1 human 293FT cells. Where indicated, cells were treated with 250 nM Torin1, 500 nM Bafilomycin A1 (BafA1), 1 uM MG132, or DMSO for 18 hours prior to cell lysis. Far1 immunoblot indicates which Far1 antibody was used to detect Far1. **(E)** *FAR1* mRNA analysis in human AC16 cardiomyocytes. Where indicated, cells were treated with 30 uM Chloroquine (CQ), 1 uM MG132, or DMSO for 18 hours prior to RNA extraction. N=3, NS=non-significant. **(F)** Confocal microscopy analysis of sgFAR1 293FT cells stably expressing mNG-FAR1. Scale bar indicates 10 um, inset scale bar indicates 1 um. **(G)** Image analysis of sgFAR1 human 293FT cells stably expressing mNG-FAR1 and labeled with lysotracker (LT) shown in Figure 2F. Where indicated, cells were grown at 0.1% O_2_ (hypoxia) or normoxia prior to imaging. N=10, *P<0.05. **(H)** Image analysis of sgFAR1 human AC16 cells stably expressing mNG-FAR1 and labeled with lysotracker (LT) shown in Figure 2G. Where indicated, cells were starved of nutrients, treated with 30 uM Chloroquine (CQ), or DMSO for 18 hours prior to imaging. N=10, *P<0.05.

**Supplementary Figure 7.**
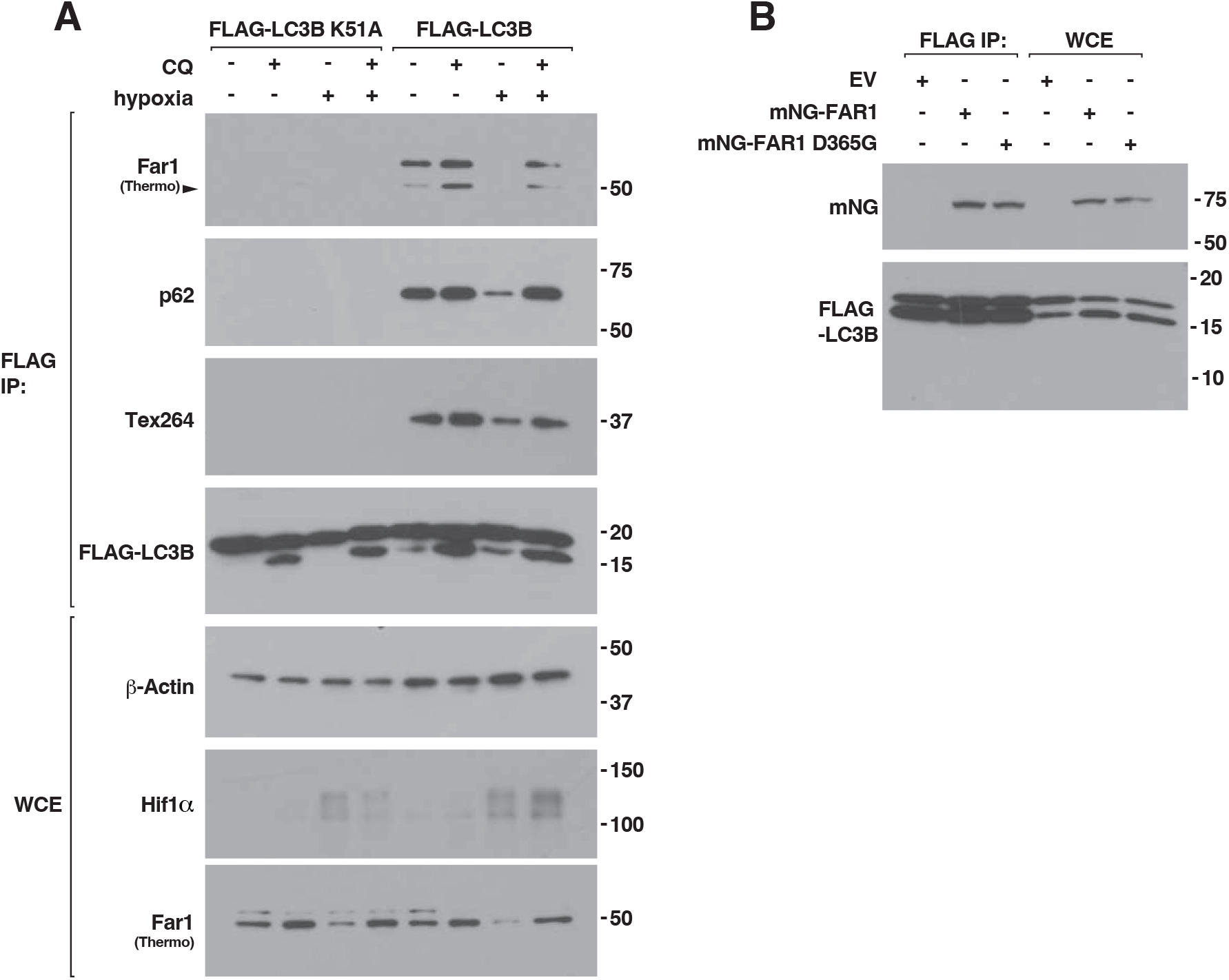
FAR1 binds LC3B. **(A)** Immunoblot analysis of anti-FLAG immunoprecipitates isolated from cardiomyocytes stably expressing FLAG-LC3B K51A or FLAG-LC3B. Where indicated, cells were treated with 30 uM Chloroquine (CQ), grown at 0.1% O_2_ (hypoxia), or treated with DMSO and grown at normoxia for 18 hours prior to cell lysis. Arrow indicates Far1. WCE, whole cell extract. Far1 immunoblot indicates which Far1 antibody was used to detect Far1. **(B)** Immunoblot analysis of anti-FLAG immunoprecipitates isolated from AC16 cells stably expressing FLAG-LC3B and mNG-FAR1, mNG-FAR1 D365G, or empty vector (EV). WCE, whole cell extract.

**Supplementary Figure 8.**
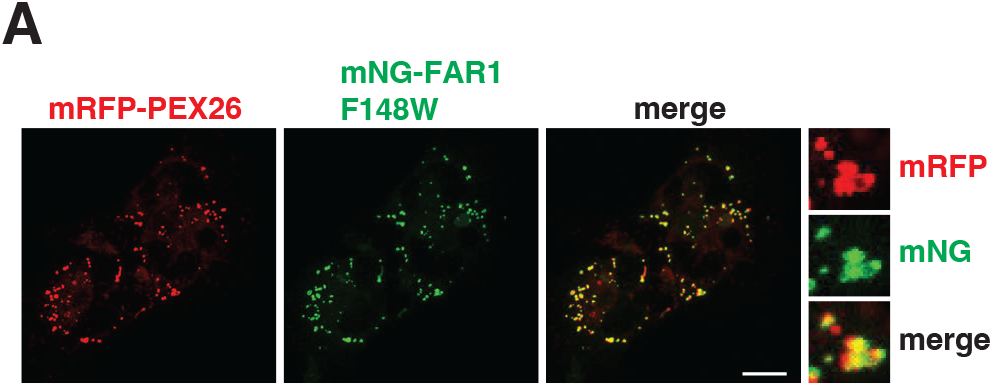
FAR1 F148W localizes to the peroxisome. **(A)** Confocal microscopy analysis of sgFAR1 human AC16 cardiomyocytes stably expressing mNeonGreen-Far1 F148W (mNG-Far1 F148W) and mRFP-PEX26. Scale bar indicates 10 um.

**Supplementary Figure 9.**
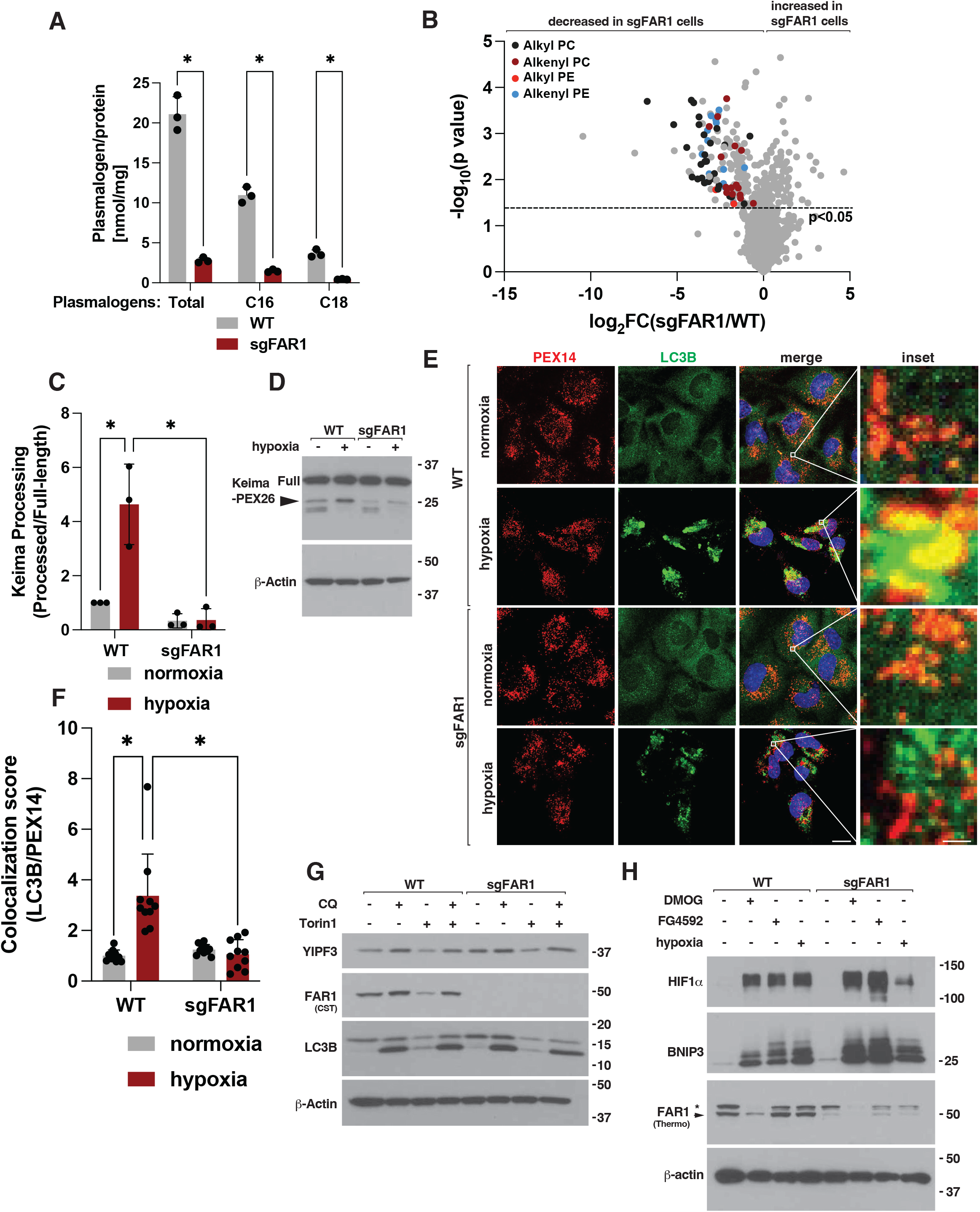
FAR1 is necessary for hypoxia induced peroxisome degradation. **(A)** Total-, C16-, and C18-plasmalogens (nmol/mg protein) in wild-type and sgFAR1 human AC16 cardiomyocytes. N=3, *indicates P<0.05. **(B)** Volcano plots comparing lipid intensities in wild-type and sgFAR1 AC16 cardiomyocytes. Black dots indicate alkyl (ether) phosphatidylcholine (PC), dark red dots indicate alkenyl PC (plasmalogens), red dots indicate alkyl (ether) phosphotidylethanolamine (PE), light blue dots indicate alkenyl PE (plasmalogens) species. Dotted line indicates P<0.05. **(C)** Densitometry analysis of Keima-processing assay as shown in wild-type and sgFAR1 human AC16 cardiomyocytes stably expressing Keima-PEX11. N=3, *indicates P<0.05. **(D)** Immunoblot analysis of wild-type and sgFAR1 human AC16 cardiomyocytes stably expressing Keima-PEX26. Where indicated, cells were grown at 0.1% O2 (hypoxia) or normoxia for 18 hours prior to cell lysis. Full indicates full-length Keima-fusion protein and arrow indicates liberated Keima protein. **(E-F)** Confocal microscopy (E) and image analysis (F) of wild-type (WT) and sgFAR1 human AC16 cardiomyocytes. Where indicated, cells were grown at 0.1% O2 (hypoxia) or normoxia for 18 hours prior to cell fixation. N=10, *P<0.05. Scale bar indicates 10 um, inset scale bar indicates 1 um. **(G)** Immunoblot analysis of wild-type (WT) and sgFAR1 human AC16 cardiomyocytes. Where indicated, cells were treated with 250 nM Torin1, 30 uM Chloroquine (CQ), or DMSO for 18 hours prior to cell lysis. Far1 immunoblot indicates which Far1 antibody was used to detect Far1. **(H)** Immunoblot analysis of wild-type (WT) and sgFAR1 human AC16 cardiomyocytes. Where indicated, cells were treated with 500 uM Dimethyloxalylglycine (DMOG), grown at 0.1% O2 (hypoxia), treated with 100 uM FG-4592, or normoxia and treated with DMSO for 8 hours prior to cell lysis. Arrow indicates FAR1 while * indicates non-specific band. Far1 immunoblot indicates which Far1 antibody was used to detect Far1.

**Supplementary Figure 10.**
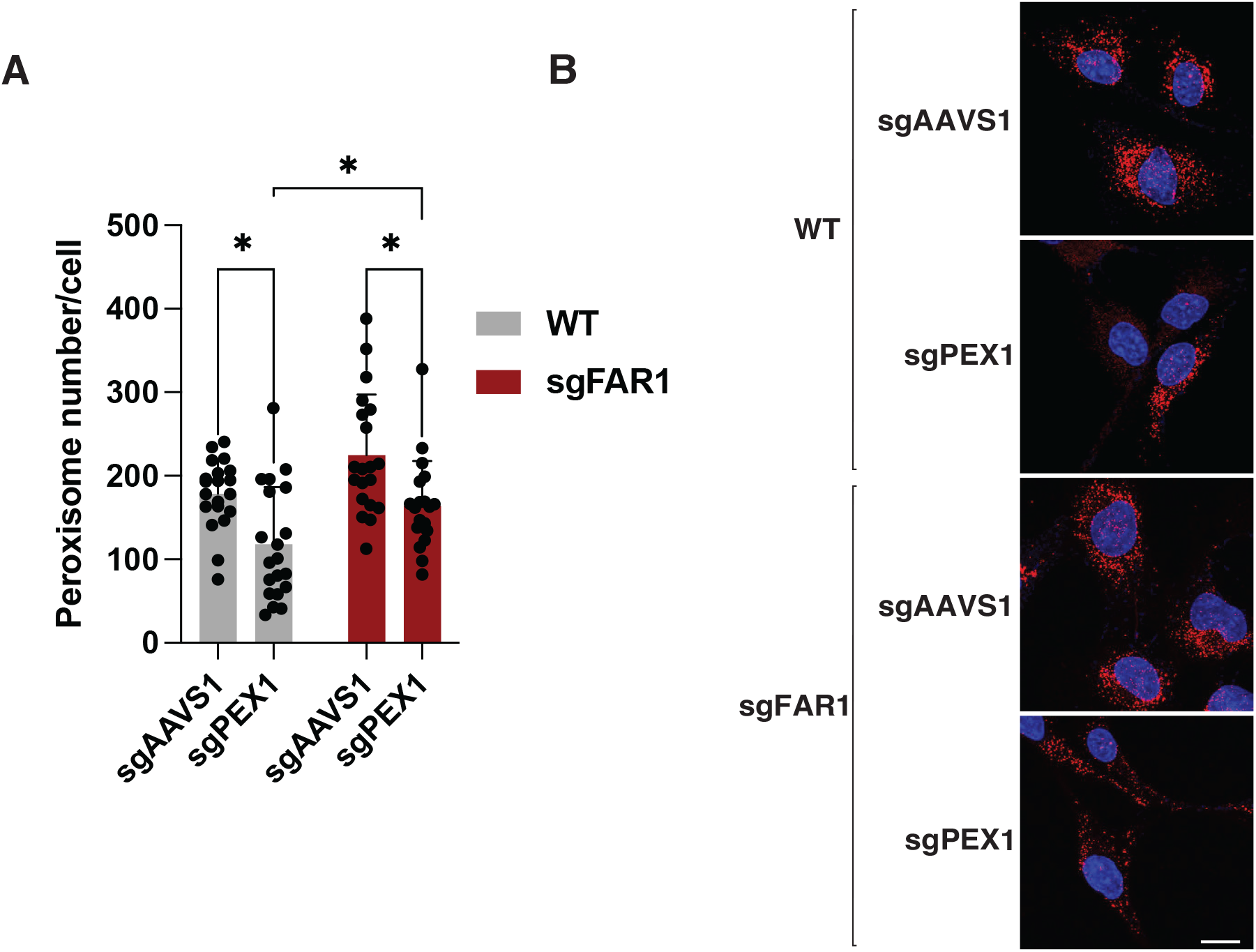
FAR1 and its role in PEX1-mediated pexophagy. (A-B) Confocal microscopy (A) and image analysis (B) of PEX14-positive peroxisomes in wild-type and sgFAR1 human AC16 cells stably expressing sgRNAs targeting sgAAVS1 (control) or sgPEX1. N>10, * indicates P<0.05.

**Supplementary Figure 11.**
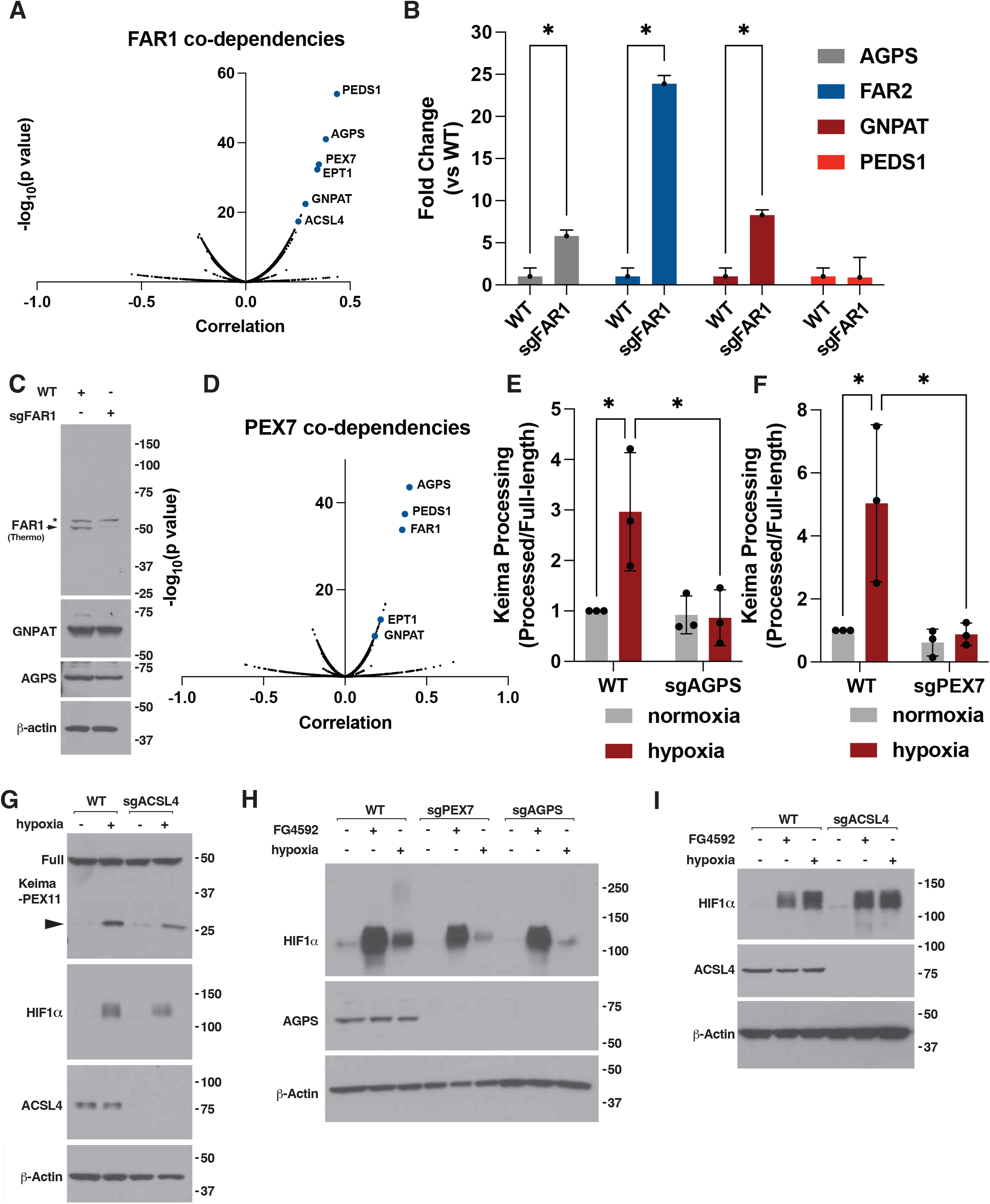
Far1 co-dependency mapping. **(A)** FAR1 co-dependency mapping using DepMap. **(B)** *AGPS*, *FAR2*, *GNPAT*, and *PEDS1* mRNA analysis, as determined by real-time polymerase chain reaction, in wild-type and sgFAR1 human AC16 cardiomyocytes. N=3, *P<0.05. **(C)** Immunoblot analysis of wild-type and sgFAR1 human AC16 cardiomyocytes. Arrow indicates FAR1 while * indicates non-specific band. Far1 immunoblot indicates which Far1 antibody was used to detect Far1. **(D)** PEX7 co-dependency mapping using DepMap. **(E-F)** Densitometry analysis of Keima-processing assay as shown in wild-type, sgAGPS (E) and sgPEX7 (F) human AC16 cardiomyocytes stably expressing Keima-PEX11. N=3, *indicates P<0.05. **(G)** Immunoblot analysis of wild-type and sgACSL4 human AC16 cells stably expressing Keima-PEX11. Full indicates full-length Keima-fusion protein and arrow indicates liberated Keima protein. **(H-I)** Immunoblot analysis of wild-type (WT), sgPEX7, sgAGPS (H), and sgACSL4 (I). Where indicated, cells were treated with 100 uM FG-4592, grown at 0.1% O2 (hypoxia), or treated with DMSO and grown at normoxia for 8 hours prior to cell lysis.

**Supplementary Figure 12.**
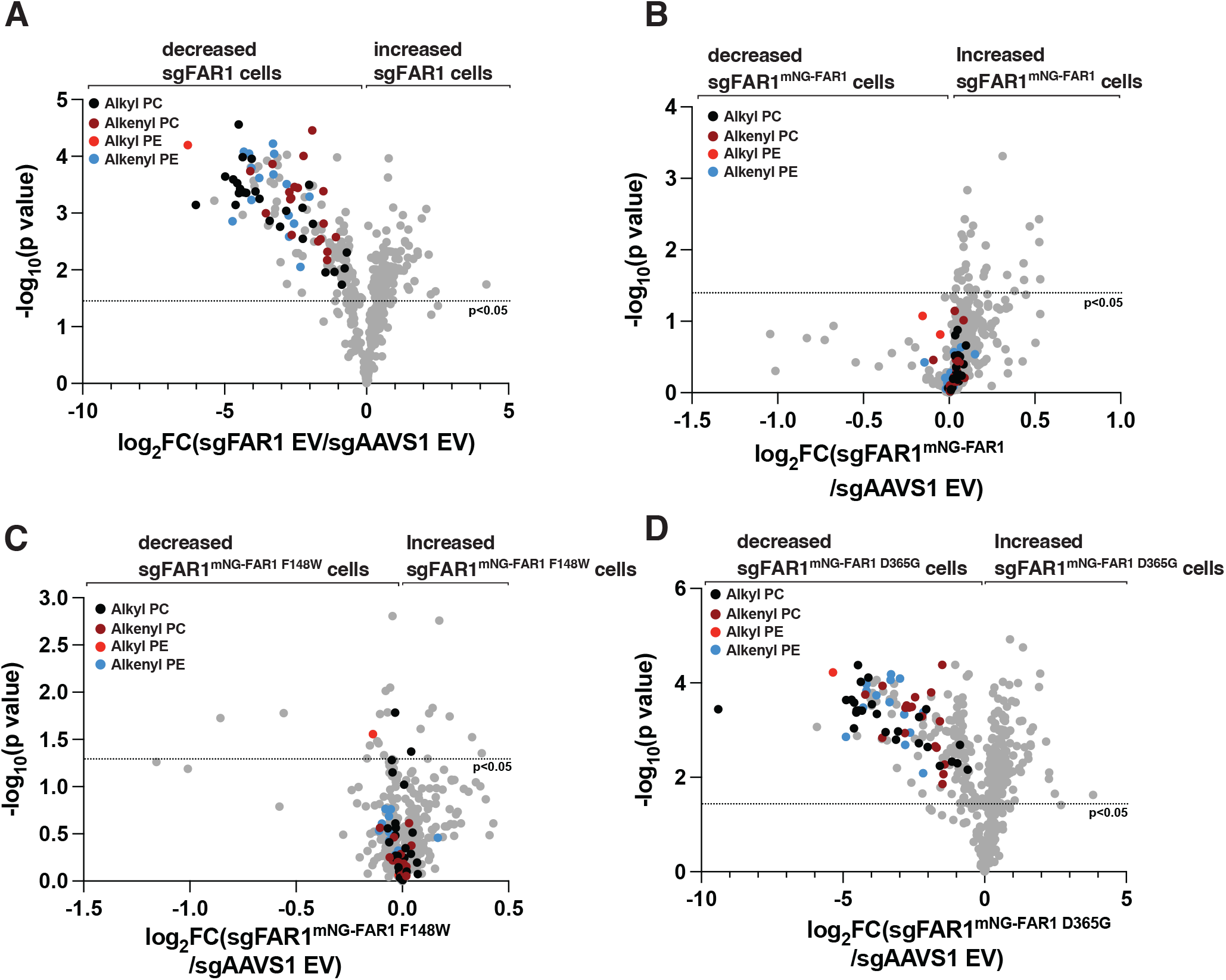
Lipidomic analysis sgFAR1 cells re-expressing FAR1 F148W or FAR1 D365G. (A-D) Volcano plots comparing lipid intensities in wild-type sgAAVS1 cells stably expressing empty vector (EV) and sgFAR1 293FT (A) cells stably expressing EV (A), mNG-FAR1 (B), mNG-FAR1 F148W (C), mNG-FAR1 D365G (D). All volcano plots are compared against sgAAVS1 293FT cells expressing EV. Black dots indicate alkyl (ether) phosphatidylcholine (PC), dark red dots indicate alkenyl PC (plasmalogens), red dots indicate alkyl (ether) phosphotidylethanolamine (PE), light blue dots indicate alkenyl PE (plasmalogens) species. Dotted line indicates P<0.05.

**Supplementary Figure 13.**
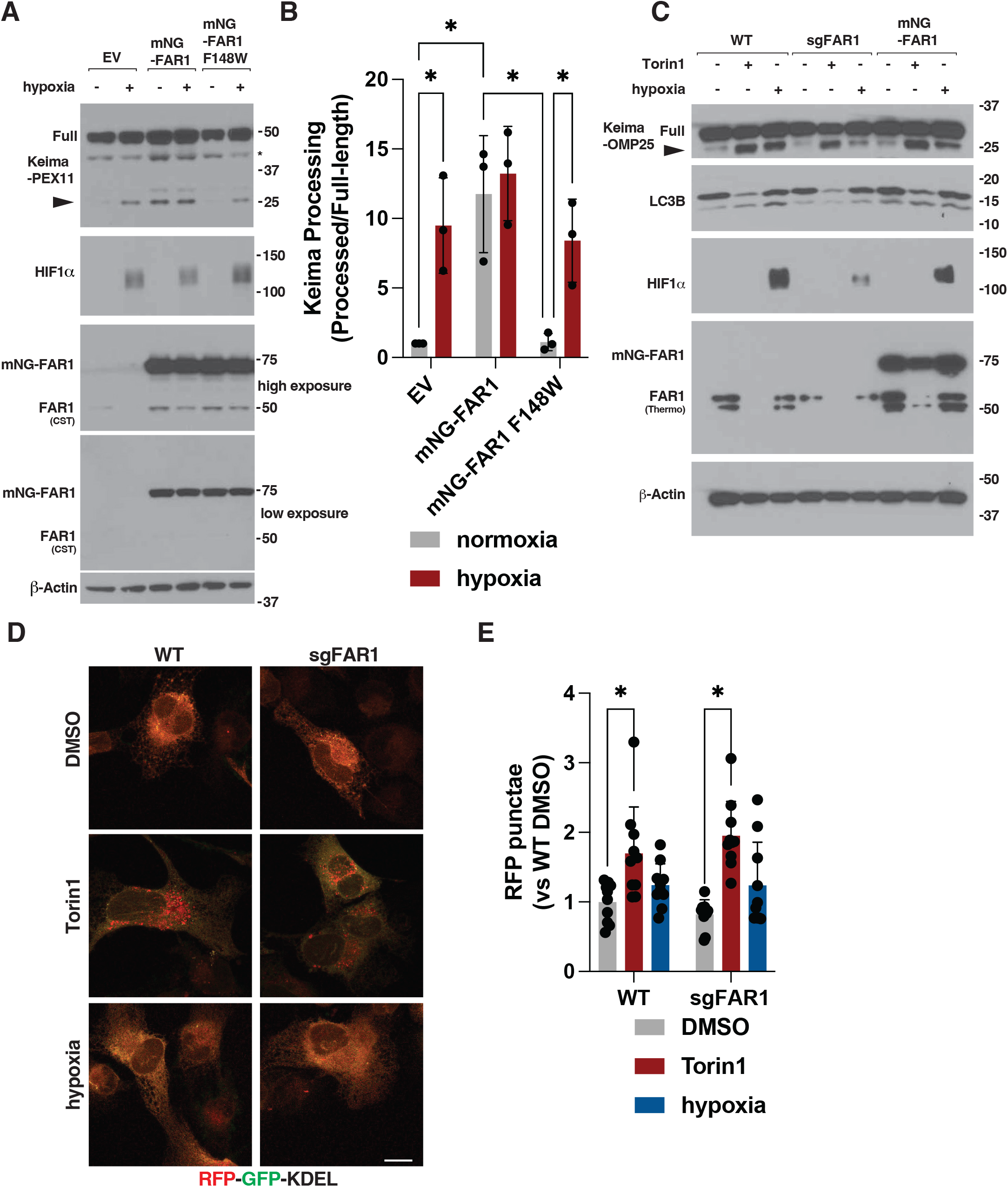
Analysis of hypoxia-induced pexophagy, mitophagy, and ER-phagy. **(A)** Immunoblot analysis of wild-type AC16 cardiomyocytes stably expressing Keima-PEX11 and mNG-FAR1, mNG-FAR1 F148W, or empty vector (EV). Where indicated, cells were grown under 0.1% O2 (hypoxia) or normoxia for 18 hours prior to cell lysis. Full indicates full-length Keima-fusion protein and arrow indicates liberated Keima protein. Far1 immunoblots indicate which Far1 antibody was used to detect Far1. **(B)** Densitometry analysis of Keima-processing assay as shown in wild-type human AC16 cardiomyocytes stably expressing Keima-PEX11 and mNG-FAR1, mNG-FAR1 F148W, or empty vector (EV). **(C)** Immunoblot analysis of wild-type (WT) and sgFAR1 human AC16 cardiomyocytes stably expressing Keima-OMP25. Where indicated, wild-type cardiomyocytes were stably expressing empty vector or mNeonGreen-Far1 (mNG-FAR1). Cells were treated with 250 nM Torin1, grown at 0.1% O2 (hypoxia), or treated with DMSO and grown at normoxia for 18 hours prior to cell lysis. Far1 immunoblot indicates which Far1 antibody was used to detect Far1. **(D-E)** Confocal microscopy (D) and RFP imaging analysis (E) of wild-type (WT) and sgFAR1 human AC16 cardiomyocytes stably expressing RFP-GFP-KDEL. Where indicated, cells were treated with 250 nM Torin1, grown at 0.1% O_2_ (hypoxia), or grown at normoxia and treated with DMSO for 18 hours prior to imaging. N=10. *indicates p<0.05. Scale bar indicates 10 um.

**Supplementary Figure 14.**
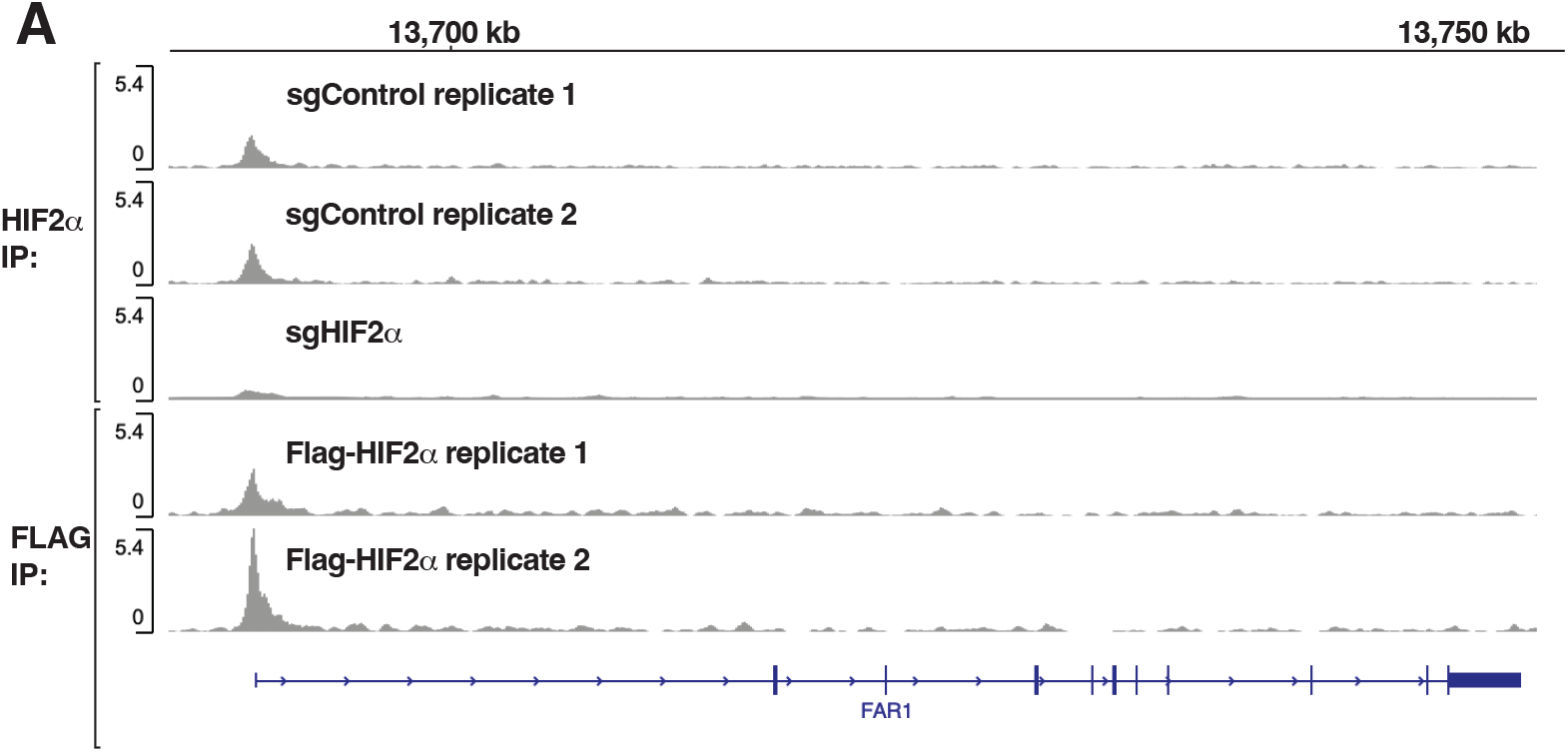
FAR1 is a direct HIF target. **(A)** FAR1 ChIP-Seq analysis of publicly available GSE253325 data. Representative Anti-HIF2α ChIP-seq tracks in 786-O cells that underwent CRISPR-based gene editing with a HIF2α sgRNA or a control sgRNA (sgCtrl) and representative Anti-FLAG ChIP-seq tracks in 786-O cells in which a FLAG-HA epitope tag was inserted at the 5’ end of the endogenous *EPAS1* open reading frame by CRISPR-HDR (FLAG-HIF2α).

**Supplementary Figure 15.**
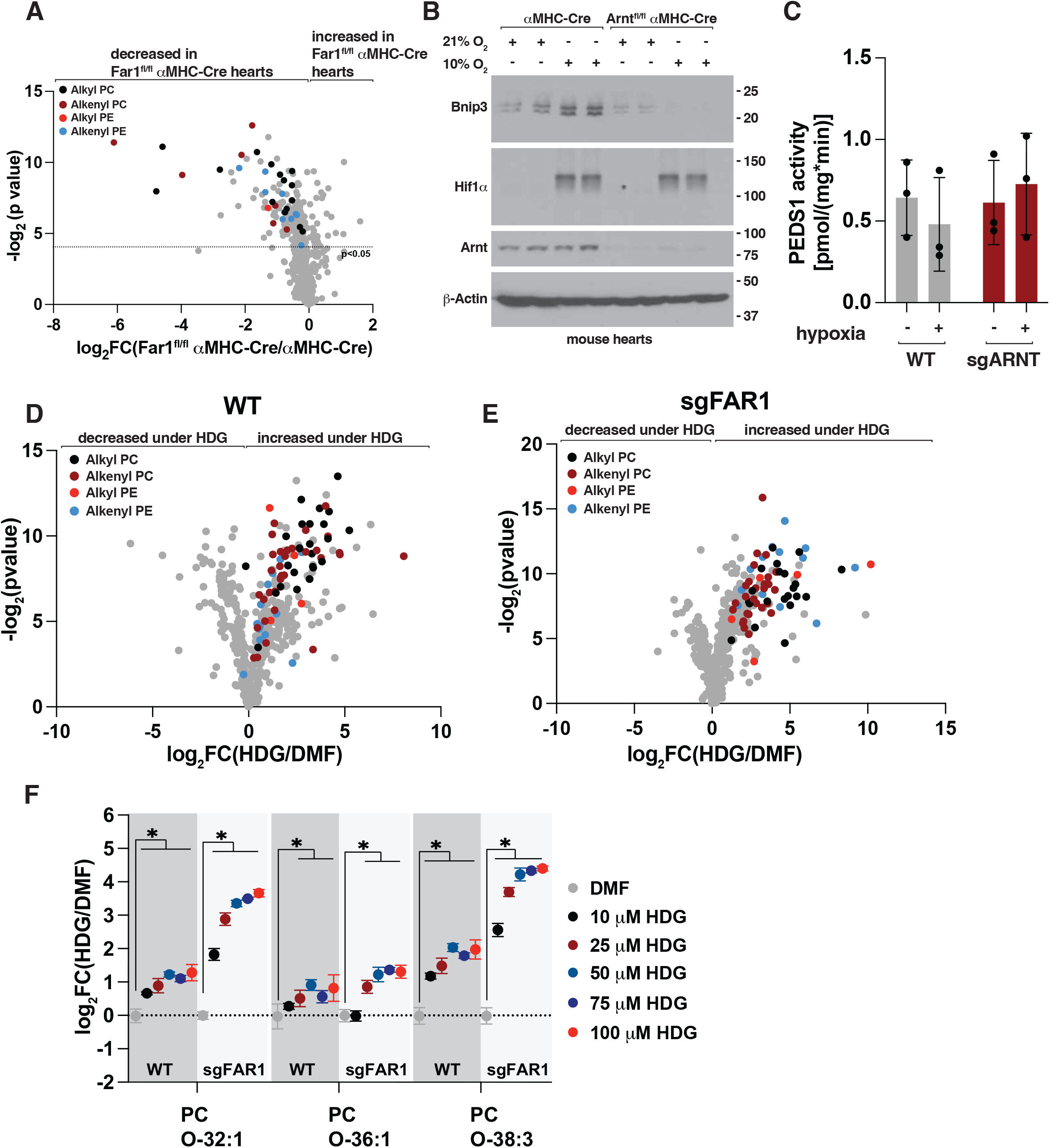
Ether lipids abundance under hypoxia and HDG. **(A)** Volcano plots comparing lipid intensities in αMHC-Cre and Far1^fl/fl^ αMHC-Cre mouse hearts. **(B)** Immunoblot analysis of Arnt^+/+^-αMHC-Cre and Arnt^fl/fl^-αMHC-Cre mouse hearts. Where indicated, mice were housed at 10% O_2_ (hypoxia) or 21% O_2_ (atmospheric oxygen) for 7 days prior to euthanasia. **(C)** PEDS1 activity [pmol/(mg*min)] in wild-type and sgARNT human AC16 cardiomyocytes. Where indicated, cells were grown at 0.1% O_2_ (hypoxia), or normoxia for 18 hours prior to cell harvesting. N=3. **(D-E)** Volcano plots comparing lipid intensities in wild-type (D) and sgFAR1 (E) human AC16 cardiomyocytes treated with 1-O-hexadecyl-*sn*-glycerol (100 uM, HDG) or DMF for 24 hours prior to lipid extraction. Black dots indicate alkyl (ether) phosphatidylcholine (PC), dark red dots indicate alkenyl PC (plasmalogens) species, red dots indicate alkyl (ether) phosphotidylethanolamine (PE), and light blue dots indicate alkenyl PE (plasmalogens) species. **(F)** Lipidomic profiling of alkyl PC species in wild-type and sgFAR1 human AC16 cardiomyocytes. Where indicated, cells were treated with 1-O-hexadecyl-sn-glycerol (HDG) at the indicated dose (uM) or DMF for 8 hours prior to lipid extraction. N=3, *P<0.05.

**Supplementary Figure 16.**
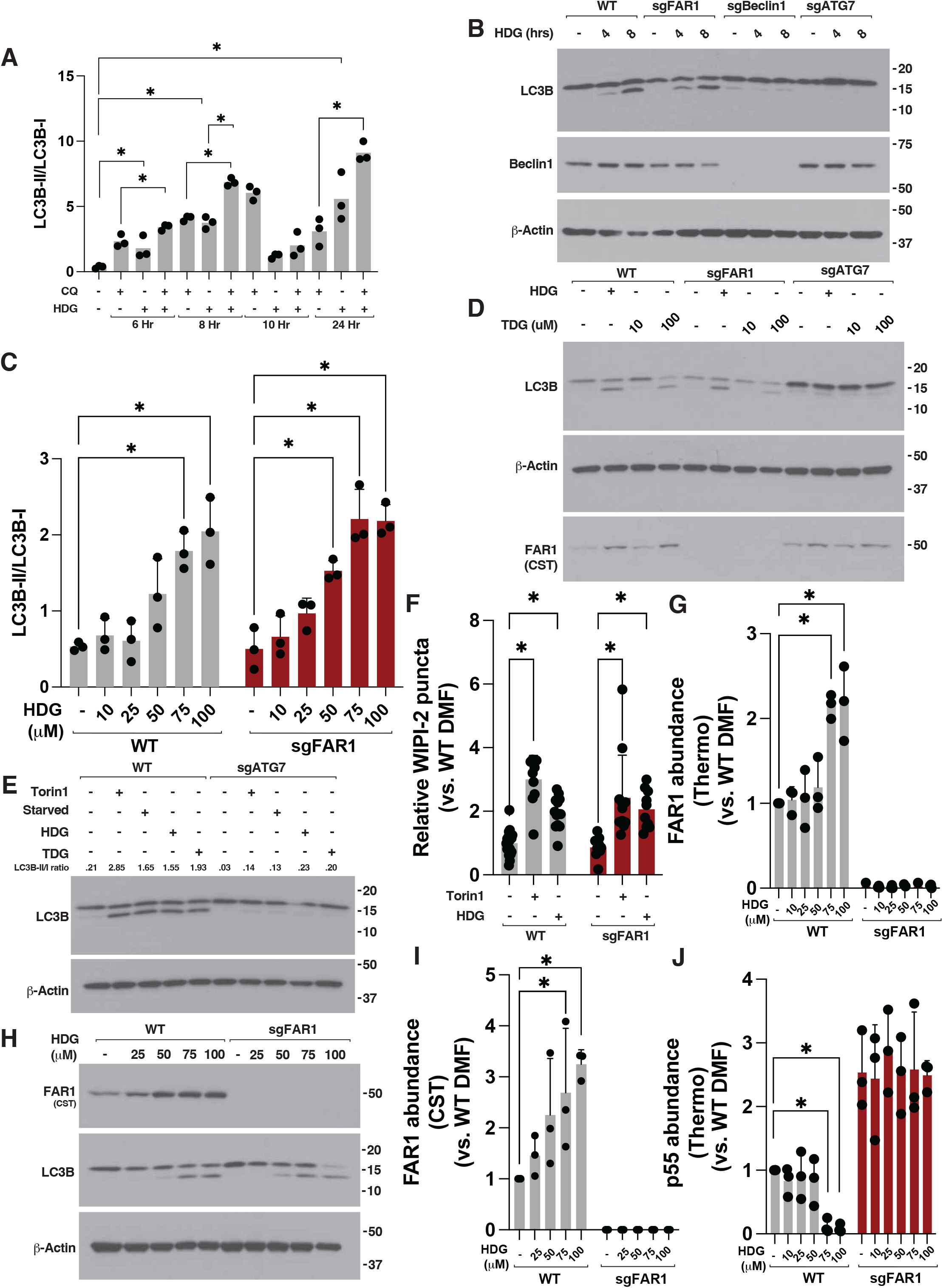
Ether lipid precursors activate autophagy. **(A)** Densitometry analysis of LC3B-II/LC3B-I ratio from immunoblot assays. Where indicated, cell lysates were prepared from wild-type human AC16 cells treated with 100 uM 1-O-hexadecyl-*sn*-glycerol (HDG), 30 uM Chloroquine (CQ), or DMF for indicated time (hours, hrs). N=3, *indicates P<0.05. **(B)** Immunoblot analysis of wild-type (WT), sgFAR1, sgBeclin1, and sgATG7 human AC16 cardiomyocytes. Where indicated, cells were treated with 100 uM 1-O-hexadecyl-*sn*-glycerol (HDG) or DMF for indicated time (hours, hrs) prior to cell lysis. **(C)** Densitometry analysis of LC3B-II/LC3B-I ratio from immunoblot assays. Where indicated, cell lysates were prepared from wild-type human or sgFAR1 AC16 cells treated with indicated concentration of 1-O-hexadecyl-*sn*-glycerol (HDG) or DMF. N=3, *indicates P<0.05. **(D)** Immunoblot analysis of wild-type (WT), sgFAR1, and sgATG7 human AC16 cardiomyocytes. Where indicated, cells were treated with 100 uM 1-O-hexadecyl-*sn*-glycerol (HDG), indicated dose of 1-O-tetradecyl glycerol (TDG), or DMF for 8 hours prior to cell lysis. Far1 immunoblot indicates which Far1 antibody was used to detect Far1. **(E)** Immunoblot analysis of wild-type (WT) and sgATG7 human AC16 cardiomyocytes. Where indicated, cells were treated with 250 nM Torin1, starved of nutrients, treated with 100 uM 1-O-hexadecyl-*sn*-glycerol (HDG), treated with 100 uM 1-O-tetradecyl glycerol (TDG), or DMF for 8 hours prior to cell lysis. **(F)** Confocal microscopy and image analysis of WIPI-2 puncta in wild-type and sgFAR1 human AC16 cells. Where indicated, cells were treated with 250 nM Torin1, 100 uM 1-O-hexadecyl-*sn*-glycerol (HDG), or DMF for 8 hours prior to cell fixation. N>10, *indicates P<0.05. **(G)** Densitometry analysis of FAR1 abundance using the Thermo PA5-53585 antibody from immunoblot assays. Where indicated, cell lysates were prepared from wild-type and sgFAR1 human AC16 cells treated with indicated dose of 1-O-hexadecyl-*sn*-glycerol (HDG) or DMF for 24 hours. N=3, *indicates P<0.05. **(H)** Immunoblot analysis of wild-type (WT) and sgFAR1 human AC16 cardiomyocytes. Where indicated, cells were treated with indicated dose of 1-O-hexadecyl-*sn*-glycerol (HDG) or DMF for 24 hours. FAR1 immunoblot indicates which FAR1 antibody was used to detect FAR1. **(I)** Densitometry analysis of FAR1 abundance using the CST FAR1 antibody from immunoblot assays. Where indicated, cell lysates were prepared from wild-type and sgFAR1 human AC16 cells treated with indicated dose of 1-O-hexadecyl-*sn*-glycerol (HDG) or DMF for 24 hours. N=3, *indicates P<0.05. **(J)** Densitometry analysis of 55 kDa band (p55) abundance detected using FAR1 Thermo PA5-53585 antibody from immunoblot assays. Where indicated, cell lysates were prepared from wild-type and sgFAR1 human AC16 cells treated with indicated dose of 1-O-hexadecyl-*sn*-glycerol (HDG) or DMF for 24 hours. N=3, *indicates P<0.05.

